# xMWAS: an R package for data-driven integration and differential network analysis

**DOI:** 10.1101/122432

**Authors:** Karan Uppal, Young-Mi Go, Dean P. Jones

## Abstract

**Summary:** Integrative omics is a central component of most systems biology studies. Computational methods are required for extracting meaningful relationships across different omics layers. Various tools have been developed to facilitate integration of paired heterogenous omics data; however most existing tools allow integration of only two omics datasets. Furthermore, existing data integration tools do not incorporate additional steps of identifying sub-networks or communities of highly connected entities and evaluating the topology of the integrative network under different conditions. Here we present xMWAS, an R package for data integration, network visualization, clustering, differential network analysis of data from biochemical and phenotypic assays, and two or more omics platforms.

**Availability:** https://sourceforge.net/projects/xmwas/

## 1 Introduction

Technological advances have led to a major paradigm shift where multi-assay molecular profiling of biological samples is increasingly being used to understand molecular mechanisms for diseases and host responses to environmental exposures (Hawkins 2010, Cancer Genome Atlas Network 2008). Most cellular processes in a biological system are dependent on complex molecular interactions (Barabasi 2011). Integrative omics allows researchers to address such complexity and answer challenging biological questions, such as function of genetic variants and unknown metabolites, mechanisms of gene regulation, signaling and metabolic pathway responses to infection and toxicity (Hawkins 2010, Chandler 2016, Uppal 2016).

Numerous data-driven/unsupervised and knowledge-based tools allow integration of data from different omics technologies and other molecular assays (Wanichthanarak 2015, Meng 2016). Most existing data integration tools allow integration of only two datasets and do not allow identification of community structure and evaluation of network changes between different conditions. Community detection reveals topological modules comprised of functionally related biomolecules (Barabasi 2011, Yang 2016). Differential network analysis (DiNA) allows characterization of nodes that undergo changes in topological characteristics between different conditions, e.g. healthy vs disease (Lichtblau 2016).

To advance these capabilities, we present, xMWAS, an R package that provides an automated workflow for integrative analysis of more than two datasets, differential network analysis, and community detection to improve our understanding of complex molecular interactions and disease mechanisms.

**Fig. 1.**
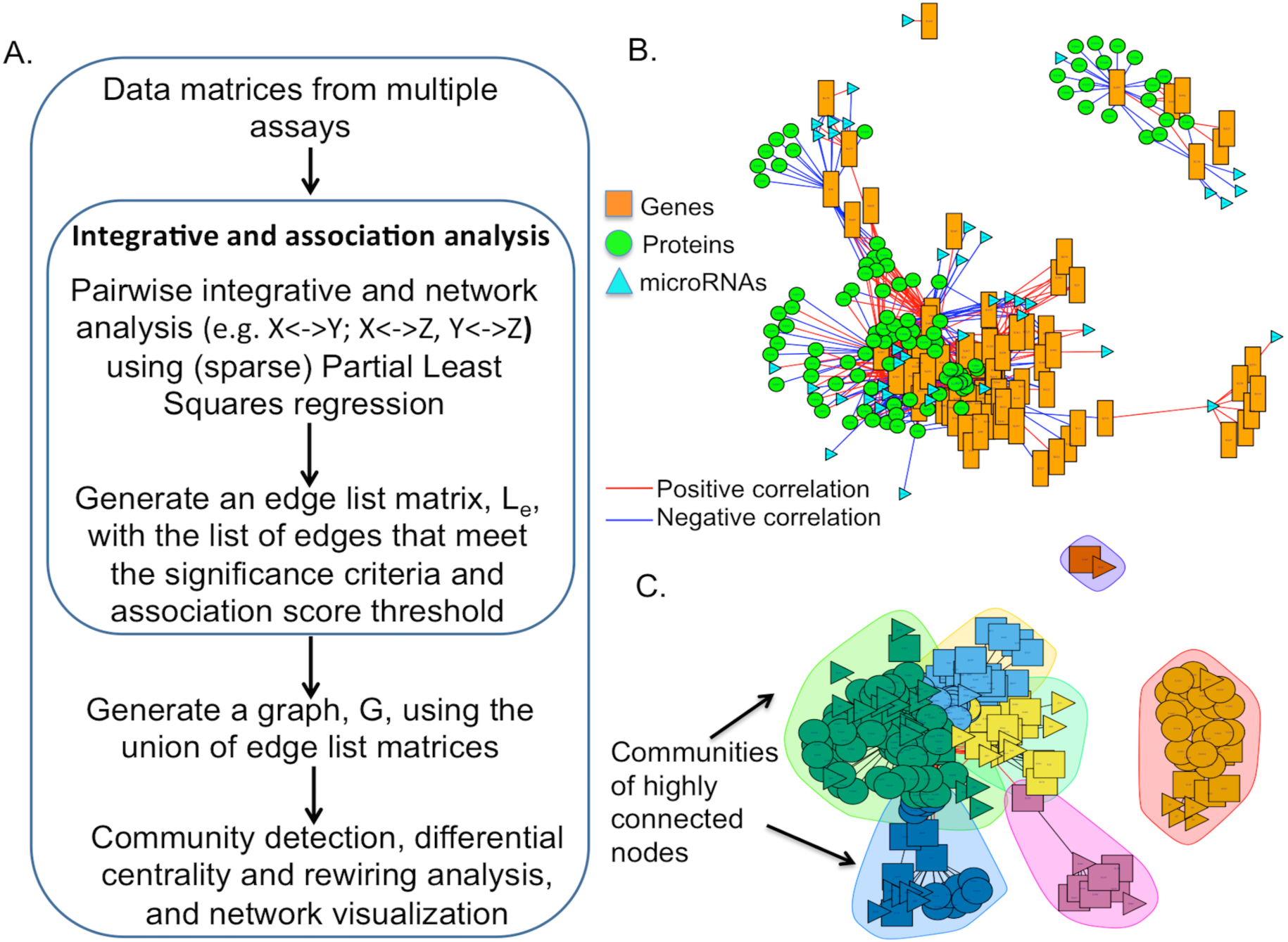
xMWAS workflow. A. Pairwise data integration and association analysis is performed using (sparse) partial least squares regression. The edge list matrices of significant associations from each pairwise comparison are merged to generate a global association network. Community detection is performed using the multilevel algorithm. B. Illustration of association network for the NCI60 dataset (Meng 2016): microRNAs (triangle), protein (circle) and transcripts (rectangle). C. Identification of communities in the NCI60 integrated network using the multilevel community detection algorithm.

## 2 Implementation

xMWAS provides an automated framework for integrative and differential network analysis. Figure 1A provides an overview of different stages of xMWAS. In stage one, xMWAS uses dimension reduction techniques such as Partial Least Squares (PLS), sparse Partial Least Squares (sPLS), and multilevel sparse Partial Least Squares (msPLS; for repeated measures) regression for pairwise integrative and association analysis between data matrices (Le Cao 2009, Liquet 2012, Gonzalez 2012). sPLS and msPLS methods perform simultaneous data integration and variable selection using a LASSO penalty for the loading vectors, which reduces the complexity of the networks (Liquet 2012). R package plsgenomics is used to determine the optimal number of latent components. The *network()* function in the *mixOmics* package is used to generate the association matrix, A_XY_, between matrices X and Y (Le Cao 2009, Gonzalez 2012). Student’s t-test is used to evaluate the statistical significance of association scores. Only the associations that satisfy the user-defined thresholds, e.g. |association score|>0.7 and *p*-value<0.01, are used for downstream analysis. The resulting graph, G_i_=(V,E), where V is a set of nodes and E is a set of edges, is used to generate an edge list matrix, L_i_., such that each row in L_i_ corresponds to an edge between nodes X_p_ and Y_q_. The same process is repeated for generating edge list matrices from all pairwise association analyses between datasets, e.g. L_i_=cor(X,Y); L_j_=cor(Y,Z), and L_k_=cor(Y,Z).

In stage two, the union of the individual edge list matrices from pairwise integrative analysis of the n datasets is used to generate a combined edge list matrix, L_e_ = 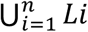. Matrix L_e_ is used to generate the integrative network graph, G=(V,E), where V corresponds to nodes and E corresponds to edges or connections between the nodes, representing positive or negative associations between multiple datasets (Figure 1B). Network graphics are generated using the *igraph* package in R.

In stage three, the multilevel community detection algorithm (Blondel 2008) is used to identify communities of nodes that are tightly connected with each other, but sparsely connected with the rest of the network (Figure 1C). Comparative studies for community detection algorithms show that the multilevel algorithm is suitable for both small and large networks with varying connectivity patterns (Yang 2016). The quality of the community structure is evaluated using the network modularity measure (Newman 2006).

Differential network analysis is performed using the differential betweenness centrality and differential eigenvector centrality methods to identify nodes that undergo changes in their topological characteristics (Odibat 2012, Lichtblau 2016). Additional description about the software input and output is provided in Supplementary Section S1.

## 3 Example

We tested xMWAS in a three-way integrative analysis using cytokine, transcriptome, and metabolome datasets from a recently published study to examine H1N1 influenza virus infection-altered metabolic response in mouse lung (Chandler 2016). For comparisons, we used data from all samples (Supplementary Figure S1A), only control samples (Supplementary Figure S1B), and only H1N1 influenza samples (Supplementary Figure S1C). Supplementary Section S2 shows that the various stages of xMWAS capture biologically meaningful information and provide deeper insights into the underlying biology, which cannot be obtained by analyzing and exploring the different layers individually.

## 4 Conclusion

xMWAS provides a platform-independent framework for integrative network analysis of two or more datasets, identification of modules of functionally related biomolecules, and differential network analysis. The results show that xMWAS can improve our understanding of disease pathophysiology and complex molecular interactions across various functional levels.

## Acknowledgements

The authors acknowledge members of the Clinical Biomarkers Laboratory, Emory University for testing and suggesting improvements to the software.

## Funding

This project was funded by National Institutes of Health grants, ES025632, ES023485, ES019776, OD018006, HL095479, EY022618. The project was also funded in part by federal funds from the US National Institute of Allergy and Infectious Diseases, National Institutes of Health, Department of Health and Human Services under contract # HHSN272201200031C.

## Conflict of Interest

none declared.

## Supplementary Material

### Supplementary Section S1. Description of input and output for xMWAS

The main input parameters include file paths or data matrices of input datasets, file with phenotypic labels (e.g. case, control), integration method (“pls”, “spls”), threshhold for association score, and statistical significance. The software is designed to work with data from different sources and does not utilize any knowledgebase information for integration. For metabolomics studies, the software is designed to work with both targeted and untargeted datasets and does not require metabolite identification prior to integration. The output includes PDF of integrative network with cluster assignments, text files for association matrices and cluster assignments for each node and their centrality scores, and a GML format file that can be used for visualization with tools such as Cytoscape (Shannon 2003).

**Figure S1.**
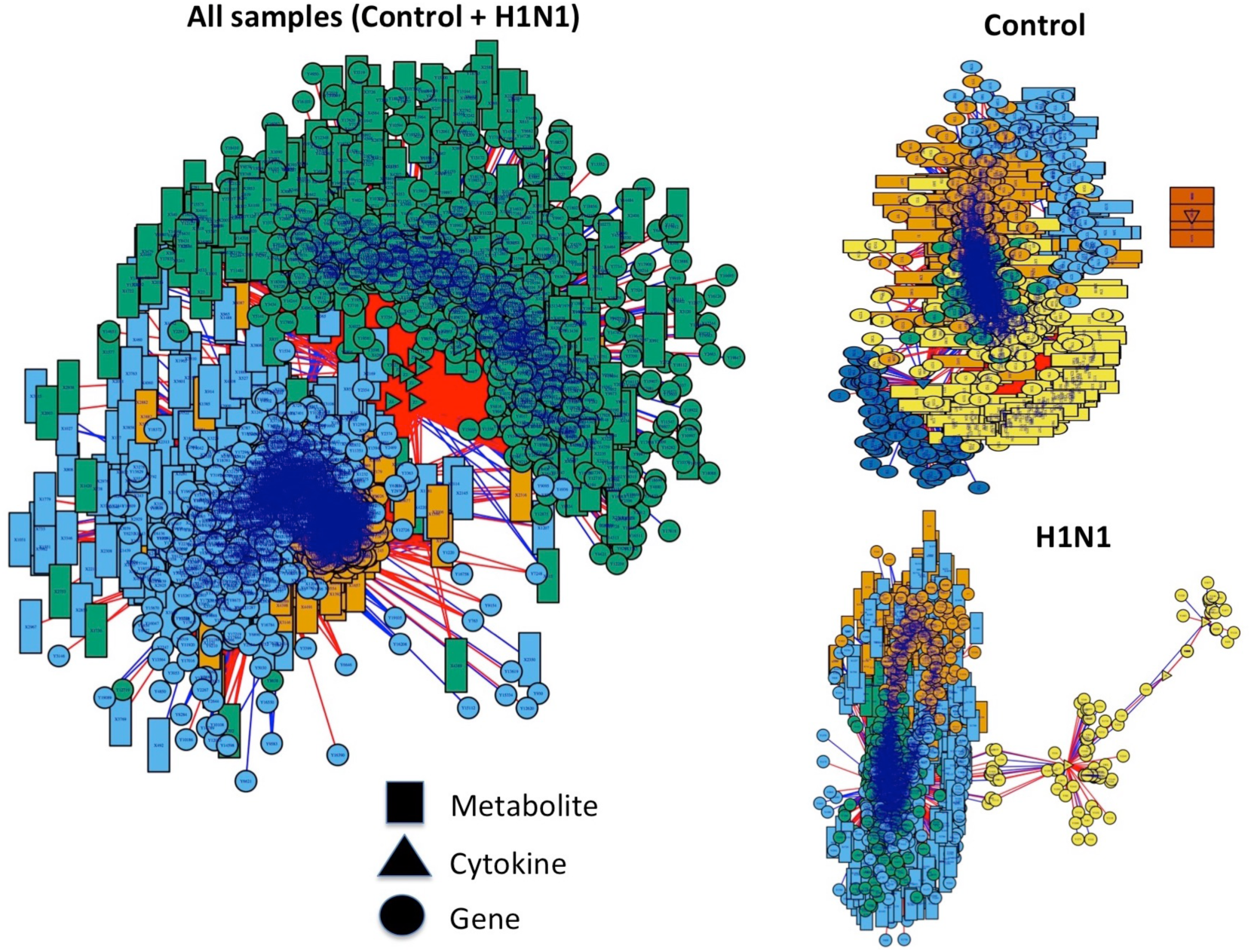
Integrative network analysis of cytokine, metabolome and transcriptome datasets from a study of H1N1 virus infection of mice. A. All samples; B. Only control samples; C. Only H1N1 infected samples. The results show that xMWAS analysis of the complex omics arrays allows identification and visualization of metabolite and transcript correlations with cytokines. The analysis further discriminates between correlations observed in controls from those observed in H1N1 virus infected mice and allows identification of nodes that undergo network changes. Details are provided in Supplementary Section S2.

### Supplementary Section S2. Evaluation results using data from the H1N1 virus infection of mice

Pathway analysis of metabolic features (*m/z* and retention time) and genes that were significantly associated with cytokines at *p*<0.05 across all samples was performed using Mummichog and MetaCore, respectively. Pathways related to immune response, lipid metabolism, amino acids metabolism, nitrogen metabolism, drug metabolism, vitamin metabolism, cytoskeleton remodeling, cell signaling, and energy metabolism were significantly enriched (Supplementary Tables S1 and S2). Previous studies have shown the role of these pathways in regulation of immune response and inflammation (Miyake 2000, Fortin 2009, Sadik 2012, Yin 2014, Chandler 2016)

Differential centrality analysis between control vs H1N1 samples showed dramatic change in the centrality of cytokines as well as genes and metabolites involved in immune response and defense mechanism (Supplementary Tables S3-6). Cytokines are known to be involved in the recruitment of the inflammatory cells and influence the adaptive immune response during influenza (Vareille 2011, Liu 2016, Chandler 2016).

Multilevel community detection identified four clusters in the integrative network generated using only H1N1 samples (Supplementary Figure S1C). Cluster 2 (blue), which comprised of four cytokines, 830 genes, and 437 metabolic features, was evaluated for significantly enriched pathways and biological processes. Pathway analysis of the metabolic features in this cluster showed enrichment of similar pathways as in the cytokine x metabolome association analysis across all samples described above (Supplementary Table S7). Pathway and process enrichment analysis using MetaCore showed enrichment of genes involved in immune responses, protein folding, signaling mechanisms, and various biological processes related to cilium movement, organization, assembly and organization (Supplementary Tables S8-S10). Previous studies have shown that the airway epithelium acts as a first layer of defense mechanism against respiratory viruses and that the respiratory viruses can lead to ciliary impairment (Vareille 2011).

**Supplementary Table S1.**
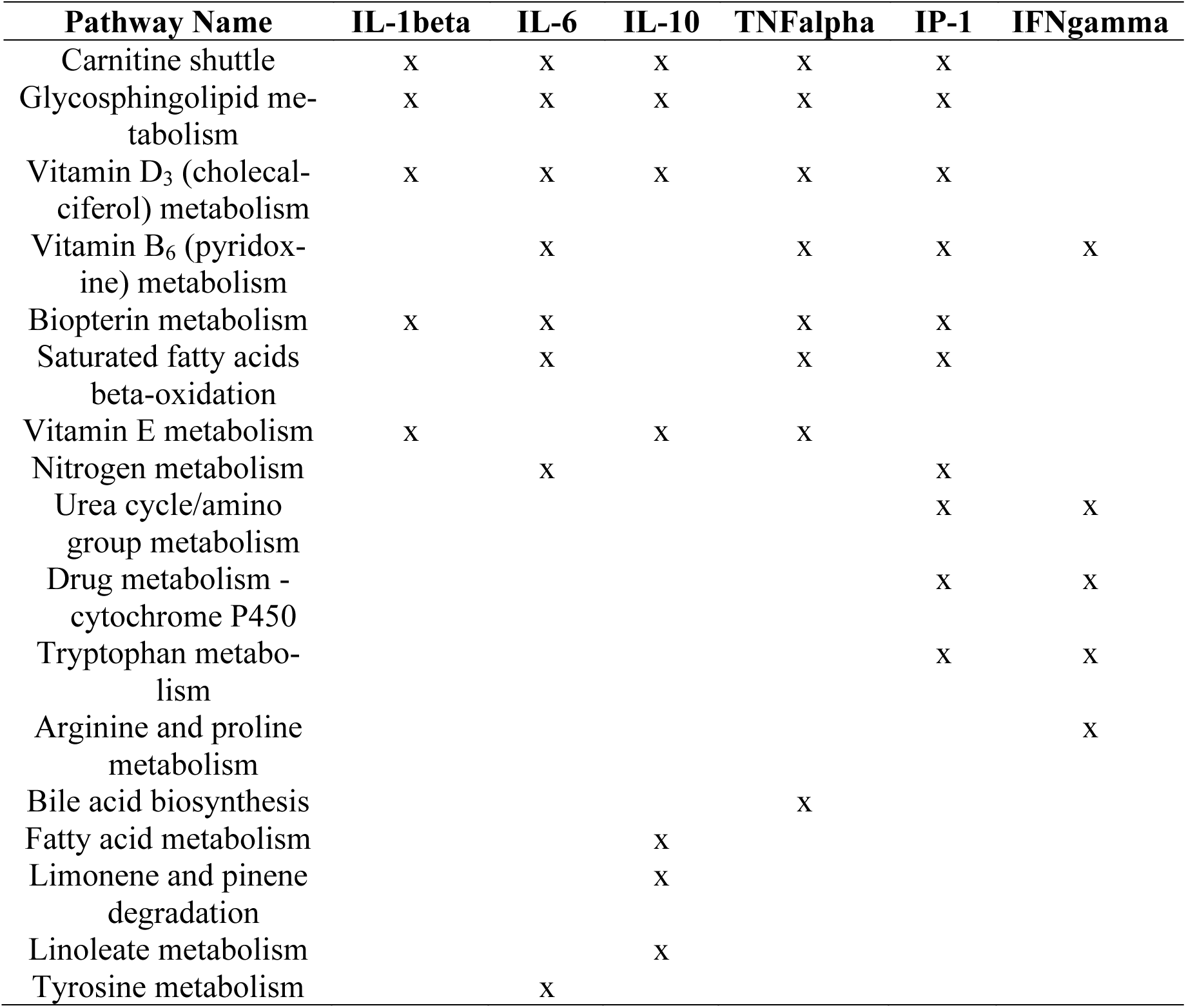
Pathway analysis of metabolic features (*m/z* and retention time) that were found to be significantly associated with the six cytokines at *p*<0.05 using Mummichog

**Supplementary Table S2.** Pathway analysis of genes that were found to be significantly associated with the six cytokines at *p*<0.05 using MetaCore

| Pathway Name | IL-1beta | IL-6 | IL-10 | TNFalpha | IP-1 | IFNgamma |
| --- | --- | --- | --- | --- | --- | --- |
| Immune response_Alternative complement pathway | x | x | x | x | x | x |
| GTP metabolism | x | x | x | x | x |  |
| Cytoskeleton remodeling_TGF, WNT and cytoskeletal remodeling | x | x |  | x | x | x |
| Cytoskeleton remodeling_Cytoskeleton remodeling | x | x |  | x | x | x |
| Alternative complement cascade disruption in age-related macular degeneration | x | x |  | x | x | x |
| ATP metabolism | x | x |  | x | x |  |
| Immune response_Lectin induced complement pathway |  | x | x |  | x | x |
| Regulation of lipid metabolism_Regulation of lipid metabolism by niacin and isoprenaline | x | x |  | x |  |  |
| Development_Oligodendrocyte differentiation from adult stem cells |  | x |  |  | x | x |
| Cell adhesion_Chemokines and adhesion | x |  |  | x |  |  |
| Expression targets of Tissue factor signaling in cancer | x |  |  | x |  |  |
| Cell adhesion_ECM remodeling |  | x |  |  |  | x |
| Immune response_Classical complement pathway |  |  | x |  |  | x |
| Development_Regulation of epithelial-to-mesenchymal transition (EMT) |  |  |  |  | x | x |
| Protein folding and maturation_Insulin processing |  |  | x |  |  |  |
| Glucocorticoid-induced elevation of intraocular pressure as glaucoma risk factor |  |  |  |  |  | x |
| Inhibition of neutrophil migration by proresolving lipid mediators in COPD |  |  |  |  |  | x |

**Supplementary Table S3.** Eigenvector centrality measures for cytokines with delta centrality (control vs H1N1) greater than 0.3

| **Name** | **All samples** | **Control** | **H1N1** | **Delta**<br>**(Control vs H1N1)** |
| --- | --- | --- | --- | --- |
| IL-10 | 0.61 | 0.00 | 1.00 | 1.00 |
| IL-1beta | 0.66 | 0.00 | 0.98 | 0.98 |
| TNF-alpha | 0.66 | 0.00 | 0.88 | 0.88 |
| IL-6 | 0.34 | 0.50 | 0.00 | 0.50 |

**Supplementary Table S4.** Pathway analysis of metabolic features with delta centrality (control vs H1N1) greater than 0.3 using Mummichog

| **Pathways** | **Overlap_size** | **Pathway_size** | ***p*** |
| --- | --- | --- | --- |
| Carnitine shuttle | 16 | 18 | 0.00065 |
| Vitamin D3 (cholecalciferol) metabolism | 10 | 10 | 0.00065 |
| Phytanic acid peroxisomal oxidation | 4 | 4 | 0.00115 |
| De novo fatty acid biosynthesis | 8 | 14 | 0.00134 |
| Leukotriene metabolism | 11 | 23 | 0.00174 |
| Valine, leucine and isoleucine degradation | 10 | 22 | 0.00278 |
| Fatty acid activation | 6 | 11 | 0.00288 |
| Fatty Acid Metabolism | 6 | 11 | 0.00288 |
| Vitamin E metabolism | 7 | 15 | 0.00474 |
| Squalene and cholesterol biosynthesis | 7 | 15 | 0.00474 |
| Pyruvate Metabolism | 7 | 15 | 0.00474 |
| Omega-6 fatty acid metabolism | 2 | 2 | 0.01496 |
| Benzoate degradation via CoA ligation | 2 | 2 | 0.01496 |
| Fatty acid oxidation, peroxisome | 2 | 2 | 0.01496 |
| Di-unsaturated fatty acid beta-oxidation | 2 | 2 | 0.01496 |
| Prostaglandin formation from arachidonate | 14 | 40 | 0.01604 |
| Biopterin metabolism | 5 | 12 | 0.01882 |
| Bile acid biosynthesis | 16 | 47 | 0.01928 |
| Glycosphingolipid metabolism | 8 | 22 | 0.02174 |
| Vitamin B5 - CoA biosynthesis from pantothenate | 3 | 6 | 0.02583 |
| Tryptophan metabolism | 16 | 49 | 0.03124 |
| Saturated fatty acids beta-oxidation | 2 | 3 | 0.03833 |
| Glyoxylate and Dicarboxylate Metabolism | 3 | 7 | 0.04628 |

**Supplementary Table S5.**
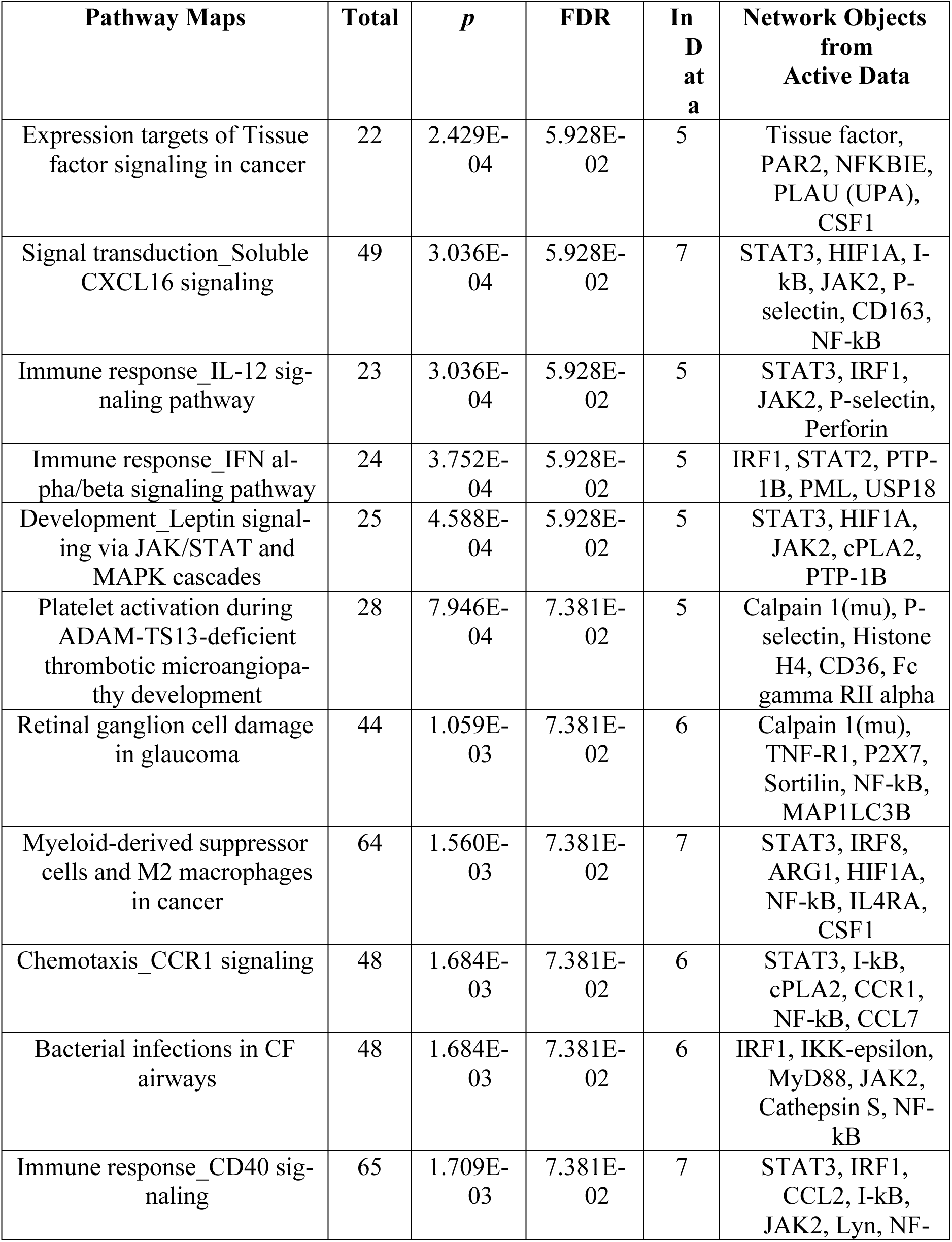
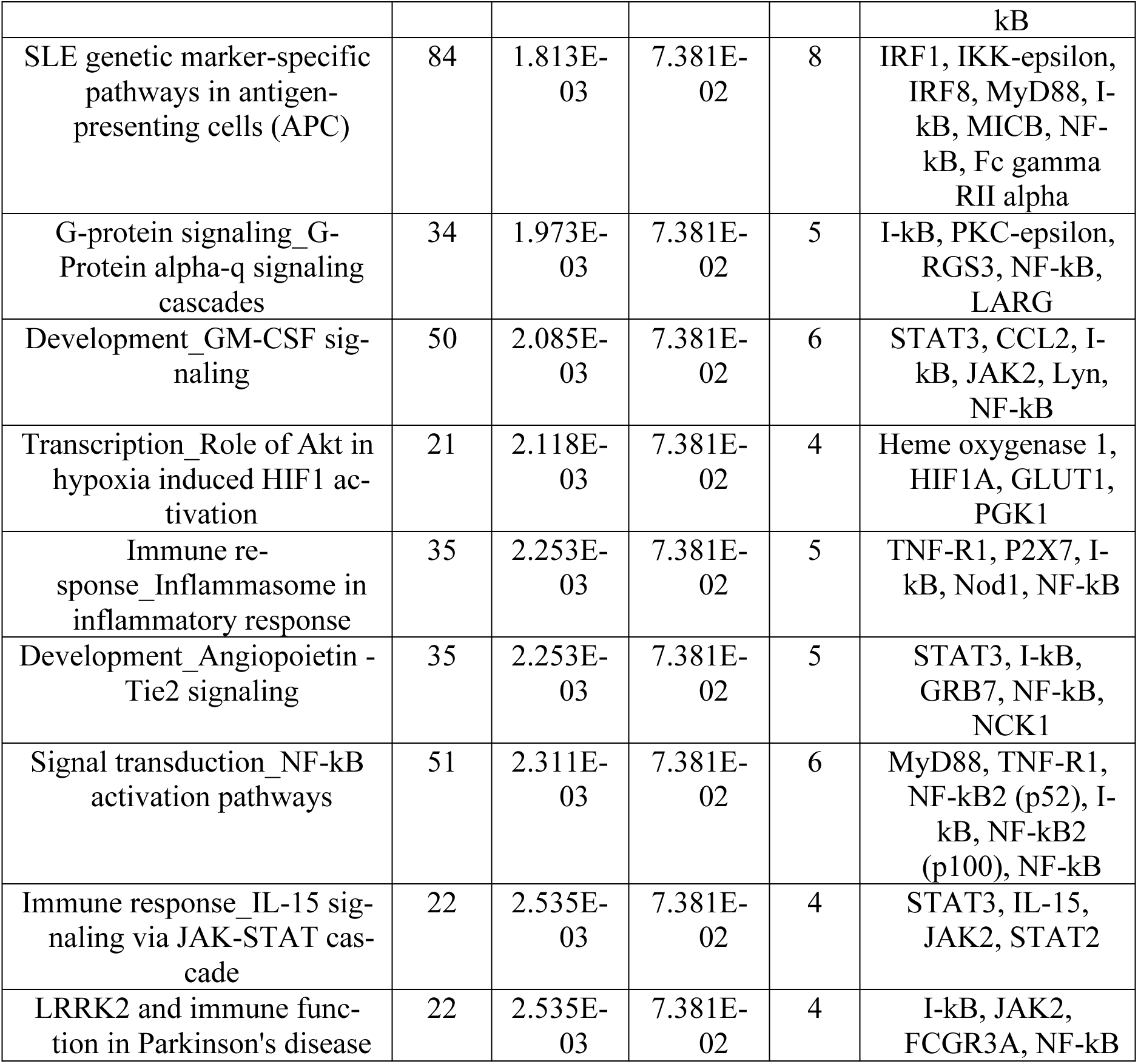
Pathway analysis of genes with delta centrality (control vs H1N1) greater than 0.3 using MetaCore. Only the top 20 significant pathways at FDR<0.1 are shown here.

**Supplementary Table S6.**
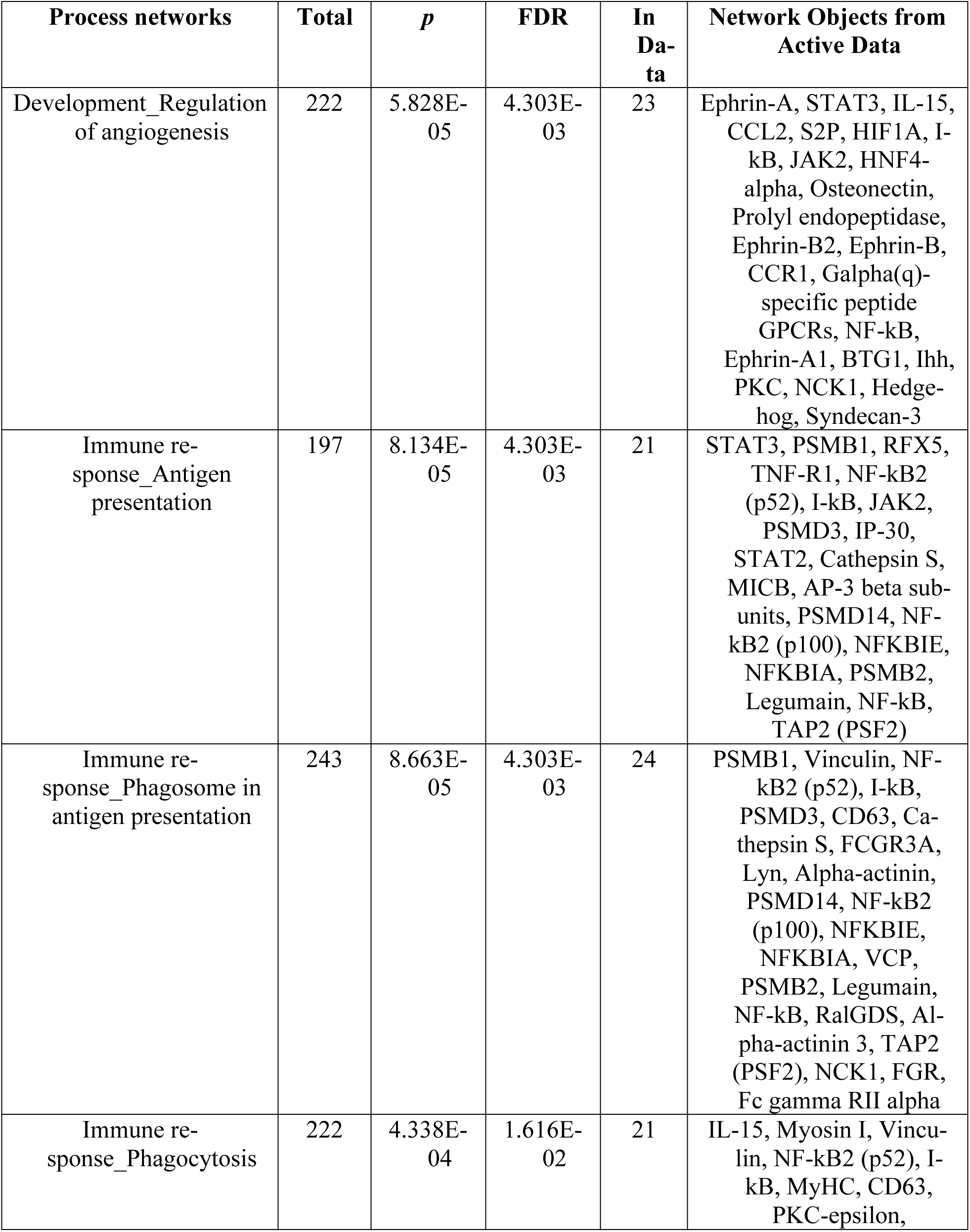
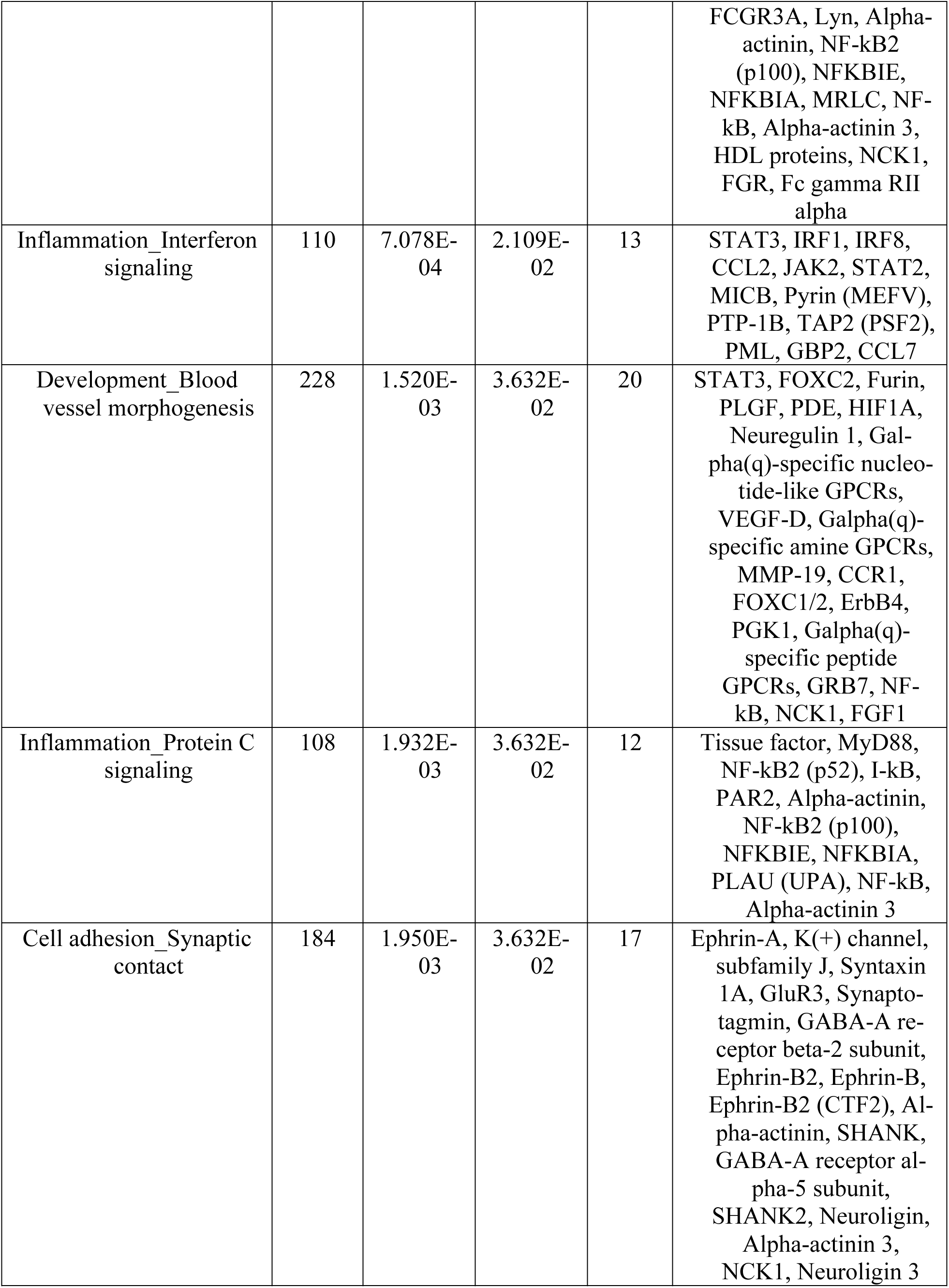
Process networks enrichment analysis of genes with delta centrality (control vs H1N1) greater than 0.3 using MetaCore.

**Supplementary Table S7.** Pathway analysis of the 437 metabolic features in cluster 2 in the H1N1 integrative network using Mummichog

| **Pathways** | **Overlap_size** | **Pathway_size** | ***p*** |
| --- | --- | --- | --- |
| Carnitine shuttle | 9 | 18 | 0.00036 |
| Vitamin D<sub>3</sub> (cholecalciferol) metabolism | 6 | 10 | 0.00037 |
| Linoleate metabolism | 7 | 21 | 0.00050 |
| Vitamin E metabolism | 5 | 15 | 0.00081 |
| Glycosphingolipid metabolism | 6 | 22 | 0.00100 |
| Bile acid biosynthesis | 10 | 47 | 0.00108 |
| Saturated fatty acids beta-oxidation | 2 | 3 | 0.00305 |
| Pyruvate Metabolism | 3 | 15 | 0.01784 |
| C21-steroid hormone biosynthesis and metabolism | 9 | 64 | 0.02414 |

**Supplementary Table S8.**
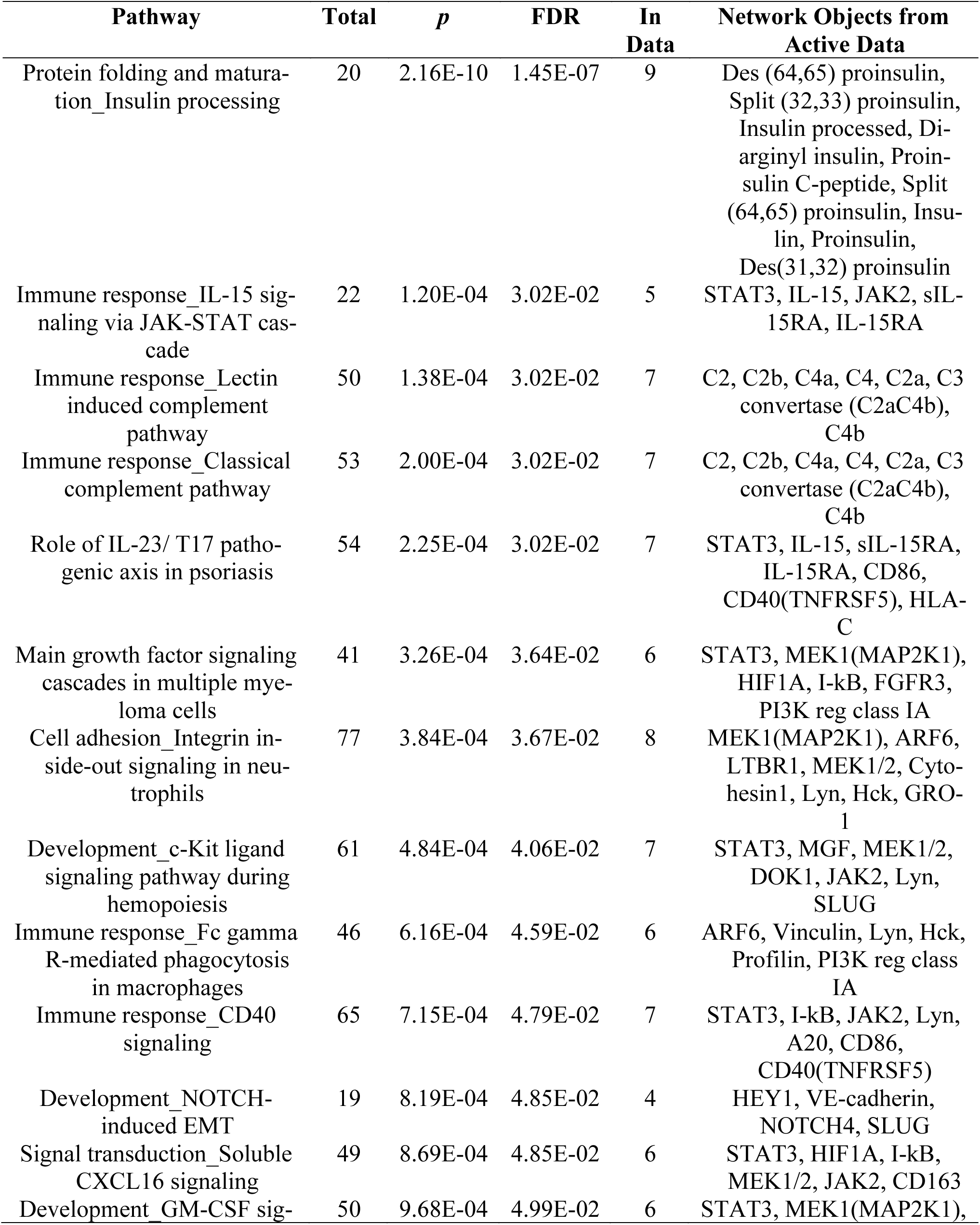
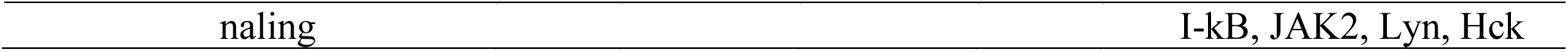
Pathway analysis of the 830 genes in cluster 2 in the H1N1 integrative network using MetaCore

| Pathway | Total | *p* | FDR | In Data | Network Objects from Active Data |
| --- | --- | --- | --- | --- | --- |
| Protein folding and maturation_Insulin processing | 20 | 2.16E-10 | 1.45E-07 | 9 | Des (64,65) proinsulin, Split (32,33) proinsulin, Insulin processed, Di-arginyl insulin, Proinsulin C-peptide, Split (64,65) proinsulin, Insulin, Proinsulin, Des(31,32) proinsulin |
| Immune response_IL-15 signaling via JAK-STAT cascade | 22 | 1.20E-04 | 3.02E-02 | 5 | STAT3, IL-15, JAK2, sIL-15RA, IL-15RA |
| Immune response_Lectin induced complement pathway | 50 | 1.38E-04 | 3.02E-02 | 7 | C2, C2b, C4a, C4, C2a, C3 convertase (C2aC4b), C4b |
| Immune response_Classical complement pathway | 53 | 2.00E-04 | 3.02E-02 | 7 | C2, C2b, C4a, C4, C2a, C3 convertase (C2aC4b), C4b |
| Role of IL-23/ T17 pathogenic axis in psoriasis | 54 | 2.25E-04 | 3.02E-02 | 7 | STAT3, IL-15, sIL-15RA, IL-15RA, CD86, CD40(TNFRSF5), HLA-C |
| Main growth factor signaling cascades in multiple myeloma cells | 41 | 3.26E-04 | 3.64E-02 | 6 | STAT3, MEK1(MAP2K1), HIF1A, I-kB, FGFR3, PI3K reg class IA |
| Cell adhesion_Integrin inside-out signaling in neutrophils | 77 | 3.84E-04 | 3.67E-02 | 8 | MEK1(MAP2K1), ARF6, LTBR1, MEK1/2, Cytohesin1, Lyn, Hck, GRO-1 |
| Development_c-Kit ligand signaling pathway during hemopoiesis | 61 | 4.84E-04 | 4.06E-02 | 7 | STAT3, MGF, MEK1/2, DOK1, JAK2, Lyn, SLUG |
| Immune response_Fc gamma R-mediated phagocytosis in macrophages | 46 | 6.16E-04 | 4.59E-02 | 6 | ARF6, Vinculin, Lyn, Hck, Profilin, PI3K reg class IA |
| Immune response_CD40 signaling | 65 | 7.15E-04 | 4.79E-02 | 7 | STAT3, I-kB, JAK2, Lyn, A20, CD86, CD40(TNFRSF5) |
| Development_NOTCH-induced EMT | 19 | 8.19E-04 | 4.85E-02 | 4 | HEY1, VE-cadherin, NOTCH4, SLUG |
| Signal transduction_Soluble CXCL16 signaling | 49 | 8.69E-04 | 4.85E-02 | 6 | STAT3, HIF1A, I-kB, MEK1/2, JAK2, CD163 |
| Development GM-CSF sig- | 50 | 9.68E-04 | 4.99E-02 | 6 | STAT3, MEK1(MAP2K1), |

|  |  |
| --- | --- |
| naling | I-kB, JAK2, Lyn, Hck |

**Supplementary Table S9.** Process networks enrichment analysis of the 830 genes in cluster 2 in the H1N1 integrative network using MetaCore. Only the top 50 significantly enriched process networks are shown here.

| Process networks | Total | *p* | FDR | In Data | Network Objects from Active Data |
| --- | --- | --- | --- | --- | --- |
| Immune re-<br>sponse_Phagosome in<br>antigen presentation | 243 | 7.255E-05 | 1.110E-02 | 22 | PSMA4, NFKBIB, ARF6, Vinculin, I-kB, PSMD3, Profilin II, Lyn, PSMA1, Alpha-actinin, PSMD14, Hck, TLR4, PSMA2, Profilin, PSMD12, HLA-C, HLA-B, Alpha-actinin 3, PI3K reg class IA (p55-gamma), NCK1, PI3K reg class IA |
| Immune re-<br>sponse_Antigen<br>presentation | 197 | 2.965E-04 | 2.269E-02 | 18 | PSMA4, STAT3, NFKBIB, RFX5, PSMD8, I-kB, JAK2, PSMD3, PSMA1, HLA-E, PSMD14, PSMA2, CD86, CD40(TNFRSF5), PSMD12, HLA-C, HLA-B, CD209 |
| Inflammation_Interferon<br>signaling | 110 | 6.422E-04 | 3.275E-02 | 12 | STAT3, MEK1(MAP2K1), MEK1/2, JAK2, MxA, Pyrin (MEFV), ISG20, CD86, SSAT, CD40(TNFRSF5), IFNAR2, SERPINB9 |
| Signal transduc-<br>tion_Insulin signaling | 171 | 1.428E-03 | 5.461E-02 | 15 | Des (64,65) proinsulin, MEK1(MAP2K1), Split (32,33) proinsulin, Insulin processed, JAK2, Diarginyl insulin, Proinsulin C-peptide, PYGM, PKA-reg (cAMP-dependent), Split (64,65) proinsulin, Insulin, Proinsulin, Des(31,32) proinsulin, FADS2, PI3K reg class IA |
| Cell adhesion_Synaptic<br>contact | 184 | 2.936E-03 | 7.812E-02 | 15 | Syntaxin 4, Synaptotag-<br>min II, Synaptotagmin,<br>Profilin II, Ephrin-B2,<br>Ephrin-B, Ephrin-B2<br>(CTF2), Alpha-actinin,<br>SHANK, SynGAP,<br>Profilin, SHANK2, Al-<br>pha-actinin 3, Syntenin<br>1, NCK1 |
| Immune re-<br>sponse_Phagocytosis | 222 | 3.063E-03 | 7.812E-02 | 17 | IL-15, NFKBIB, Vincu-<br>lin, I-kB, MyHC, Pro-<br>filin II, Lyn, Alpha-actinin, IL-15RA, Hck,<br>TLR4, Profilin, Alpha-actinin 3, HDL pro-<br>teins, PI3K reg class IA<br>(p55-gamma), NCK1,<br>PI3K reg class IA |

**Supplementary Table S10.**
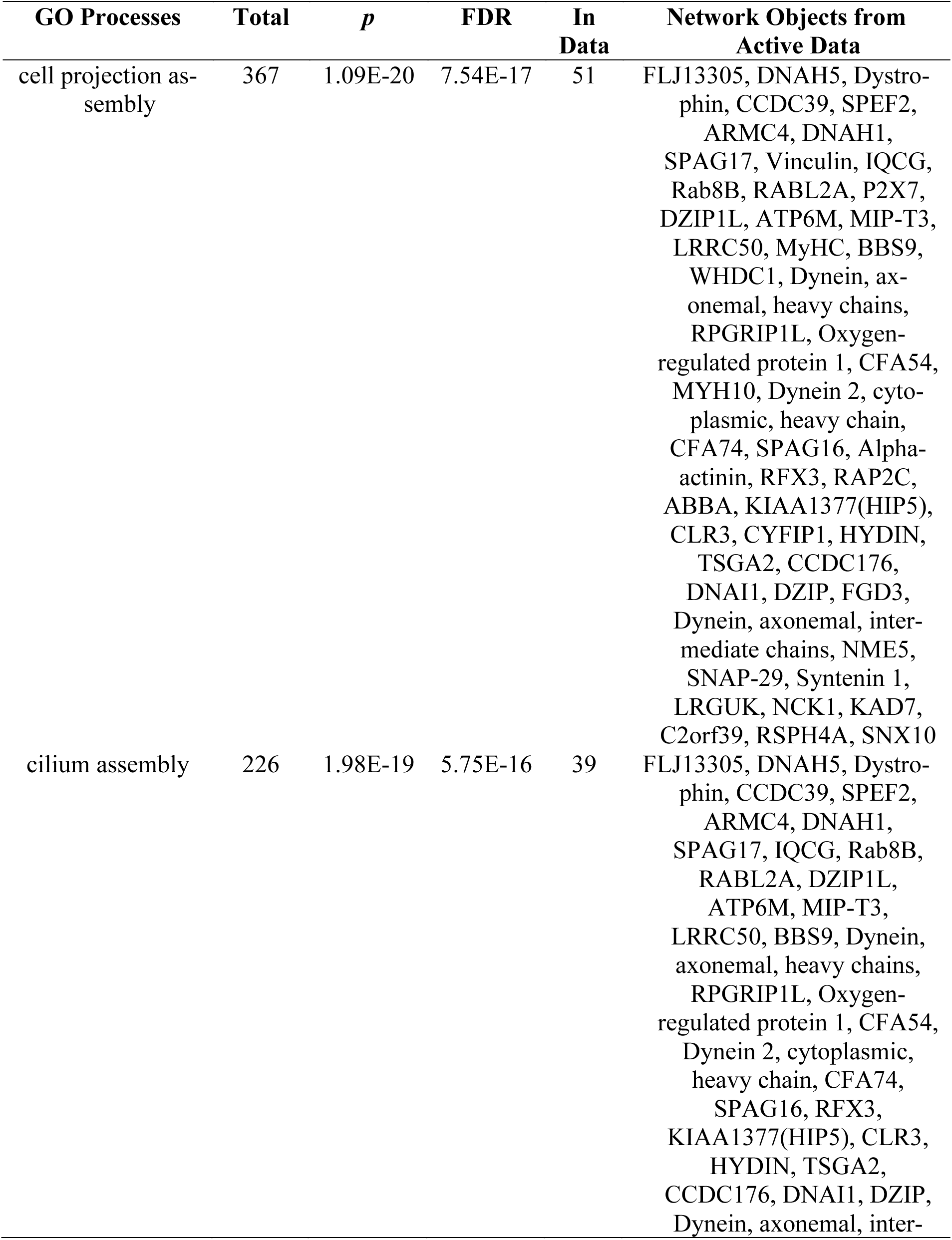
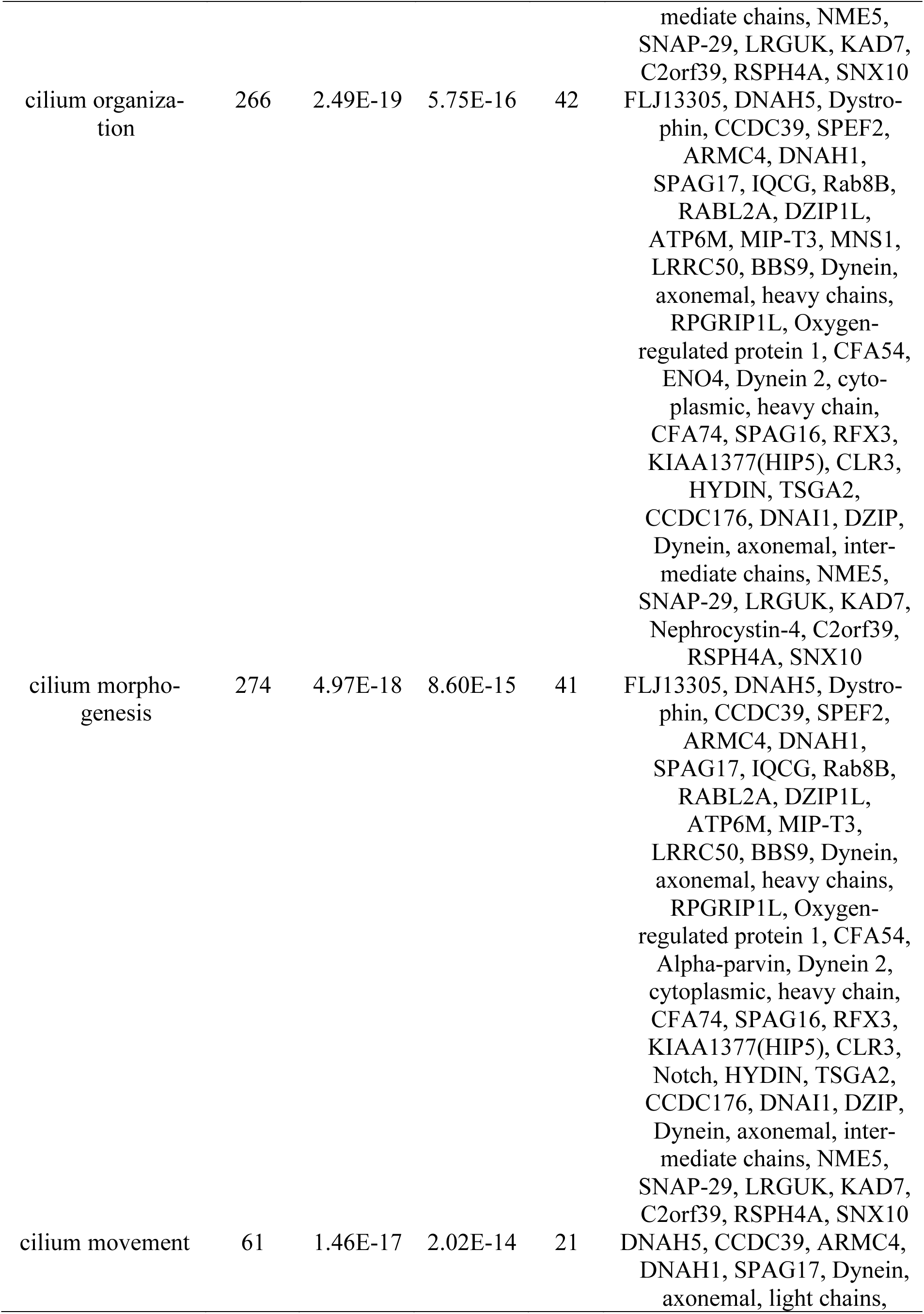
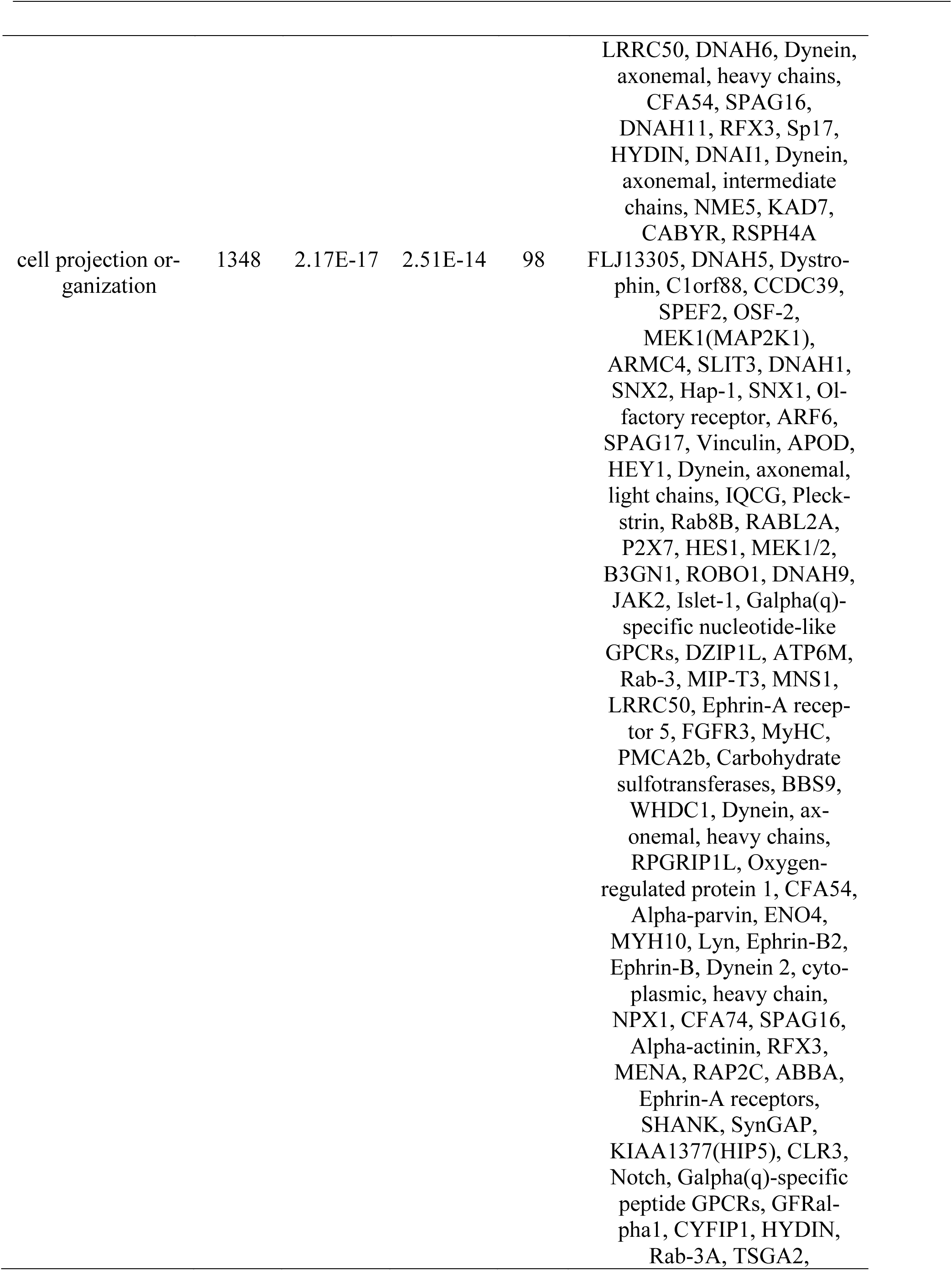
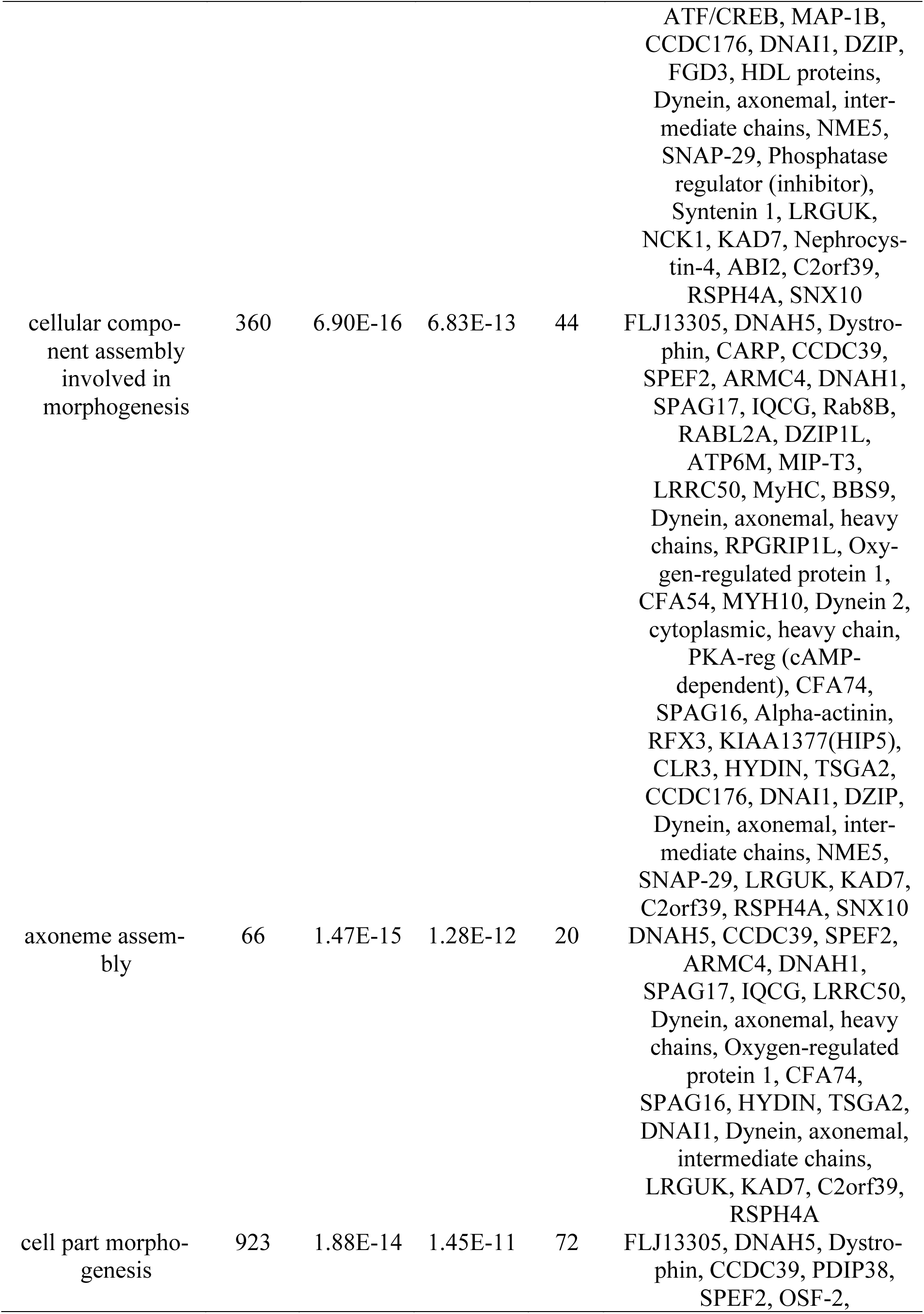
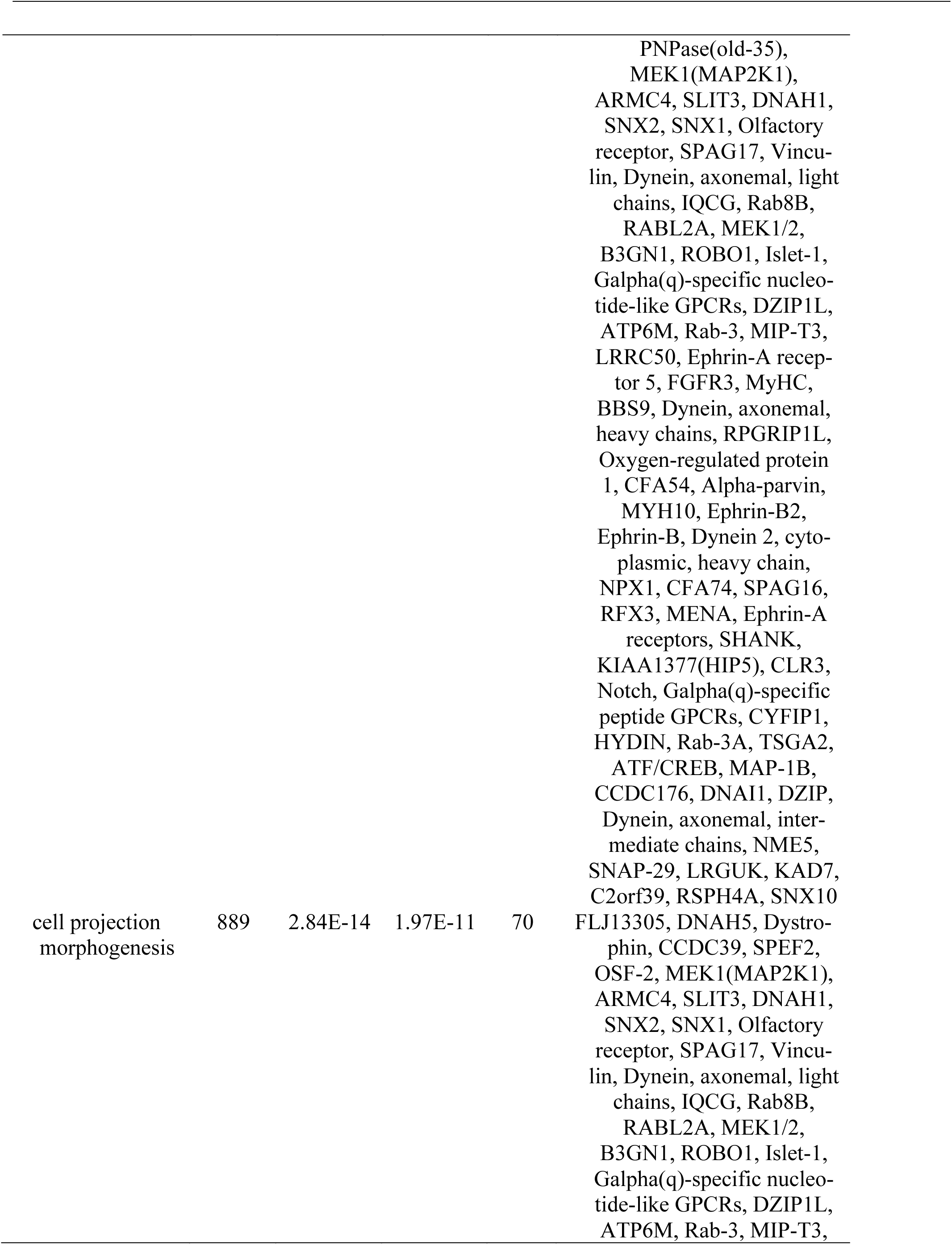
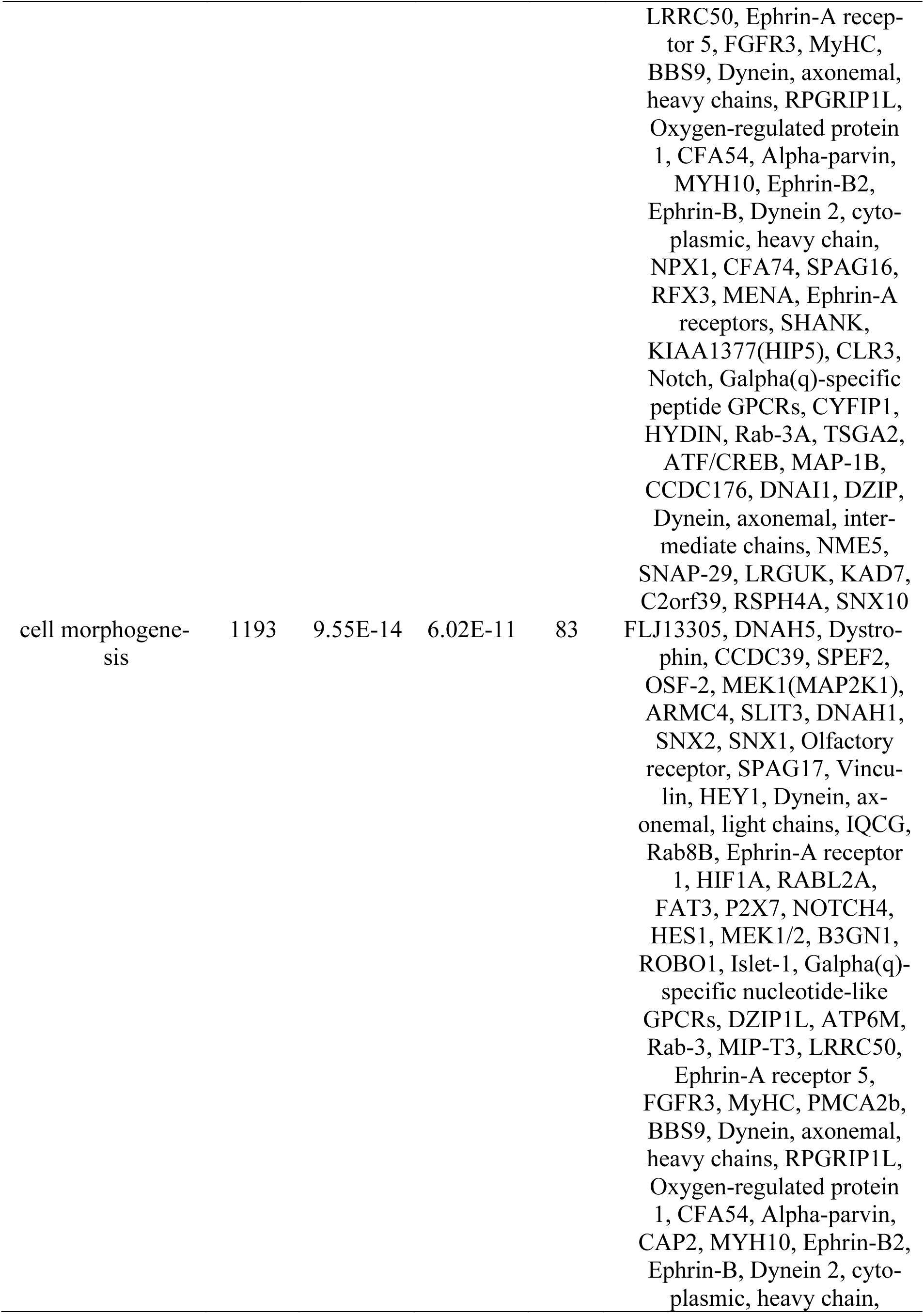
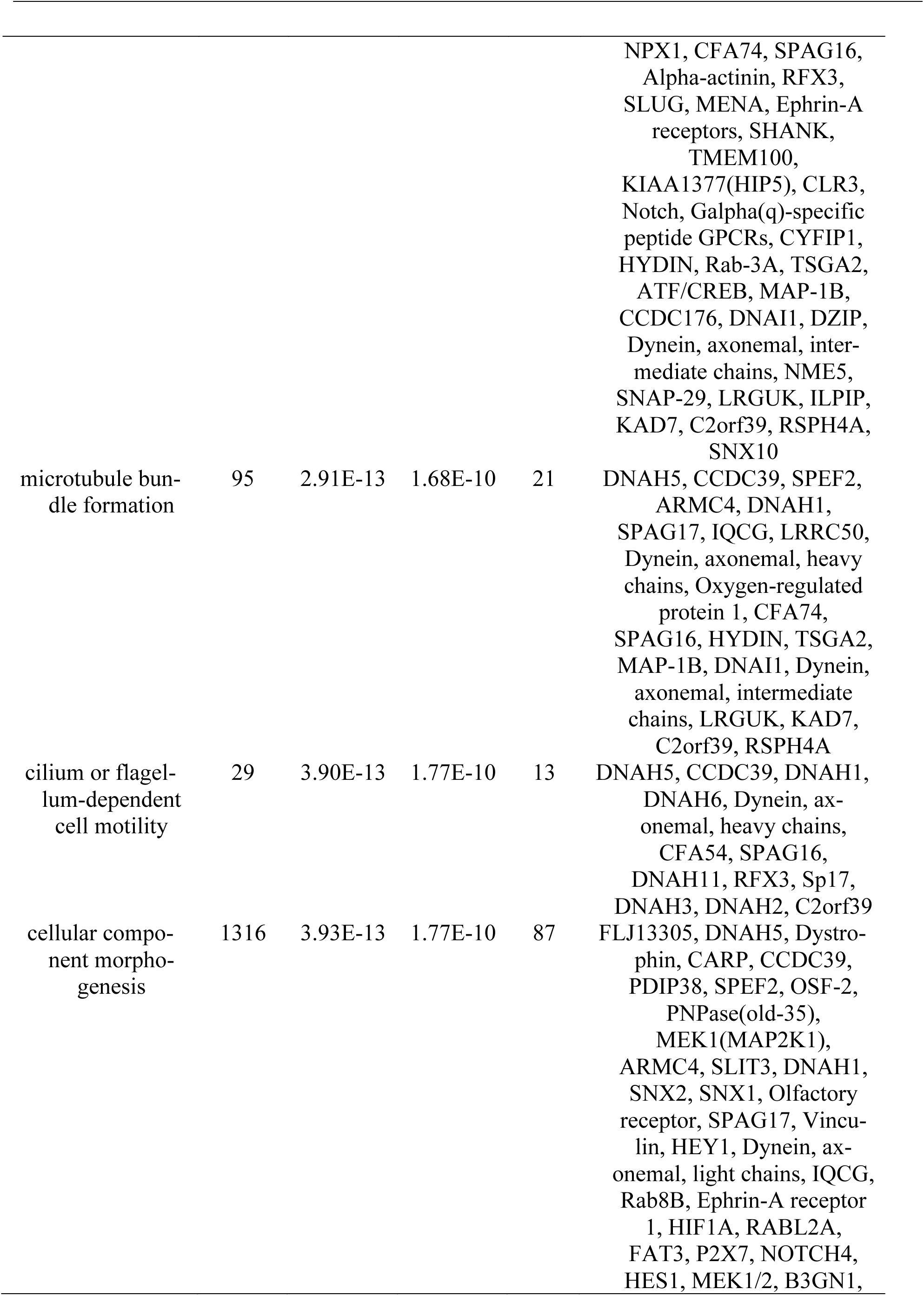
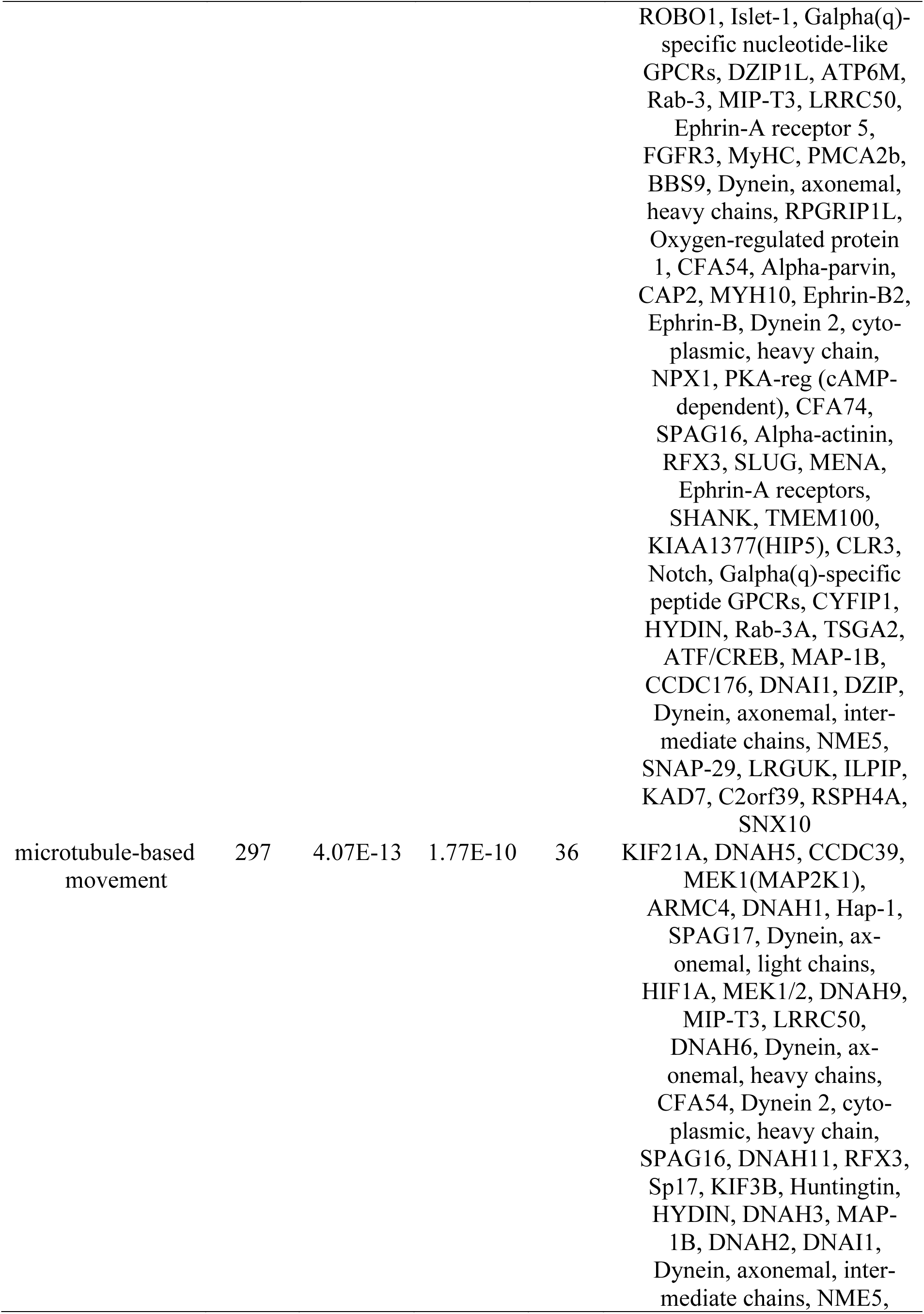
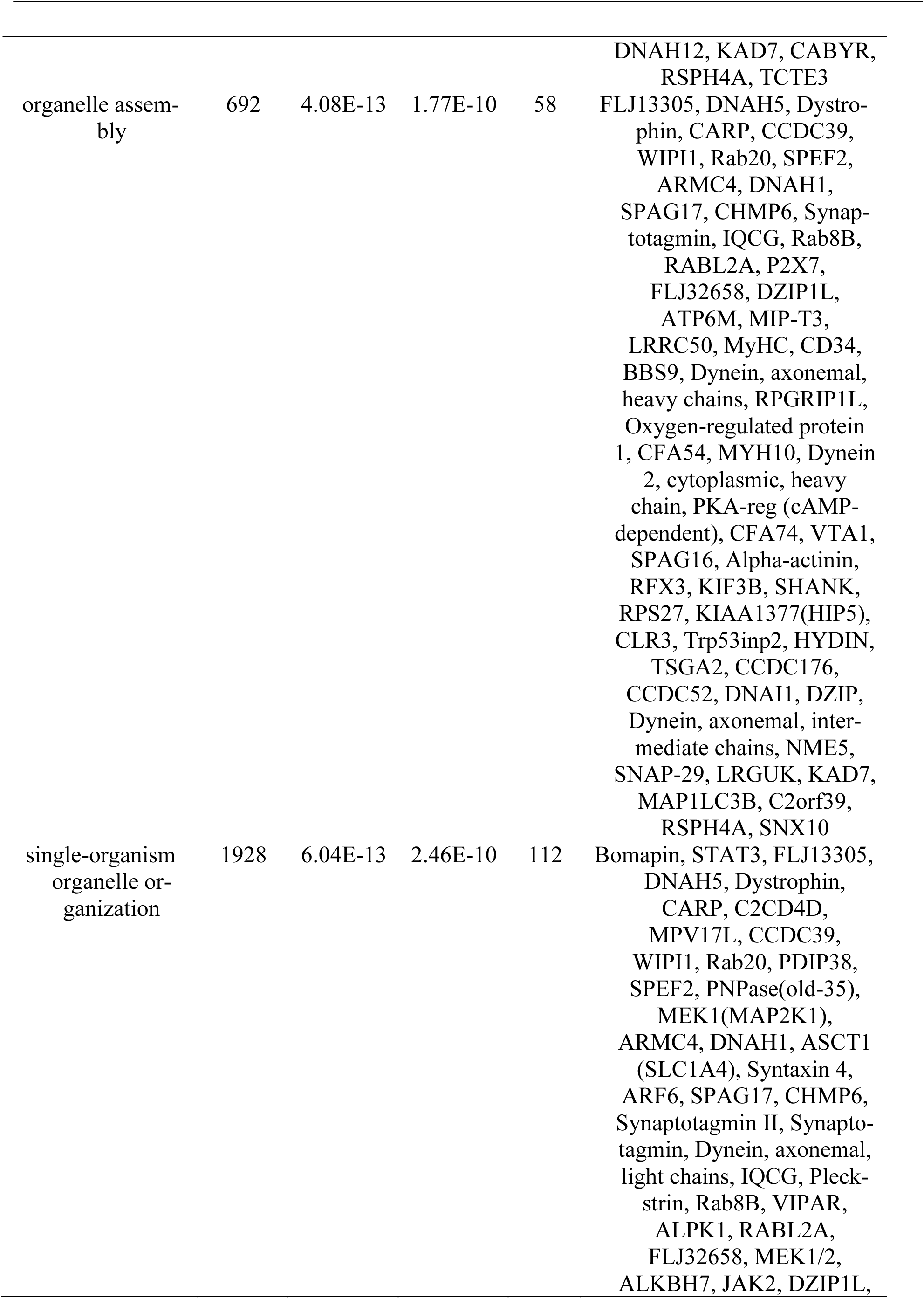
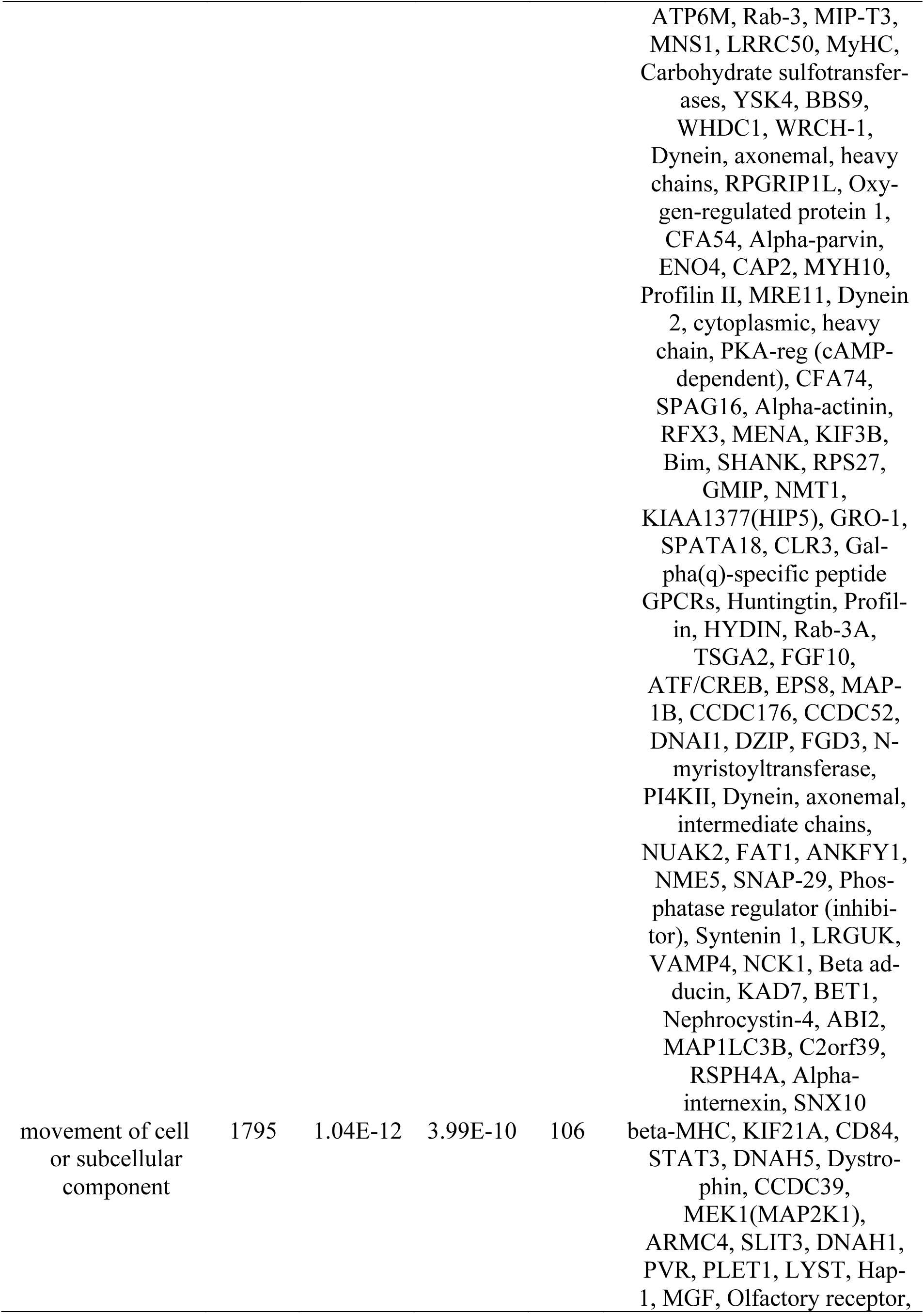
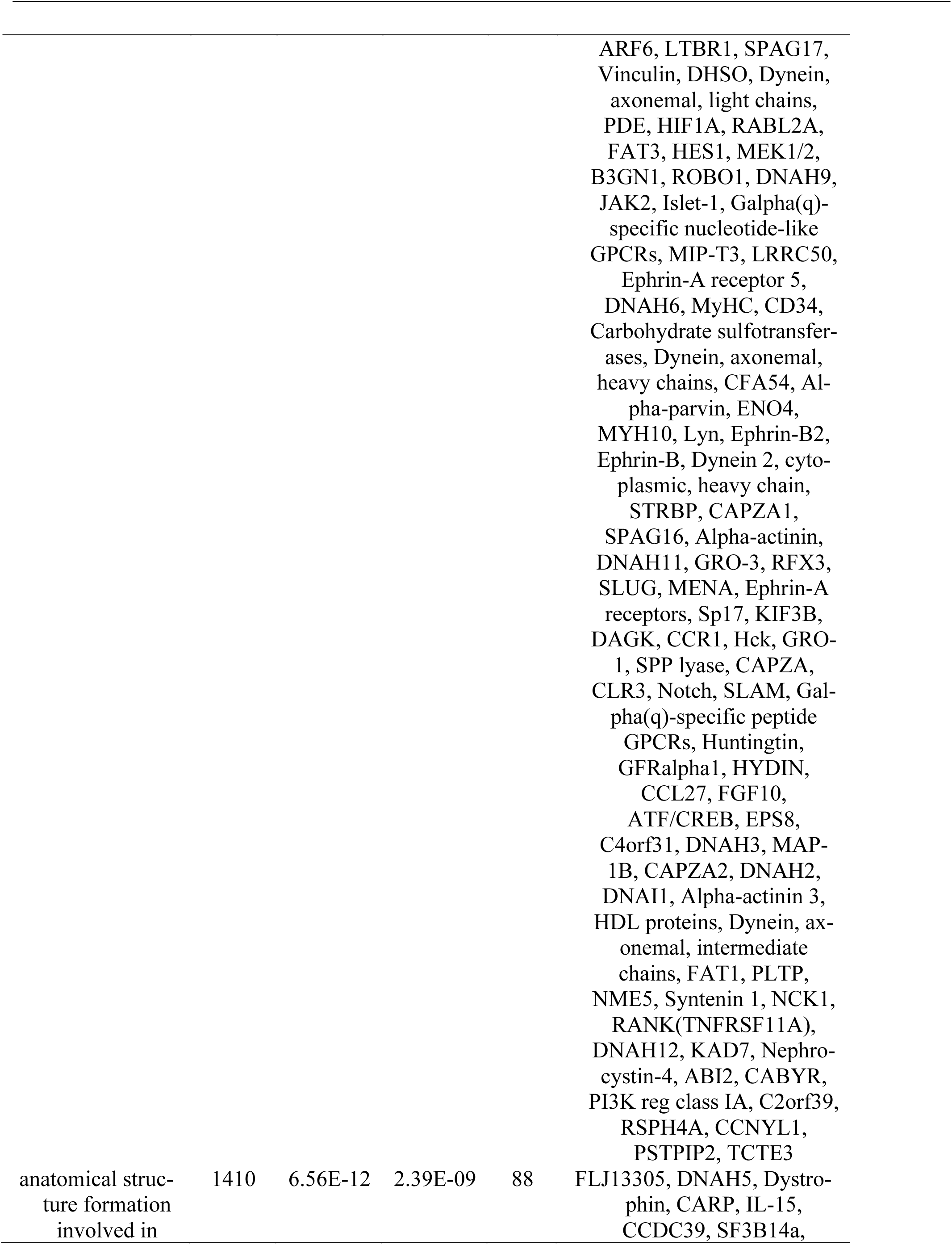
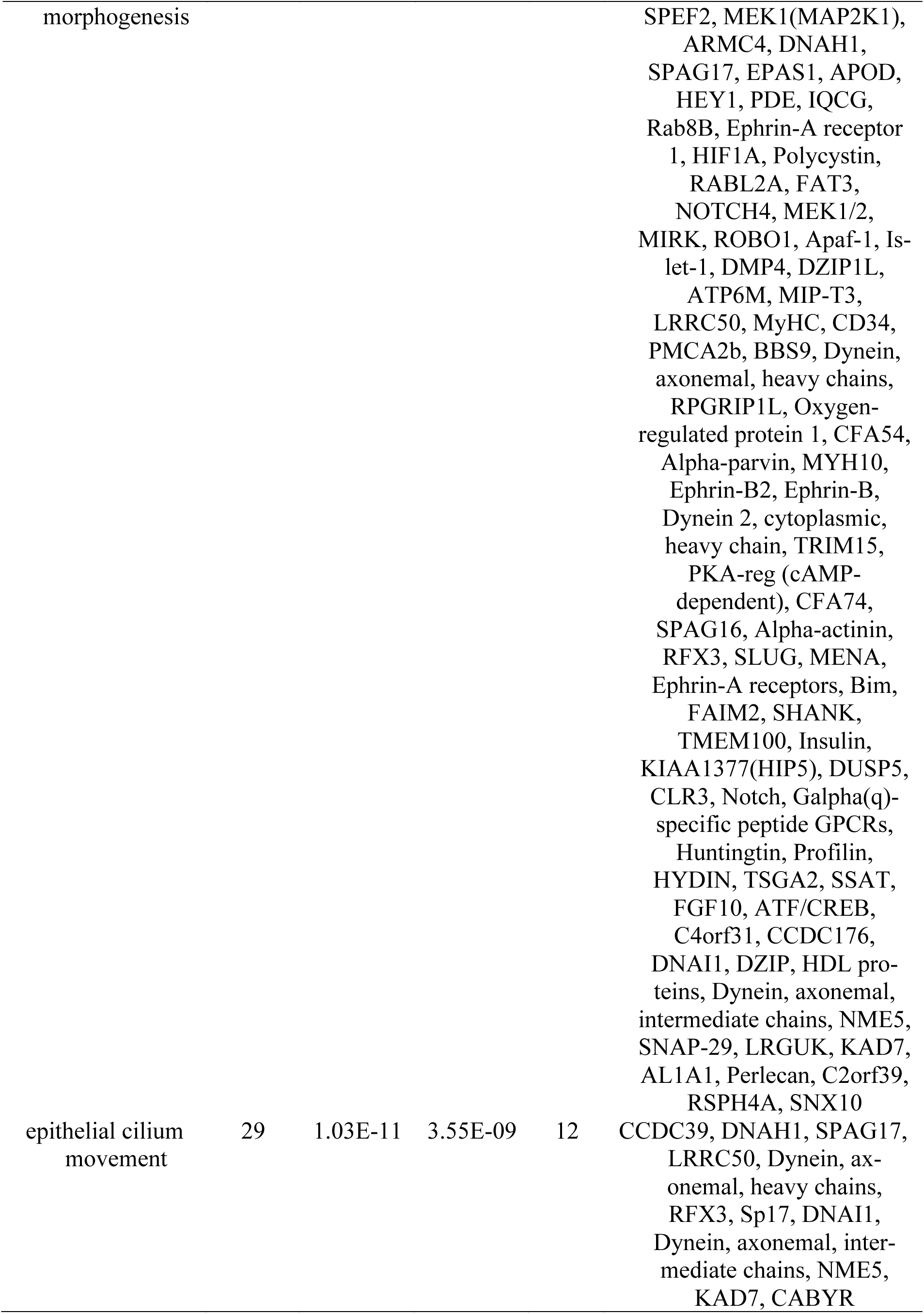
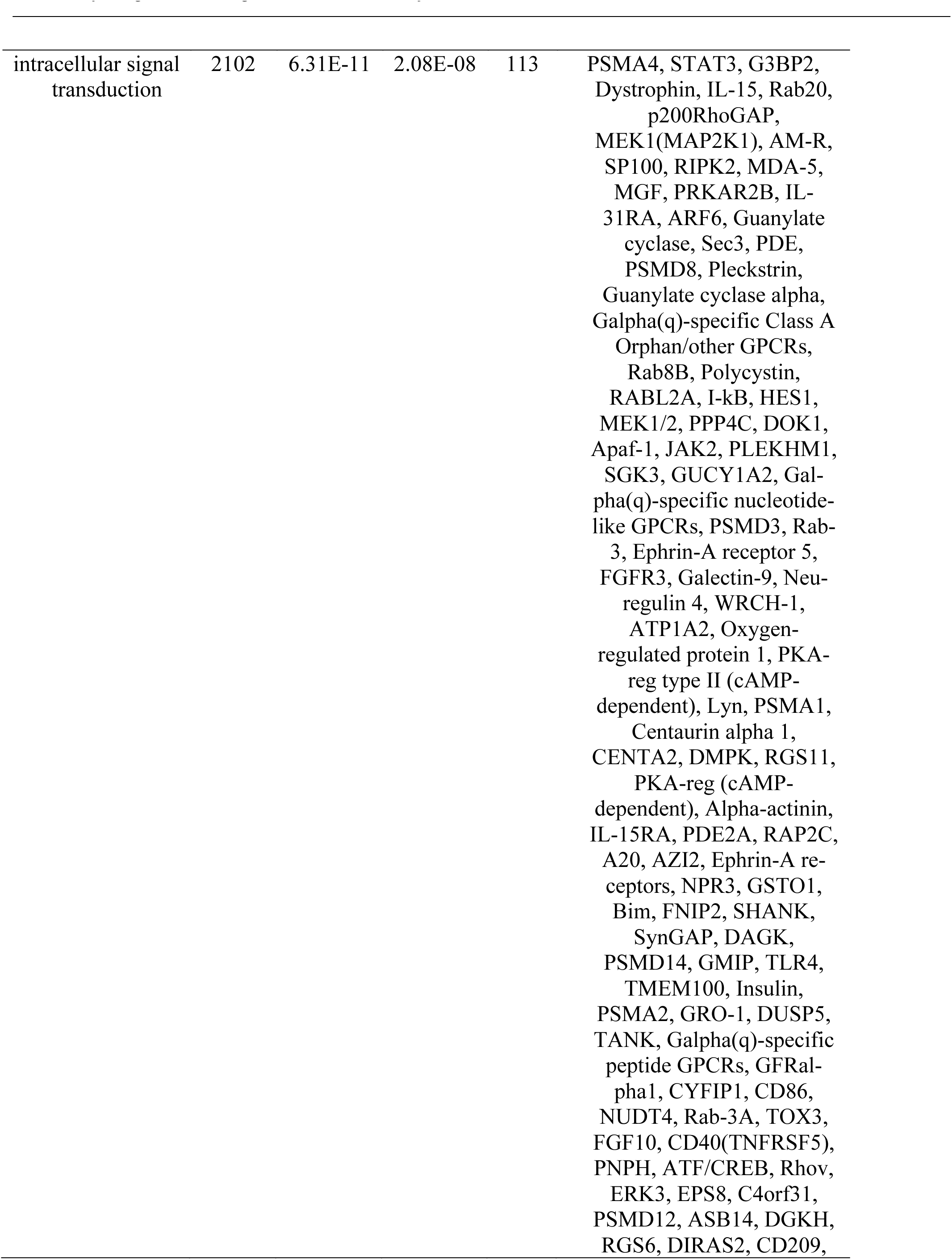
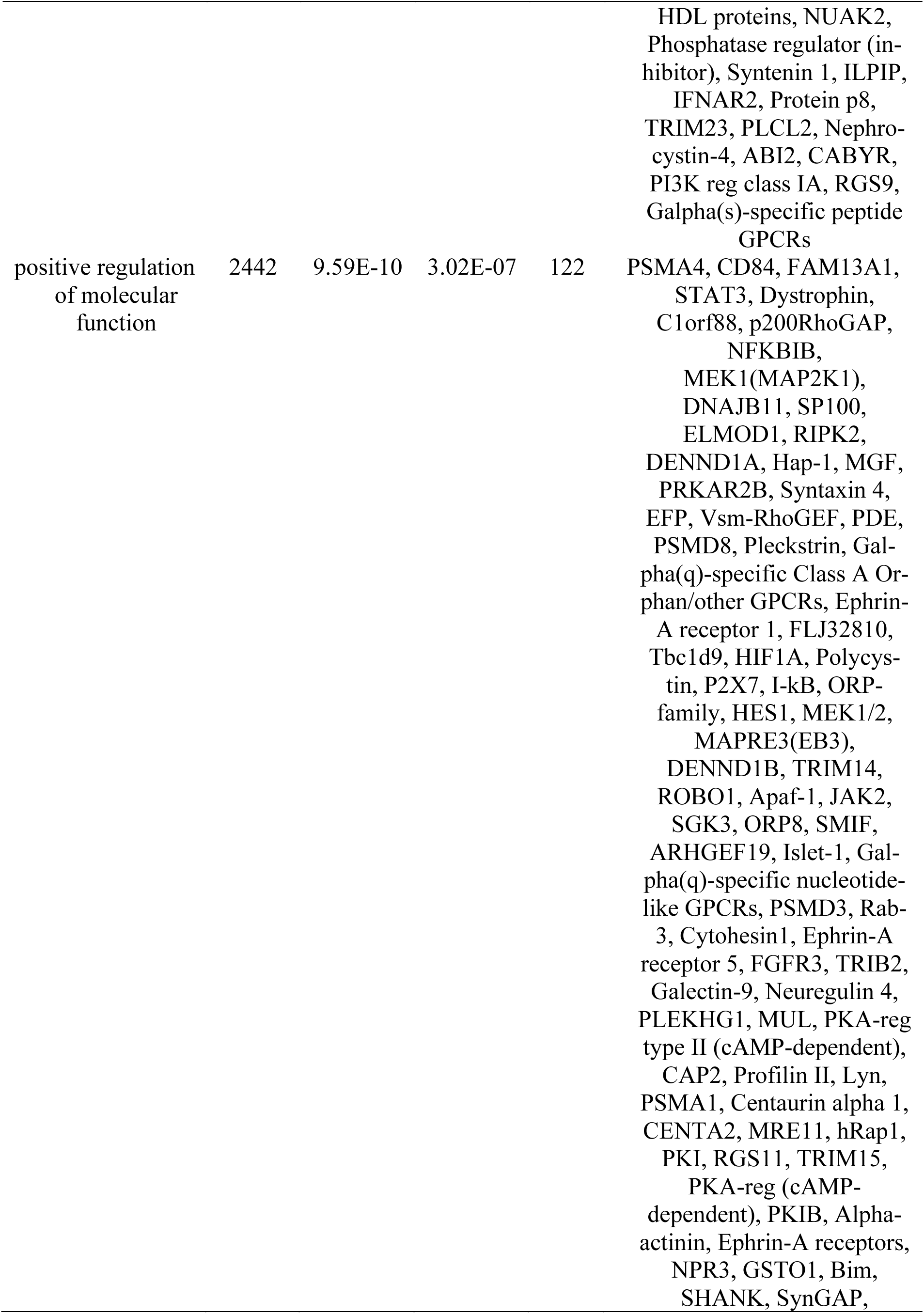
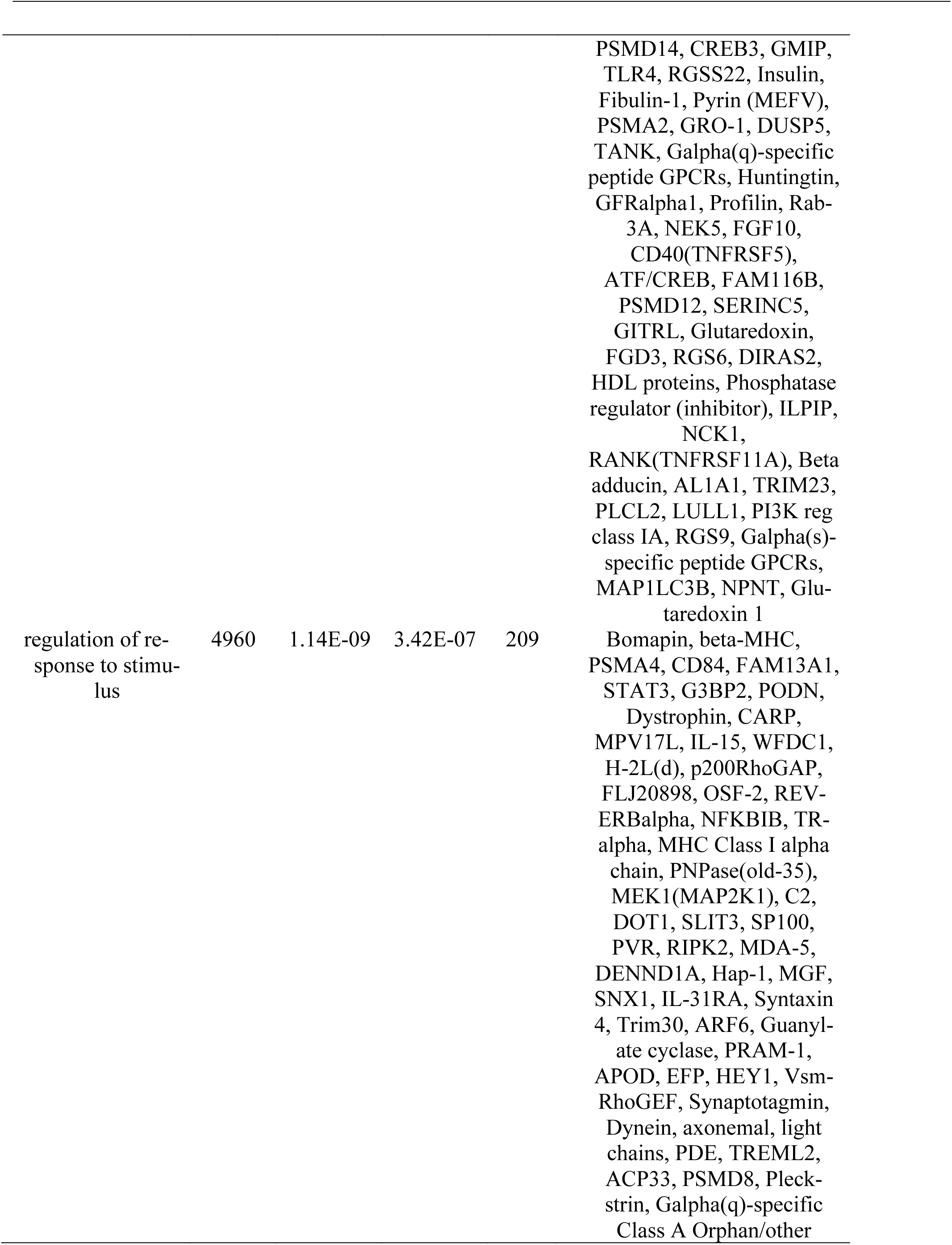
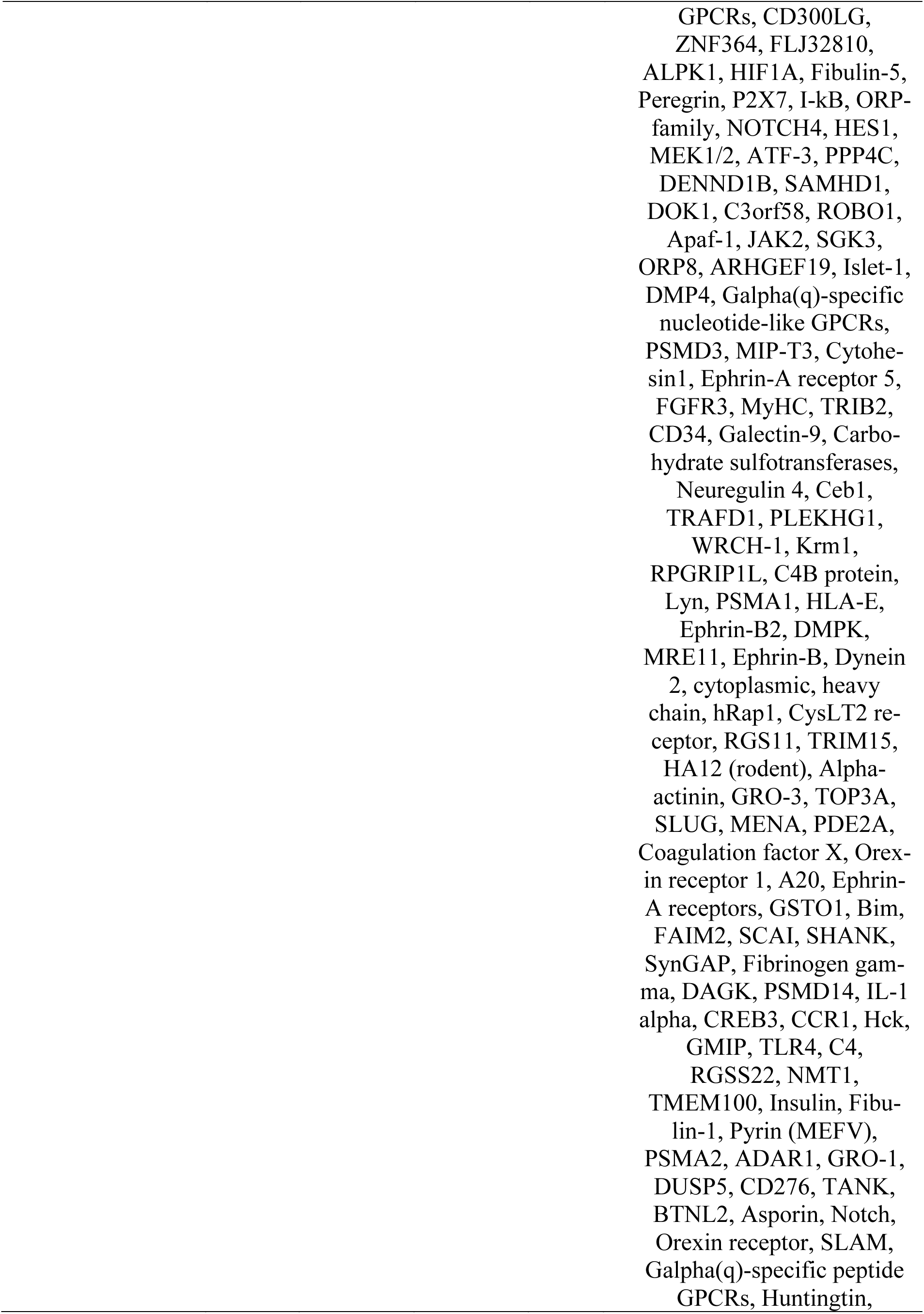
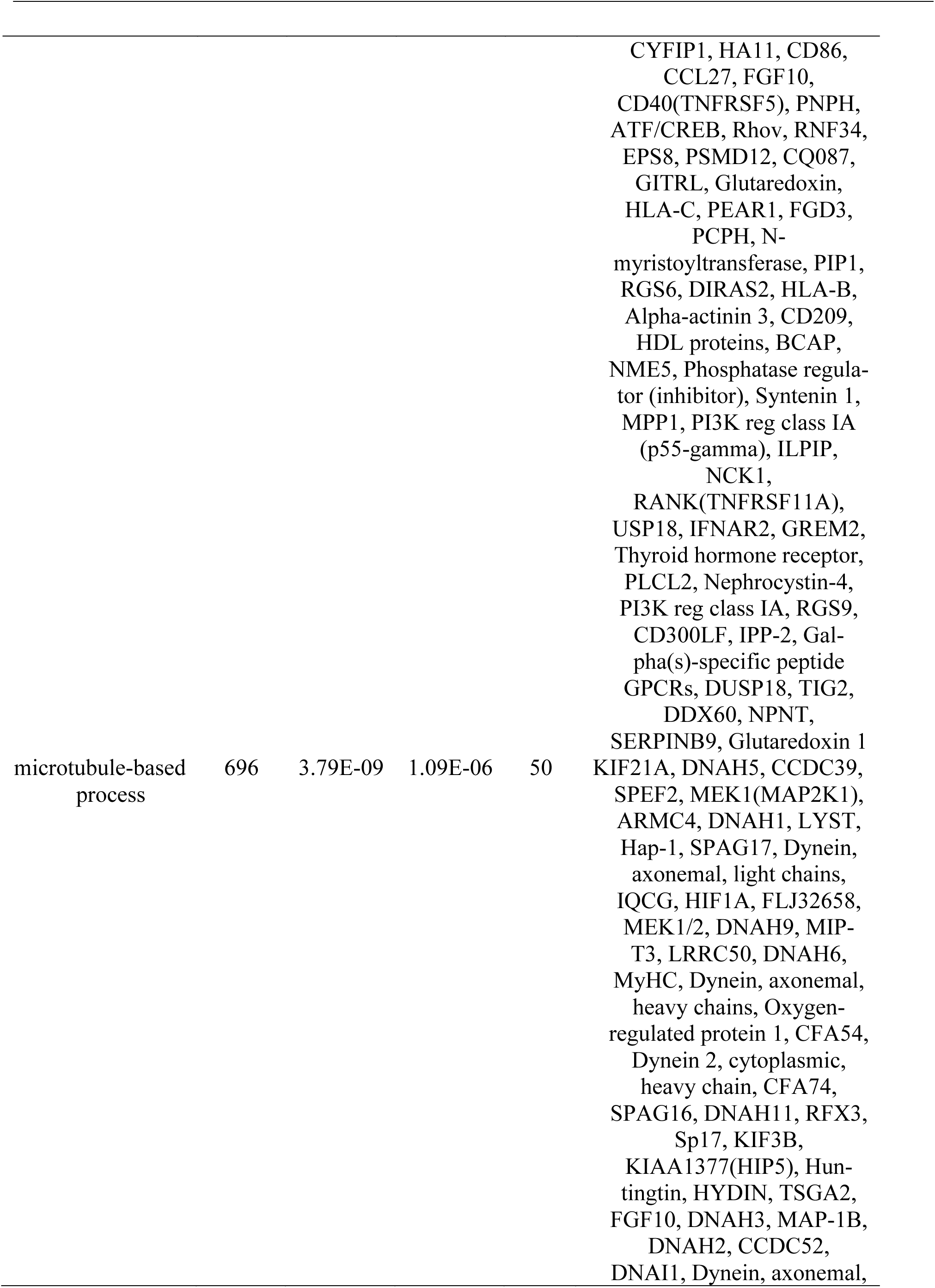
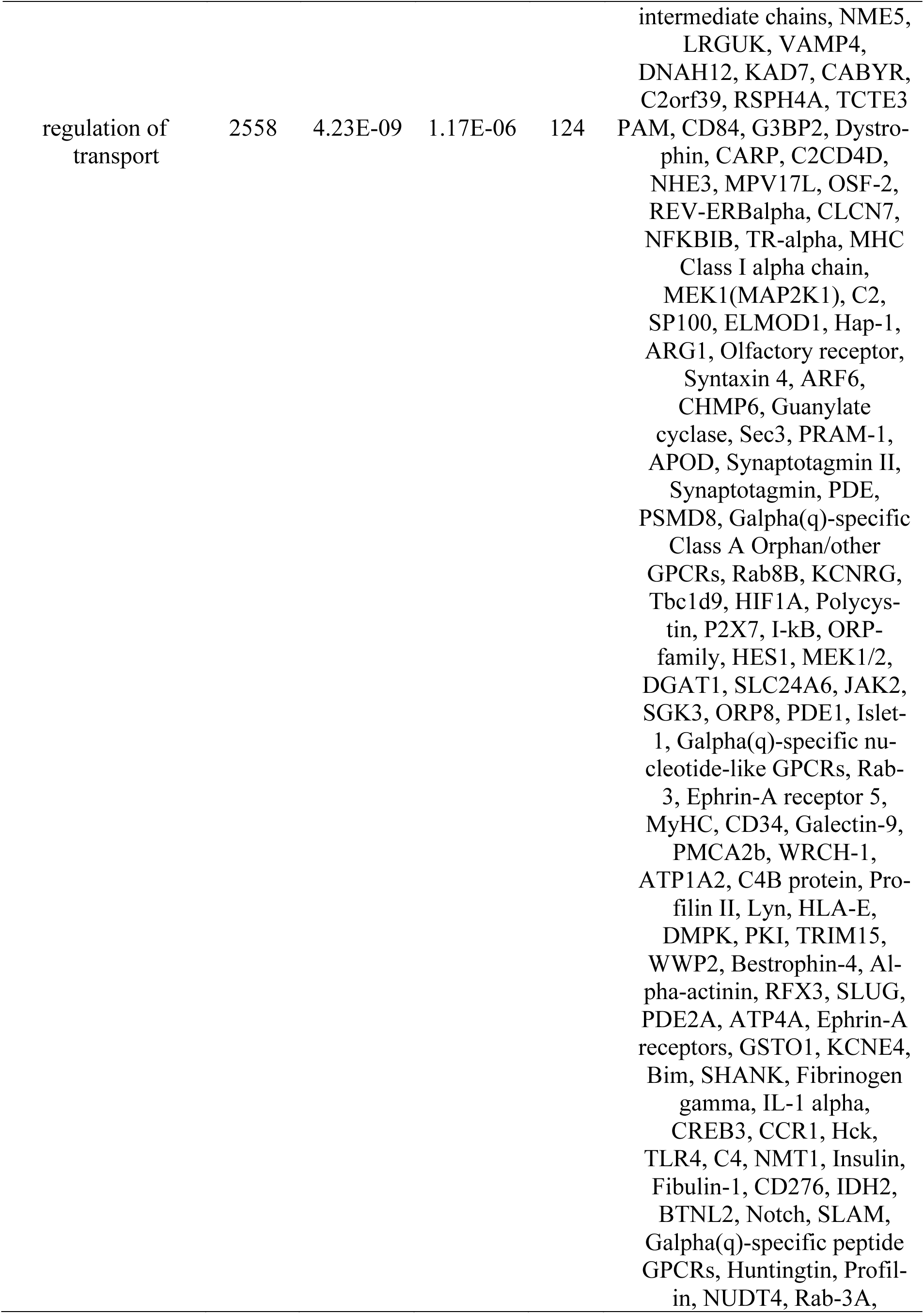
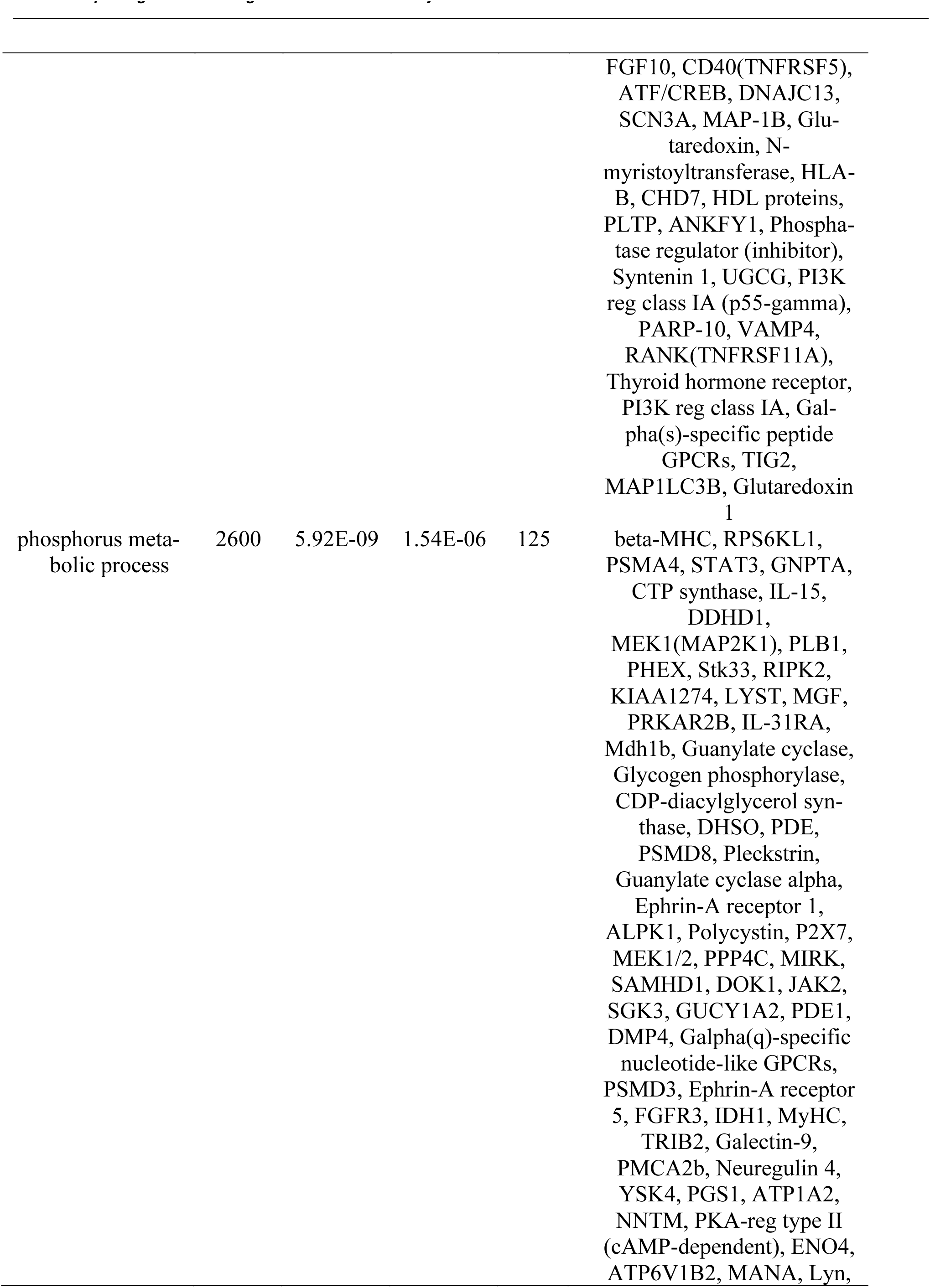
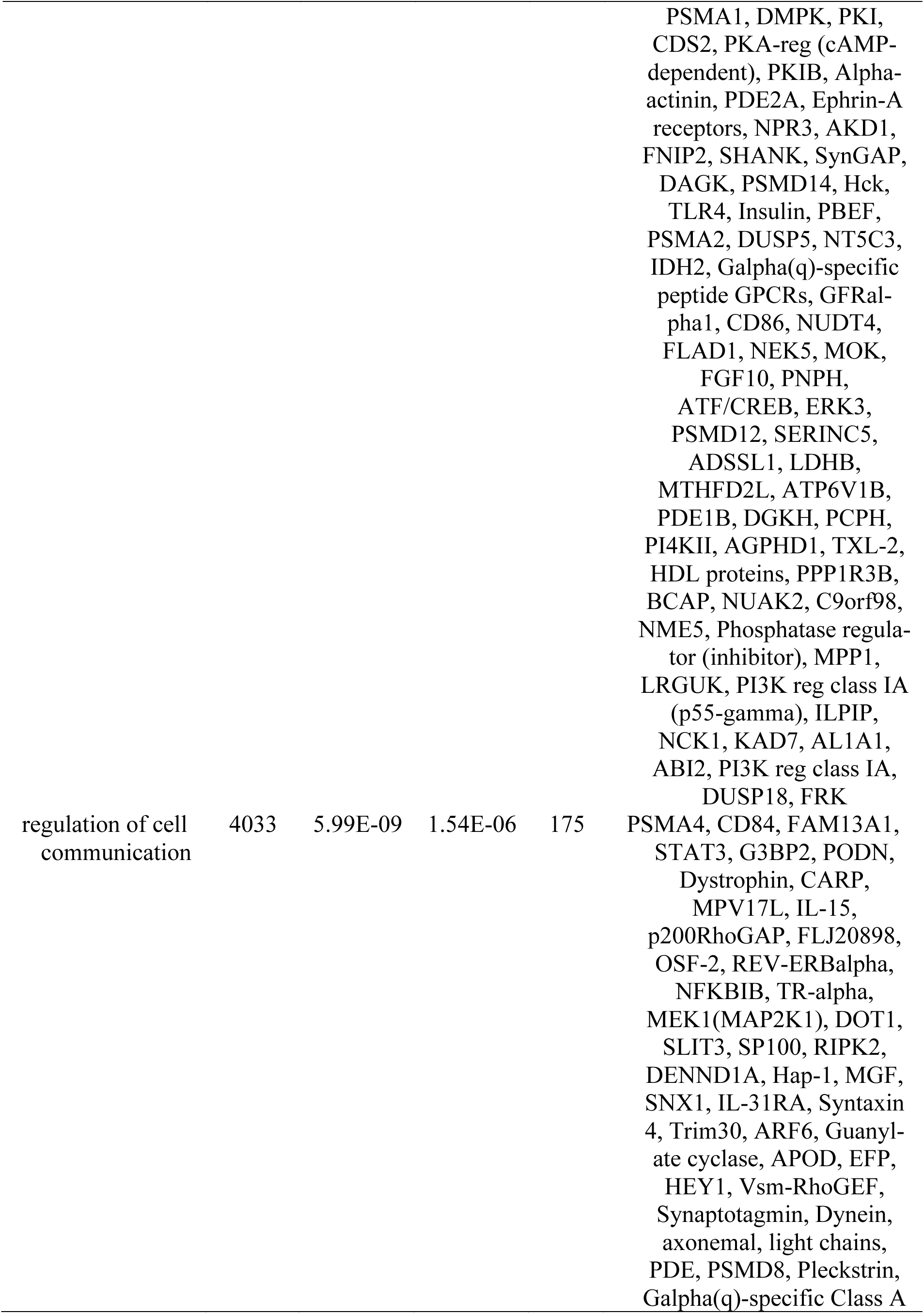
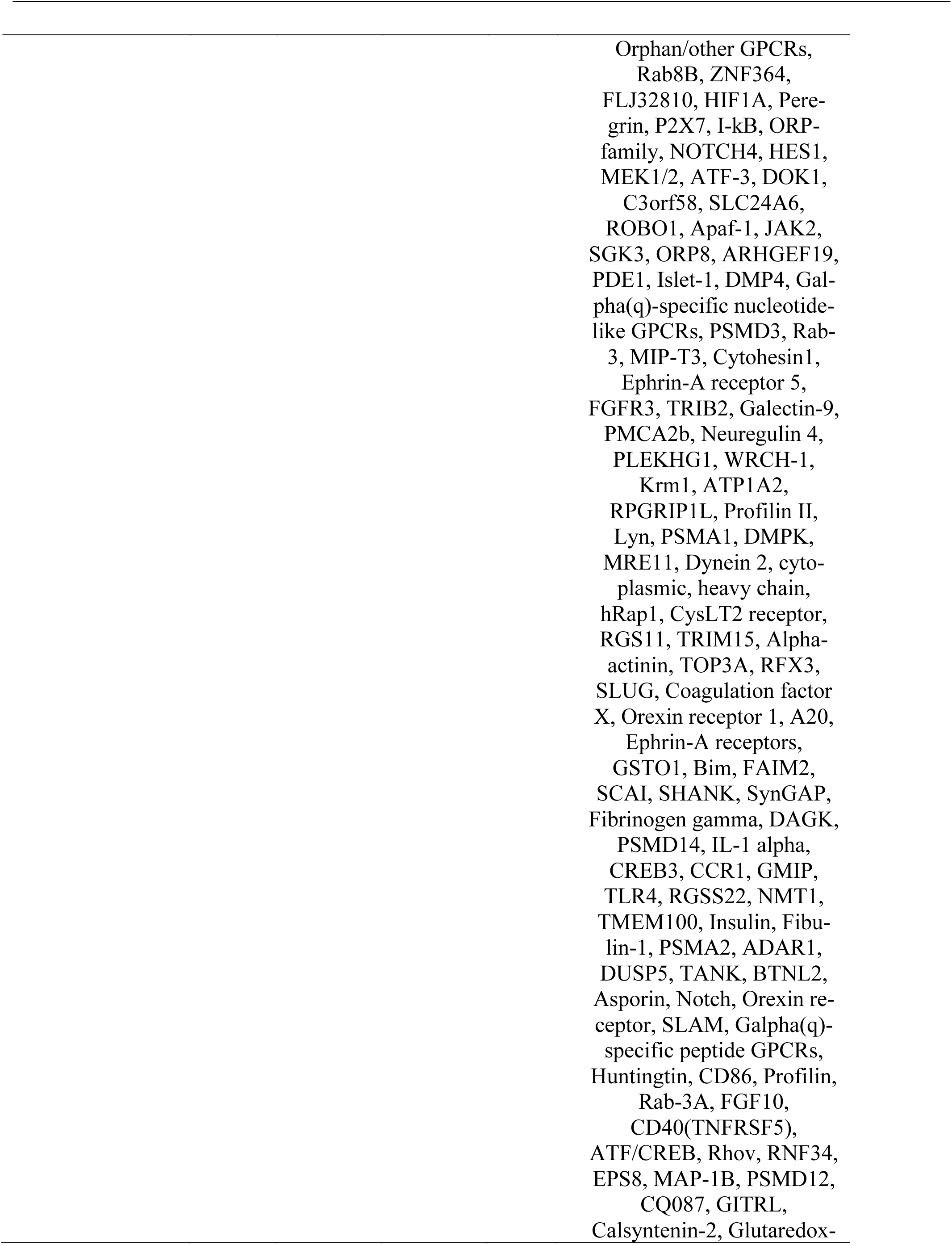
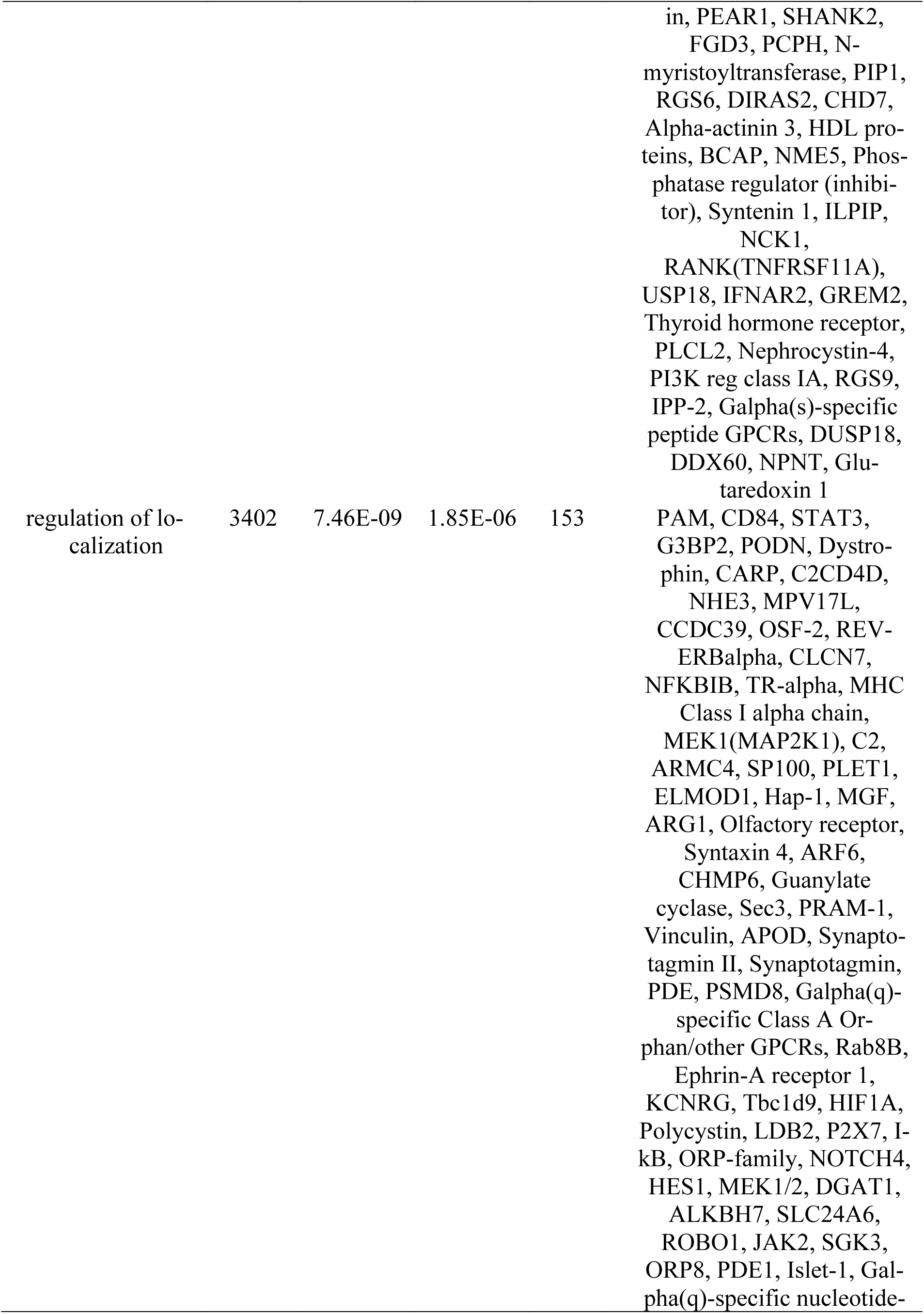
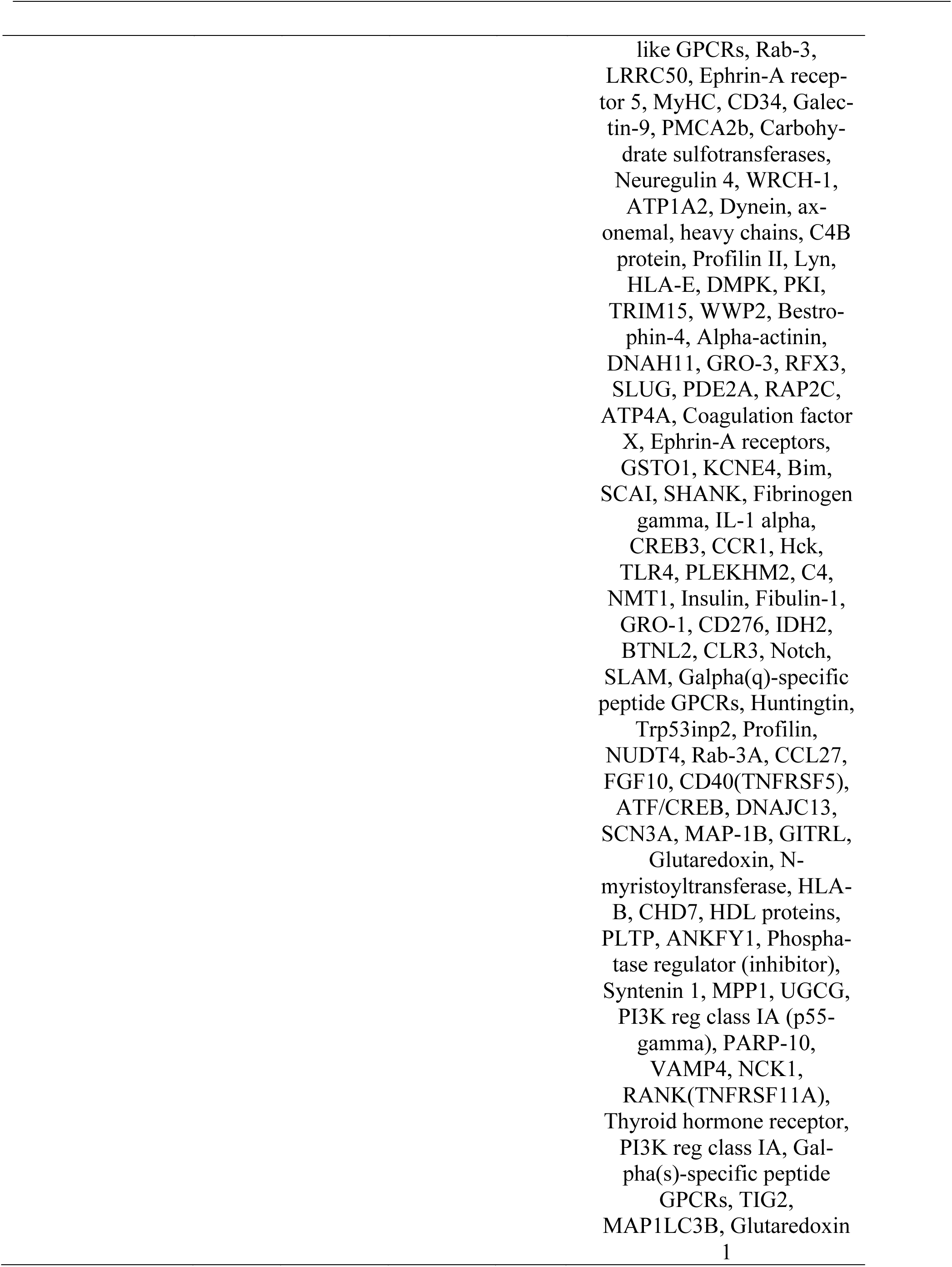
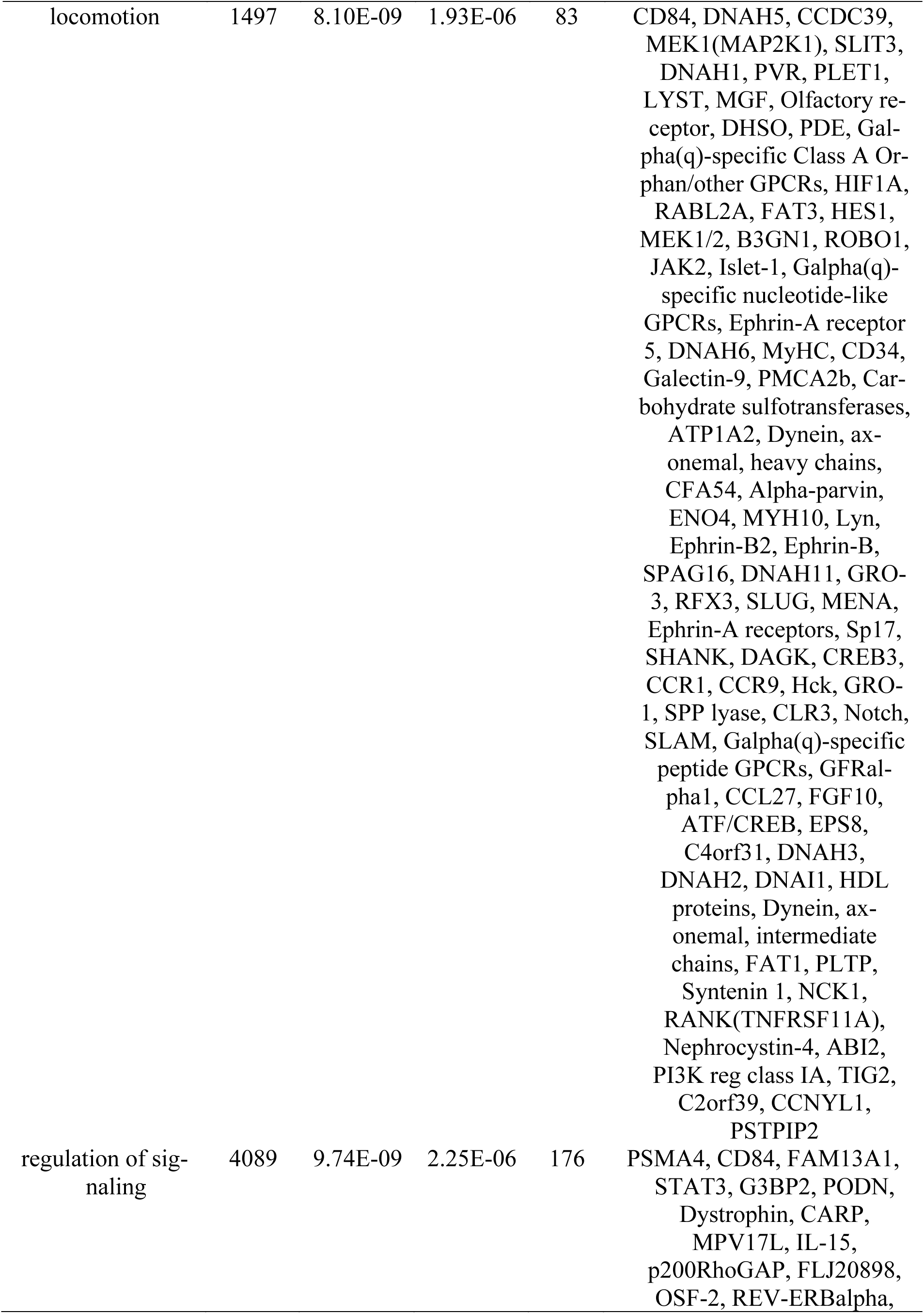
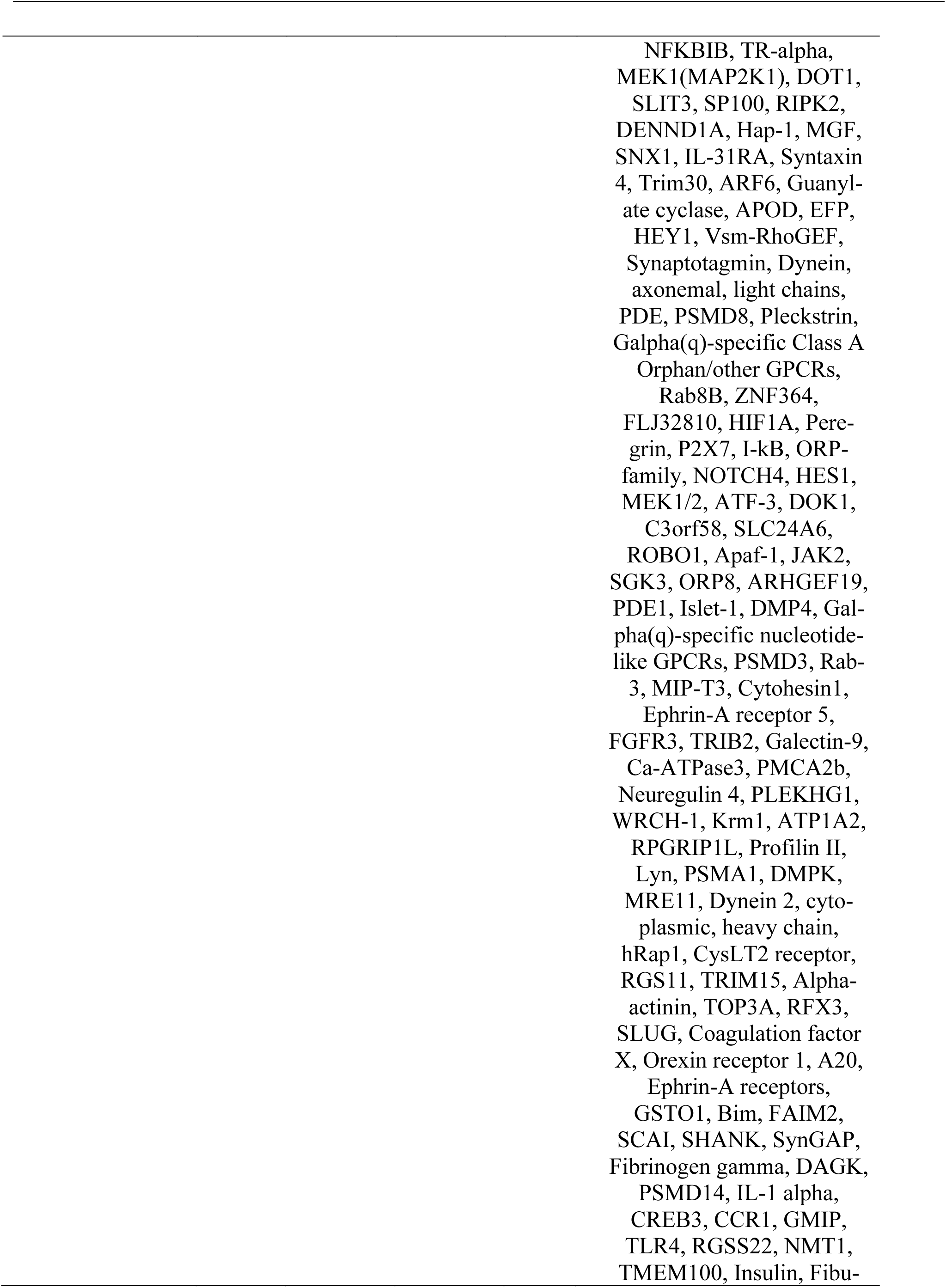
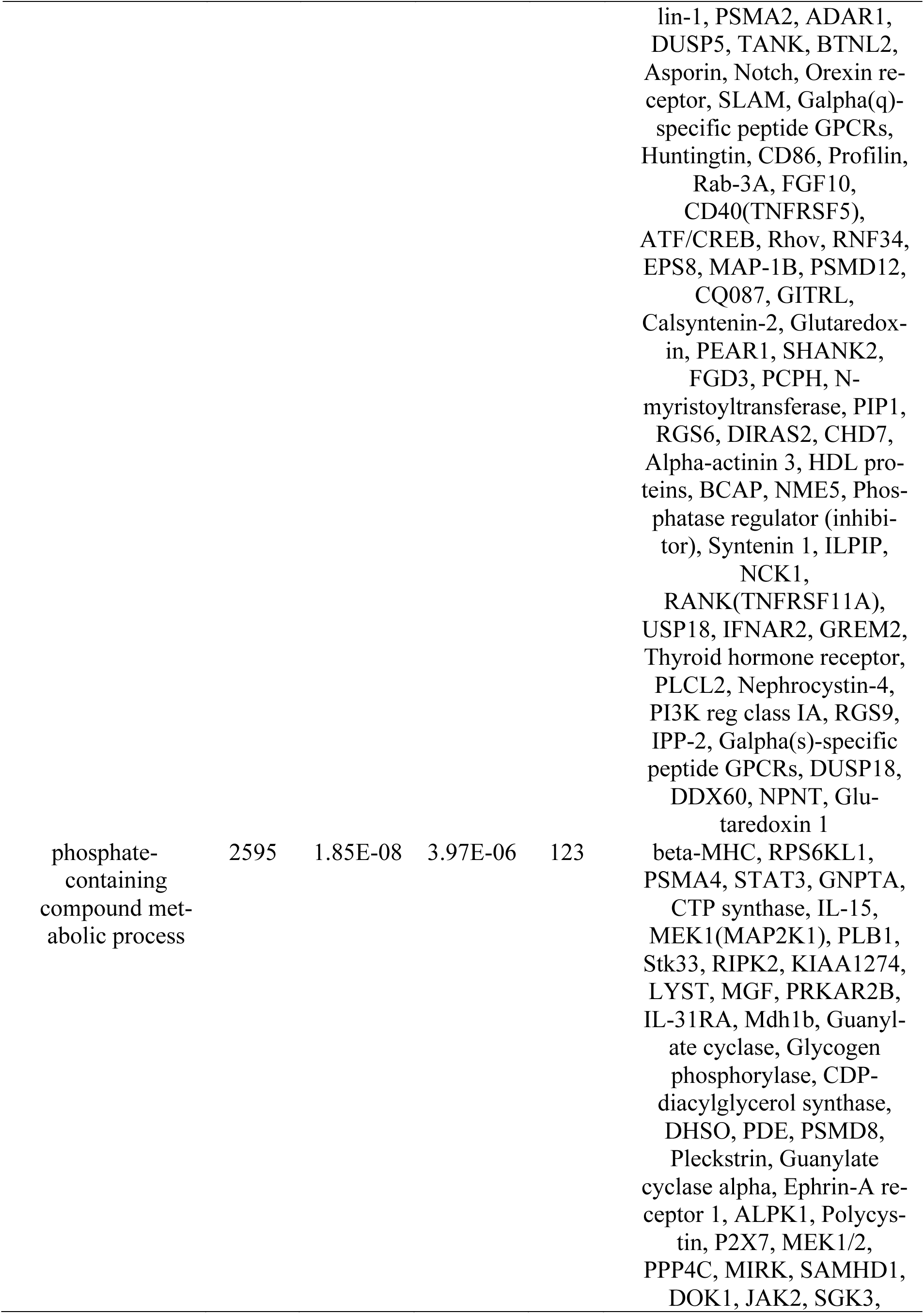
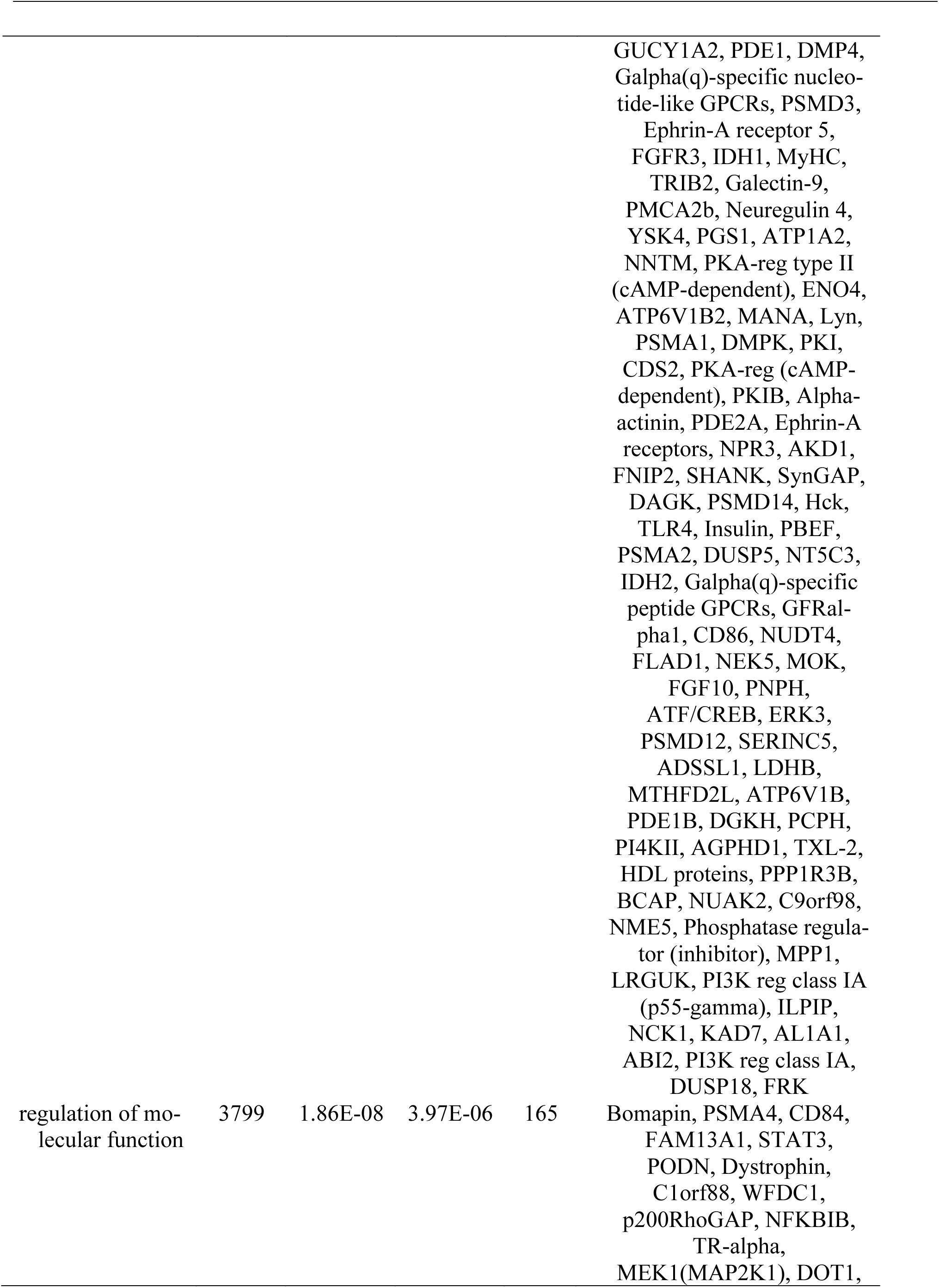
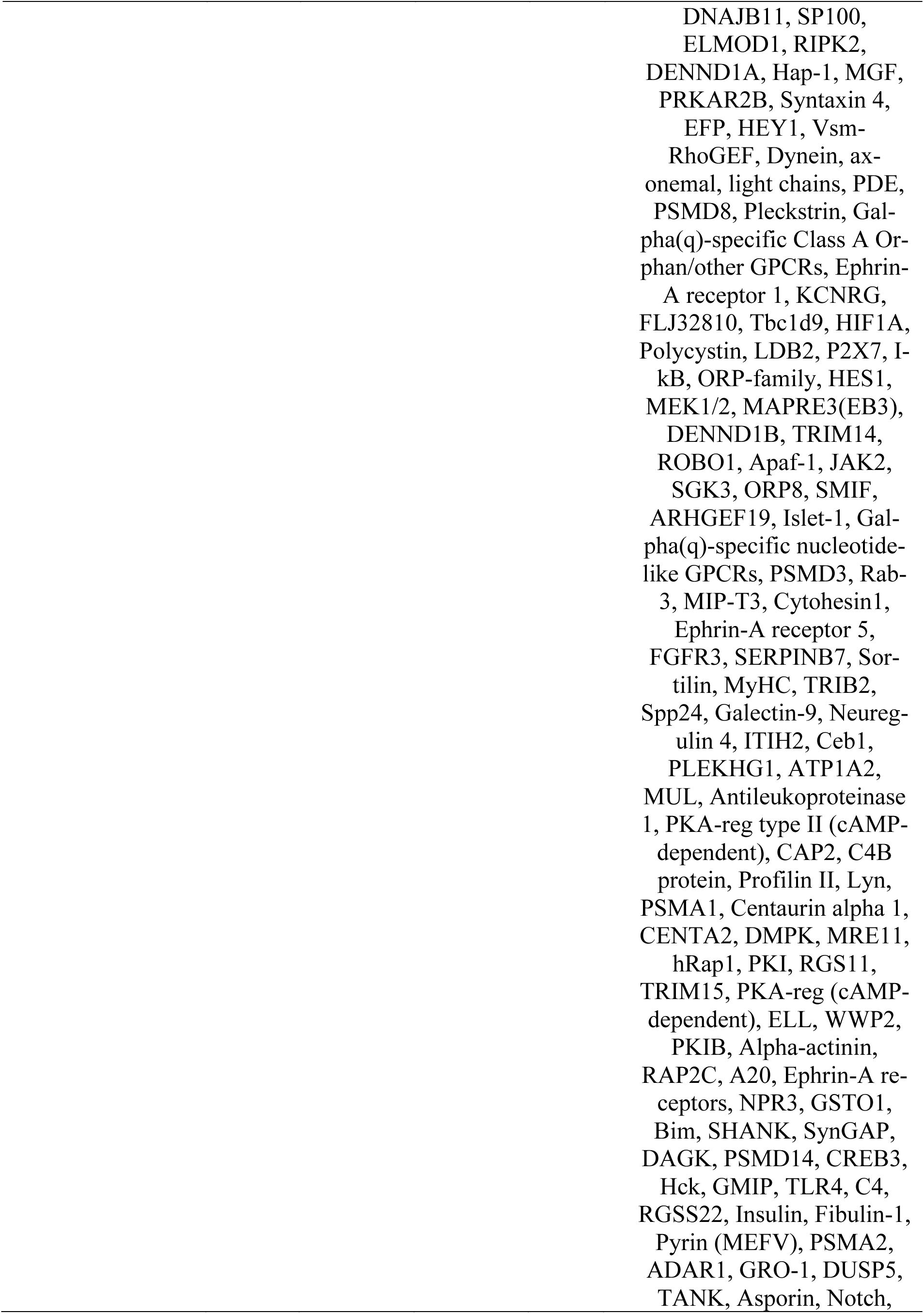
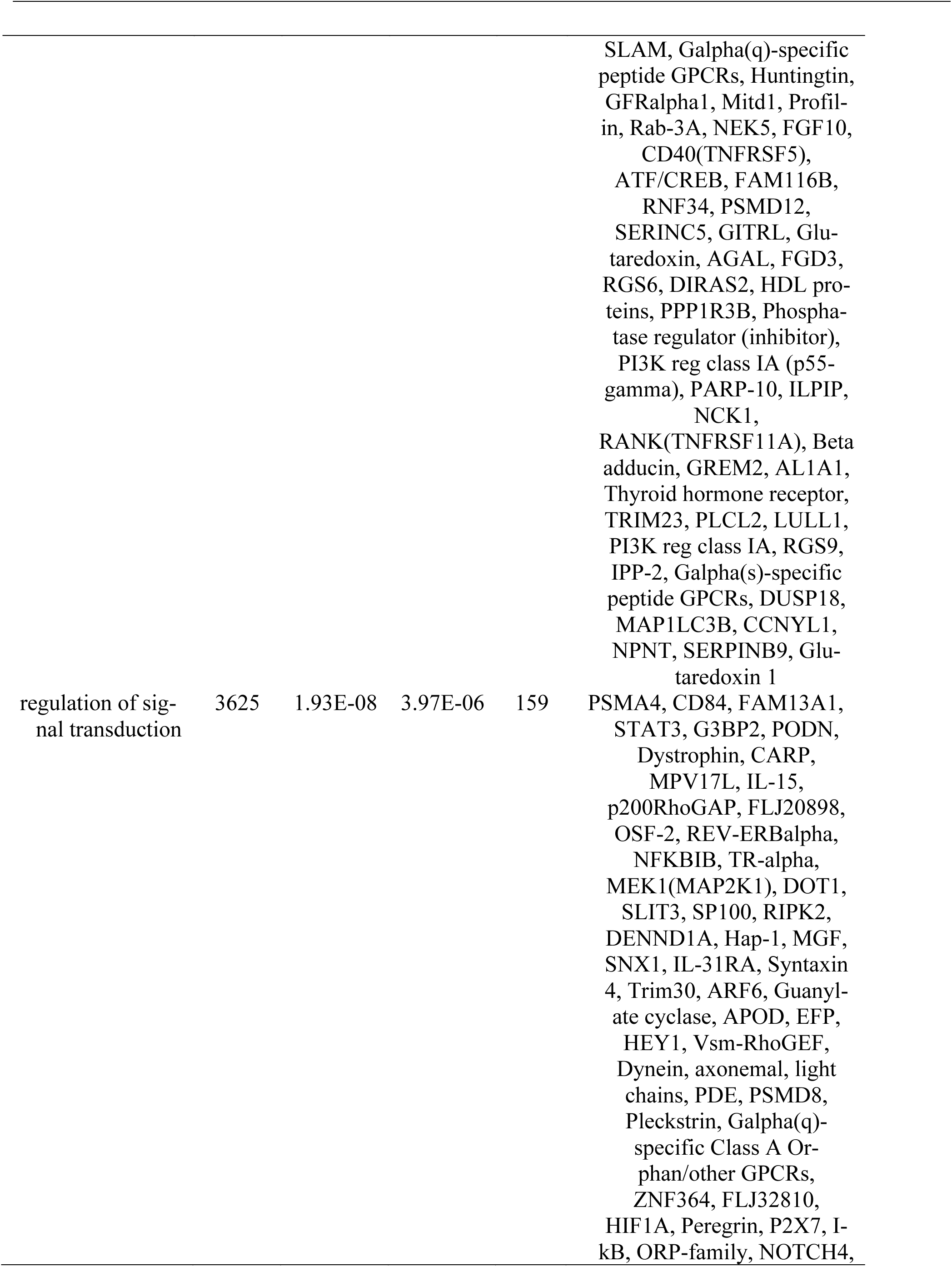
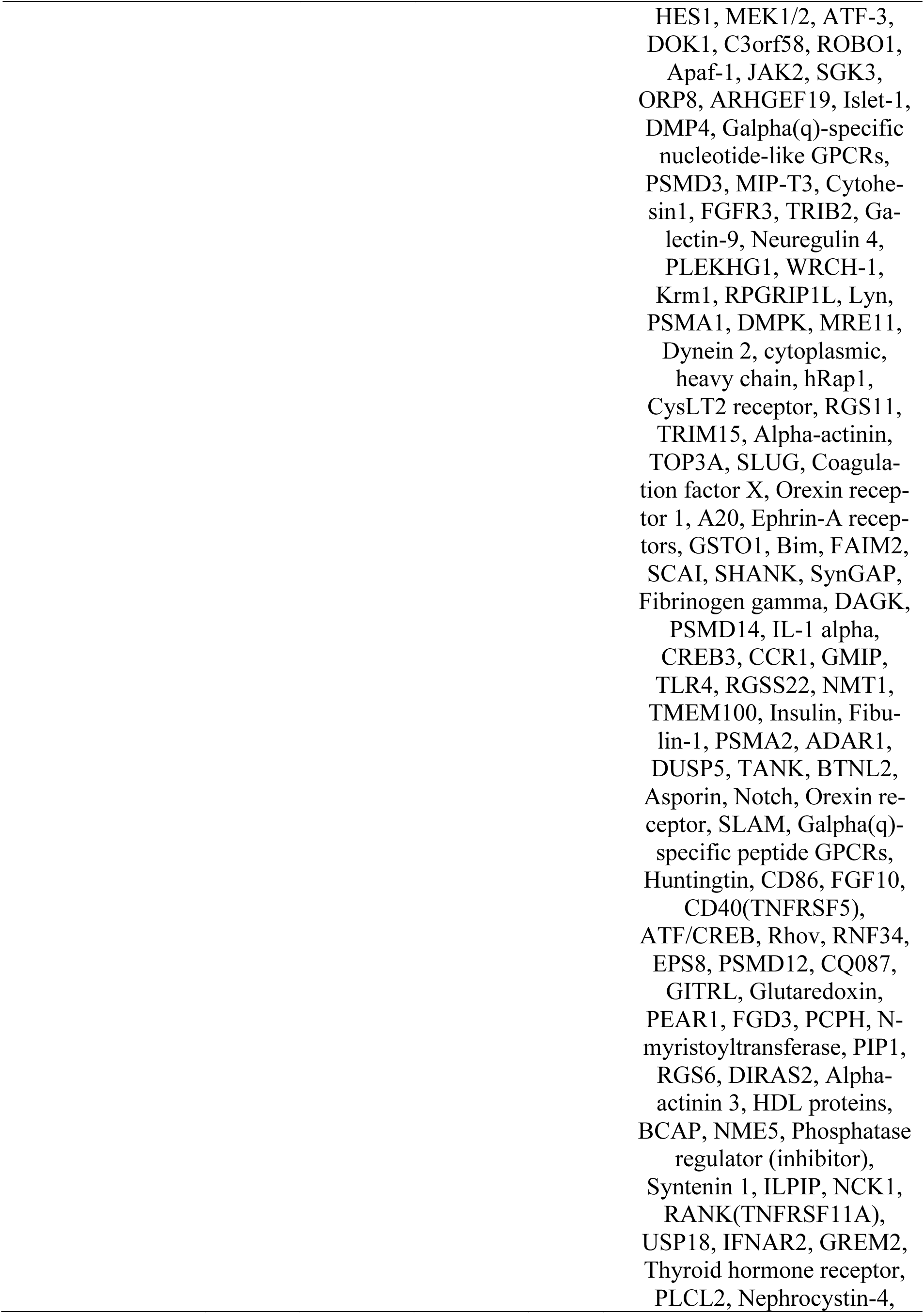
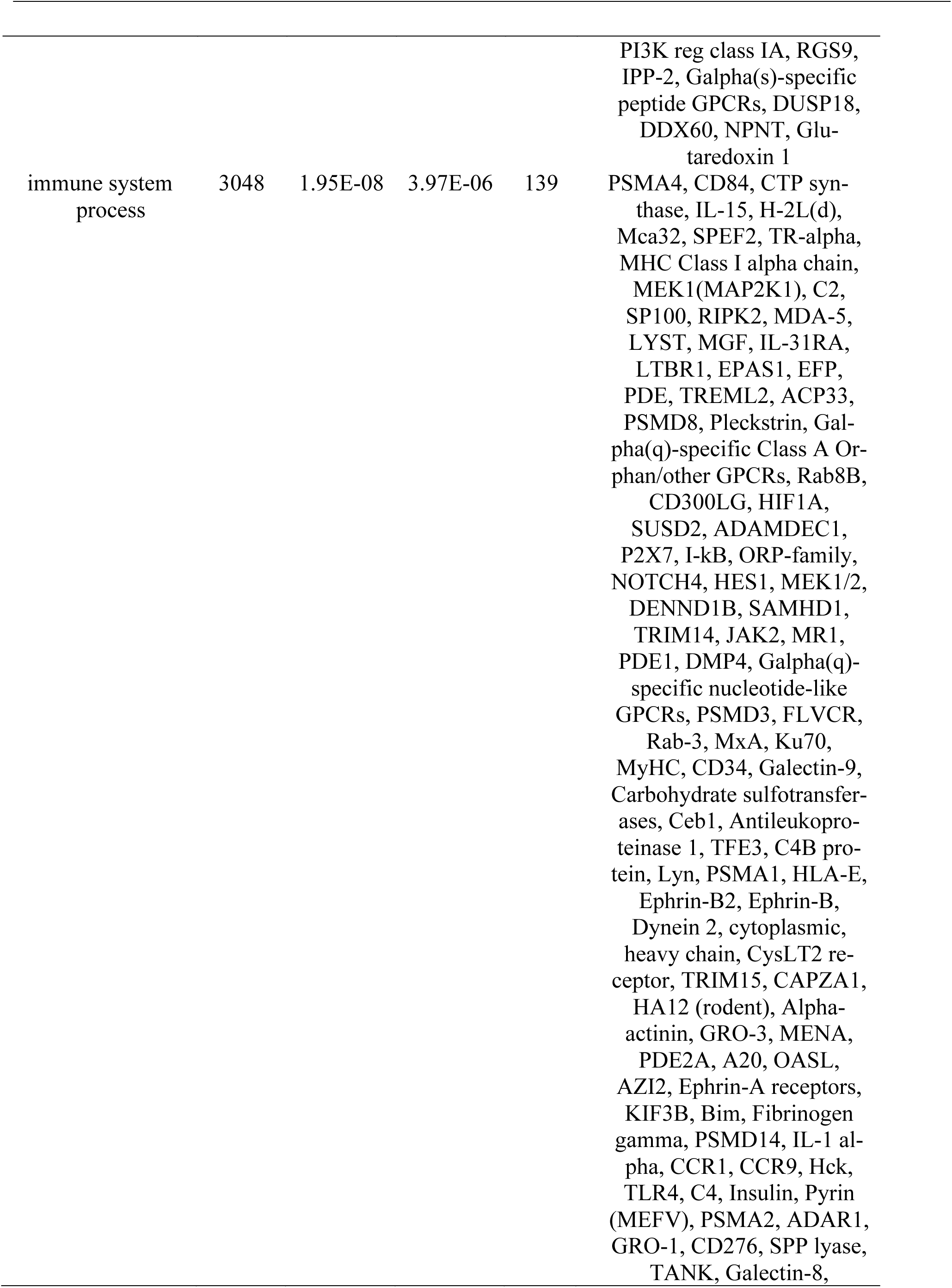
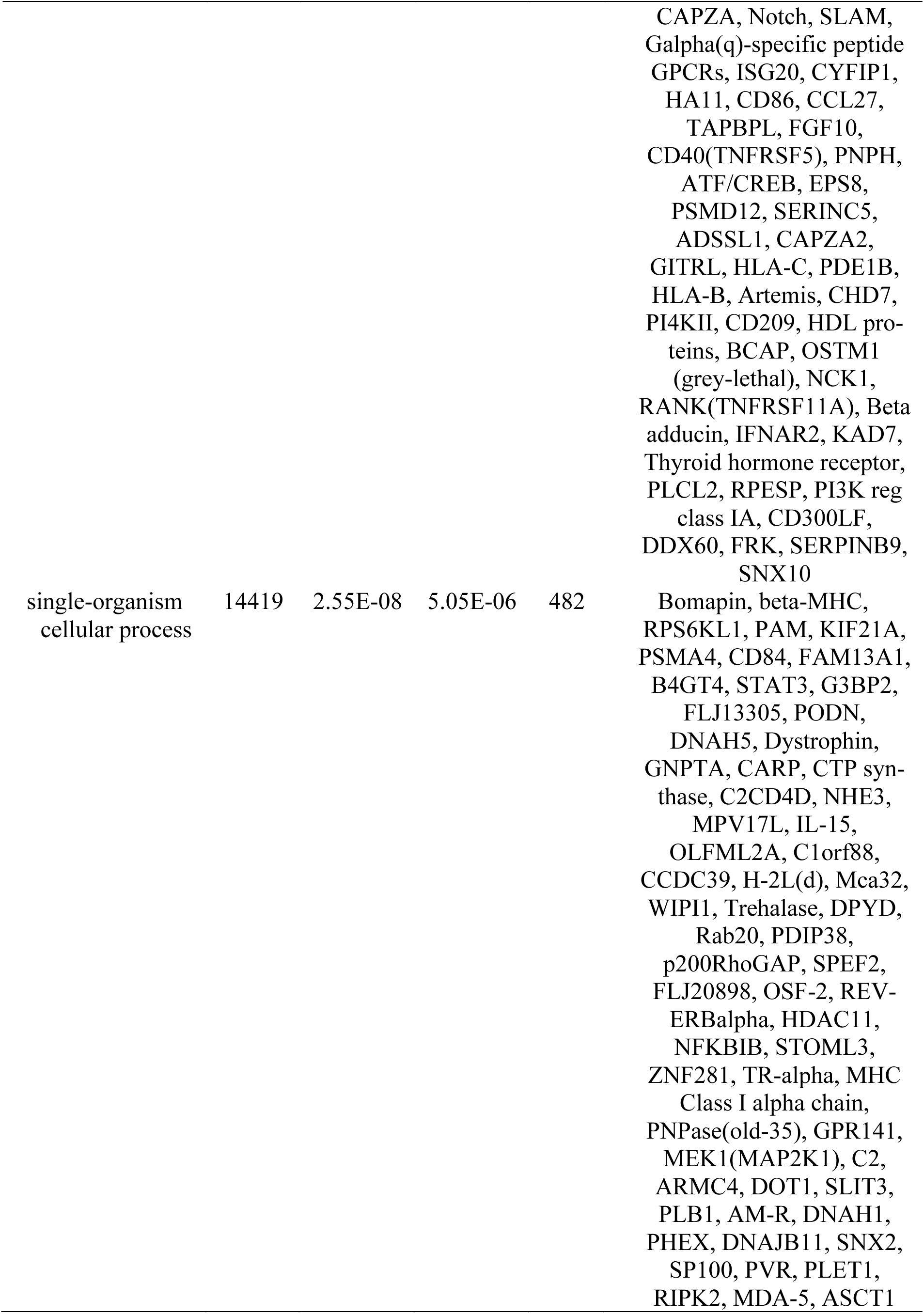
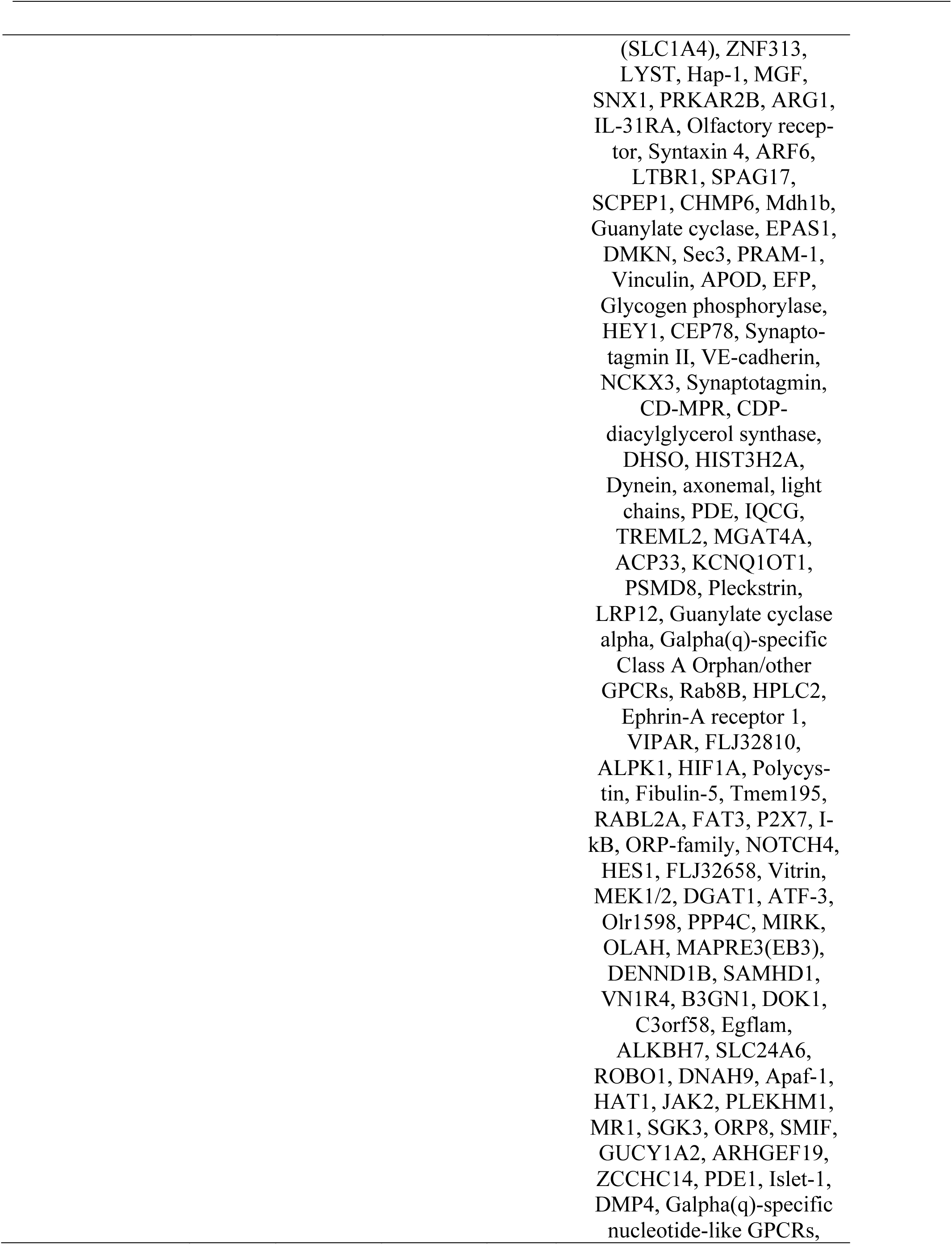
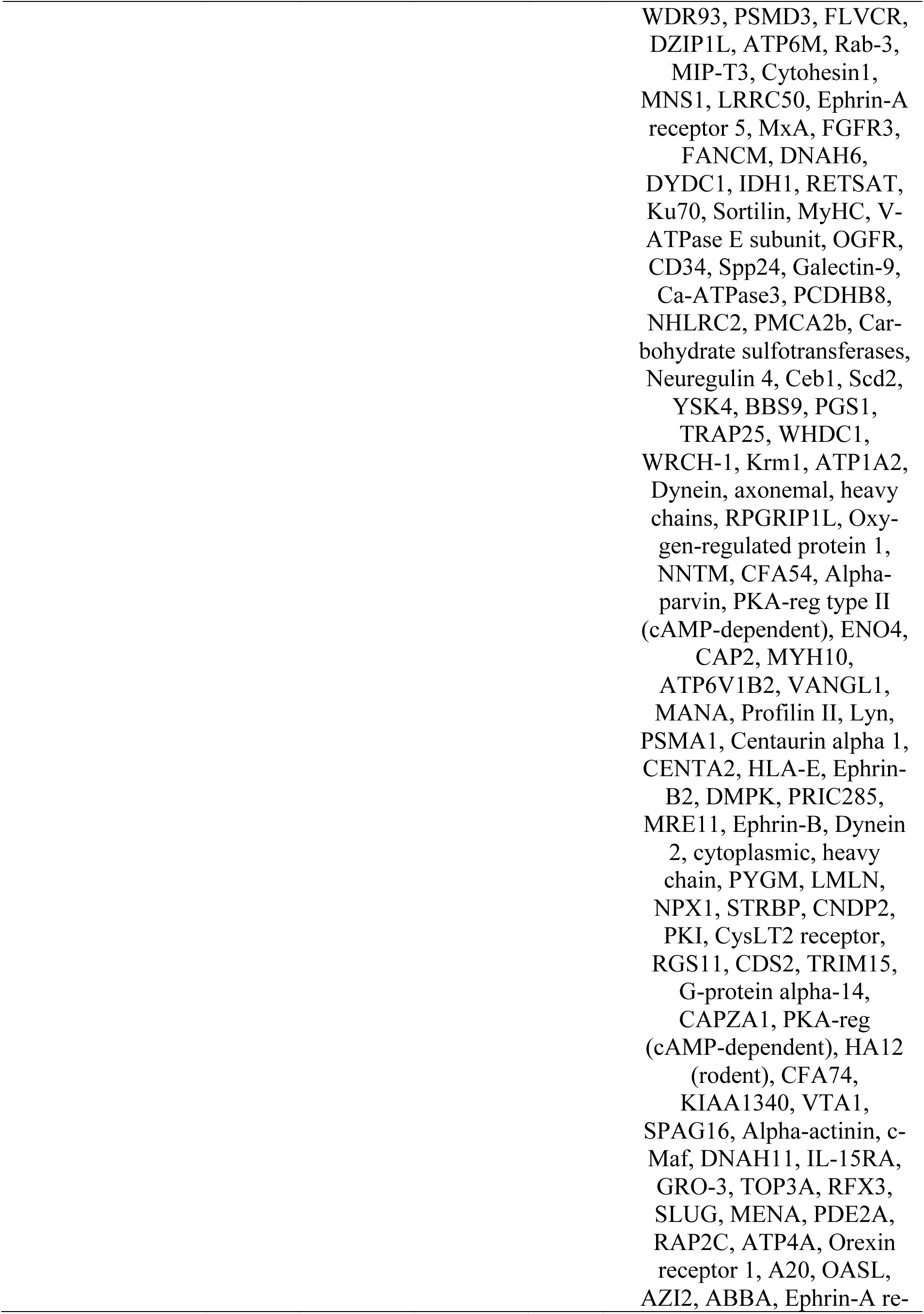
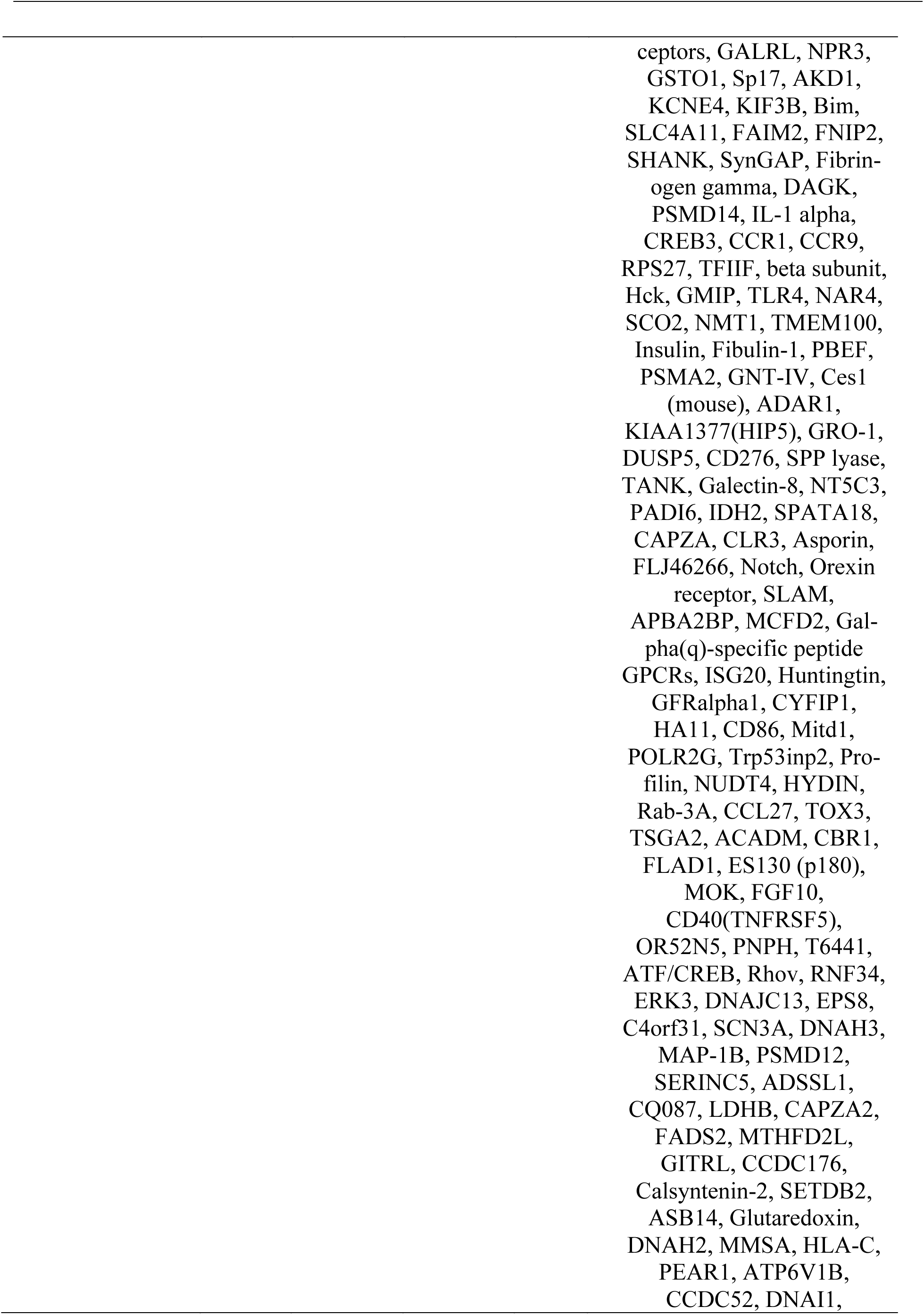
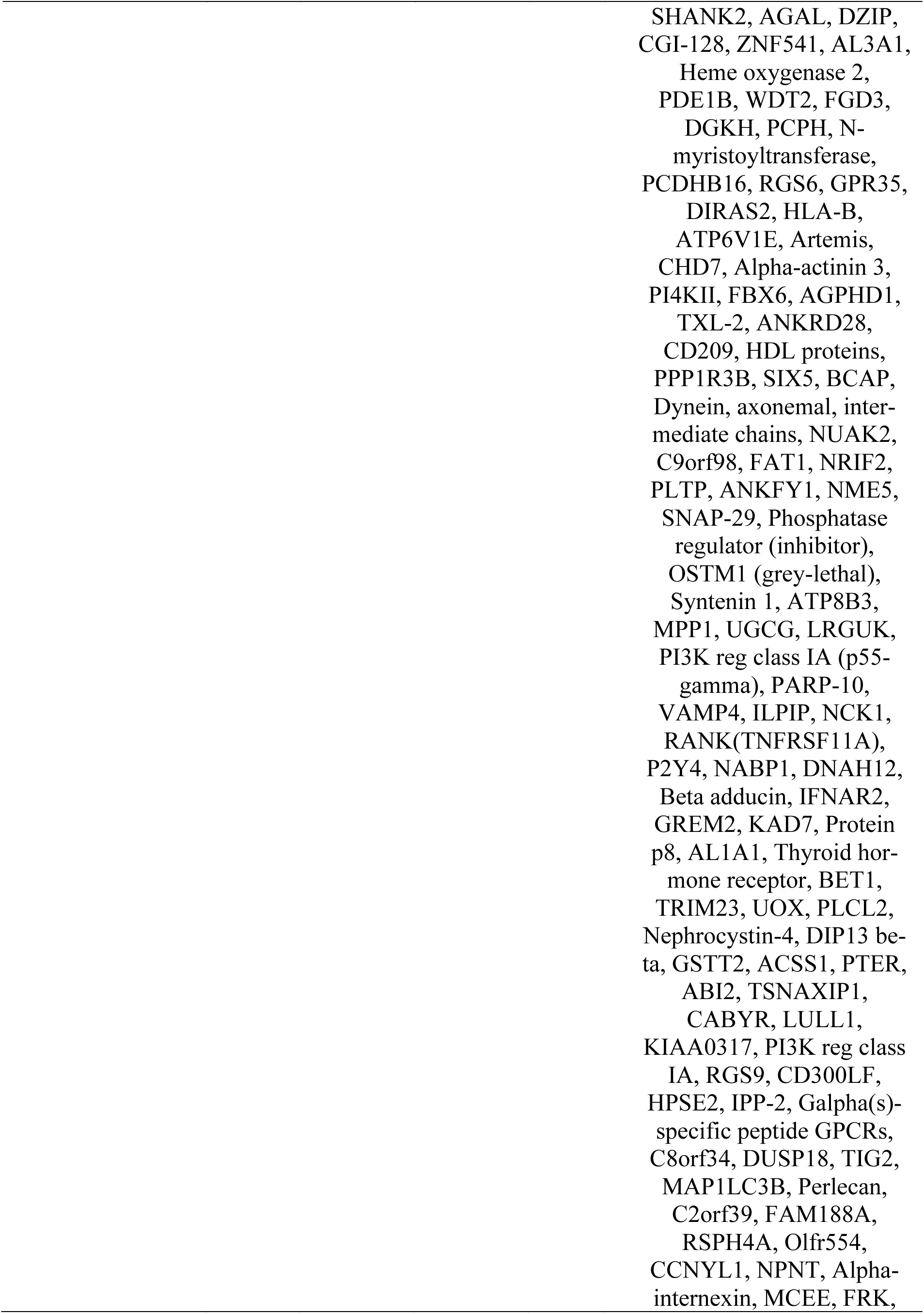
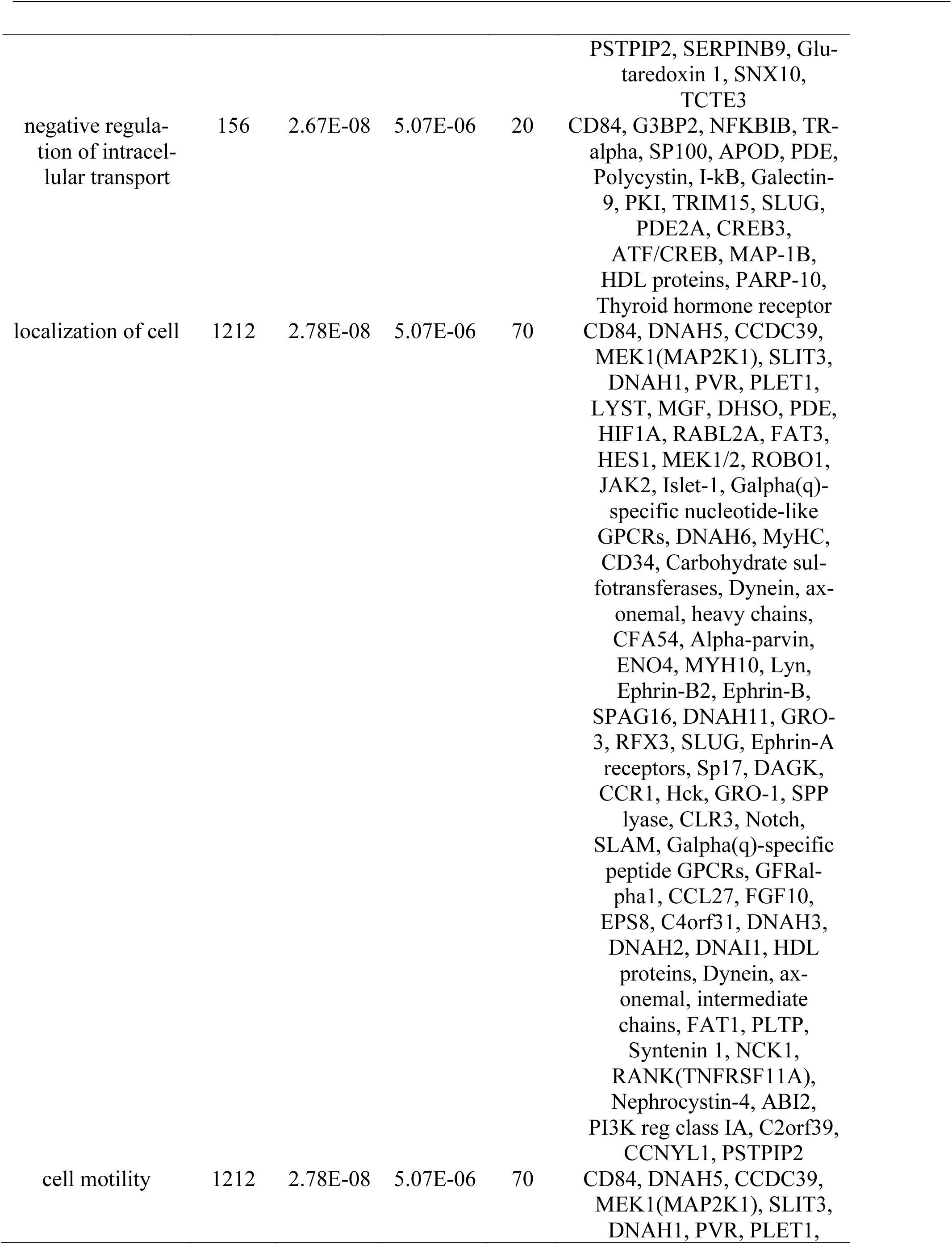
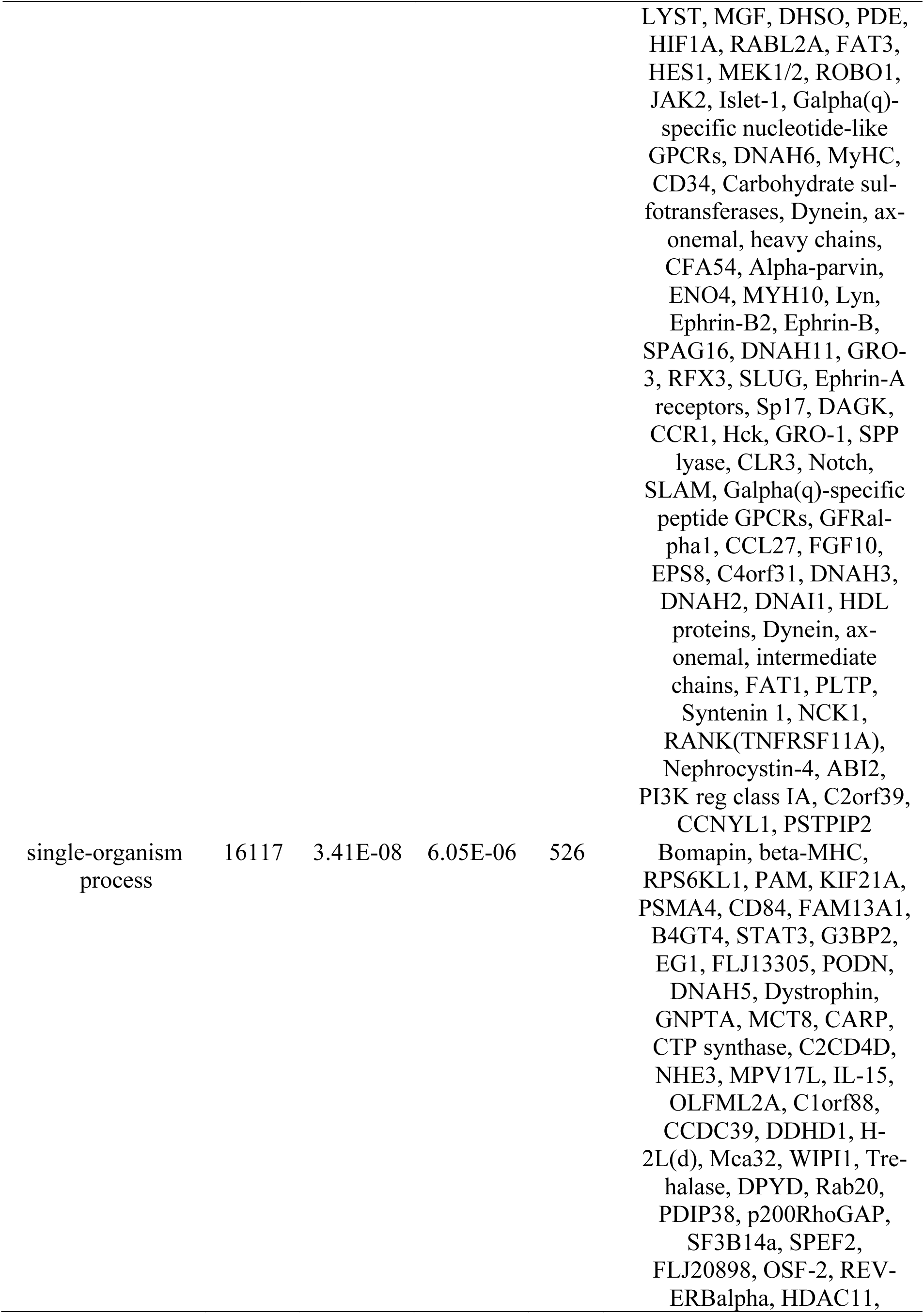
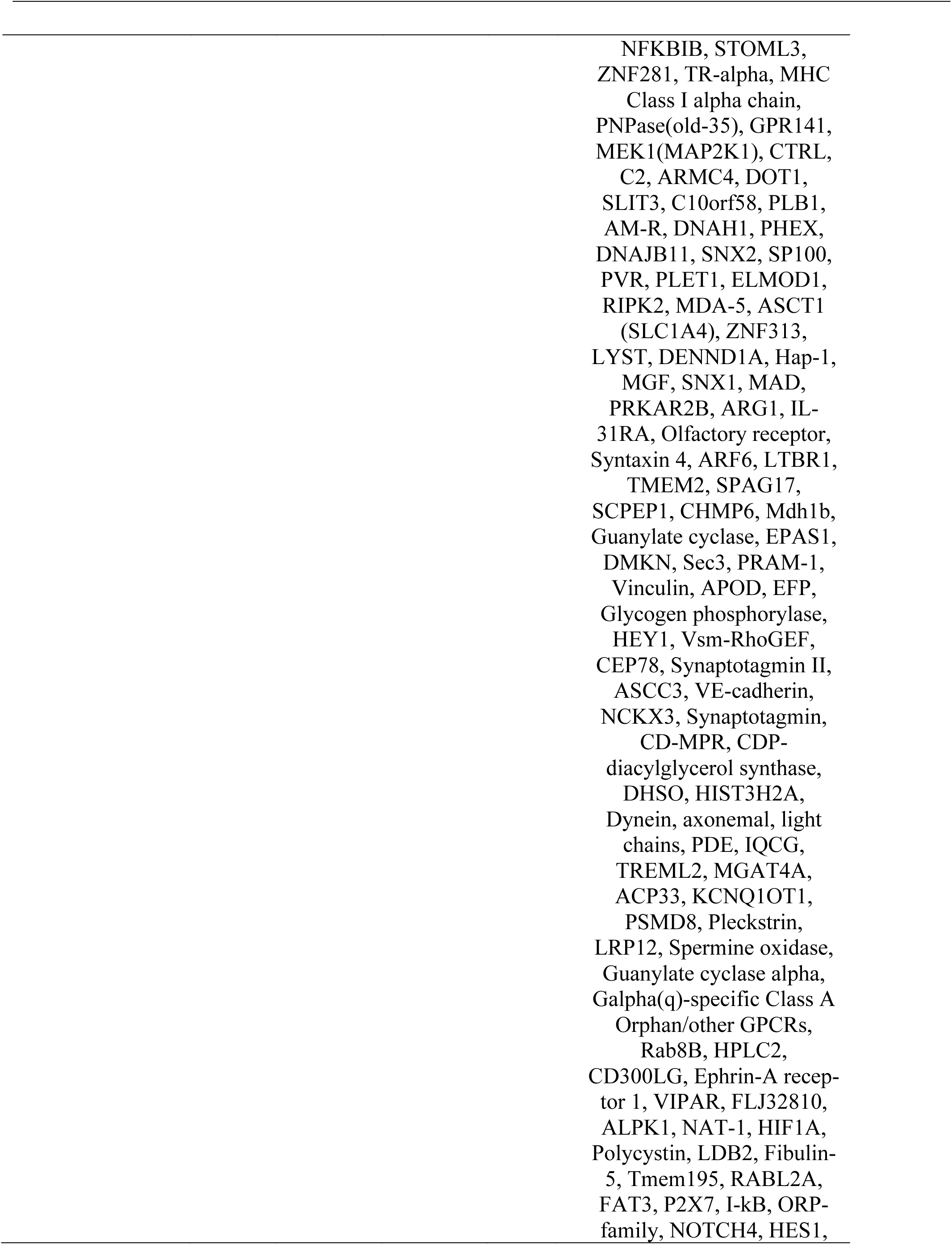
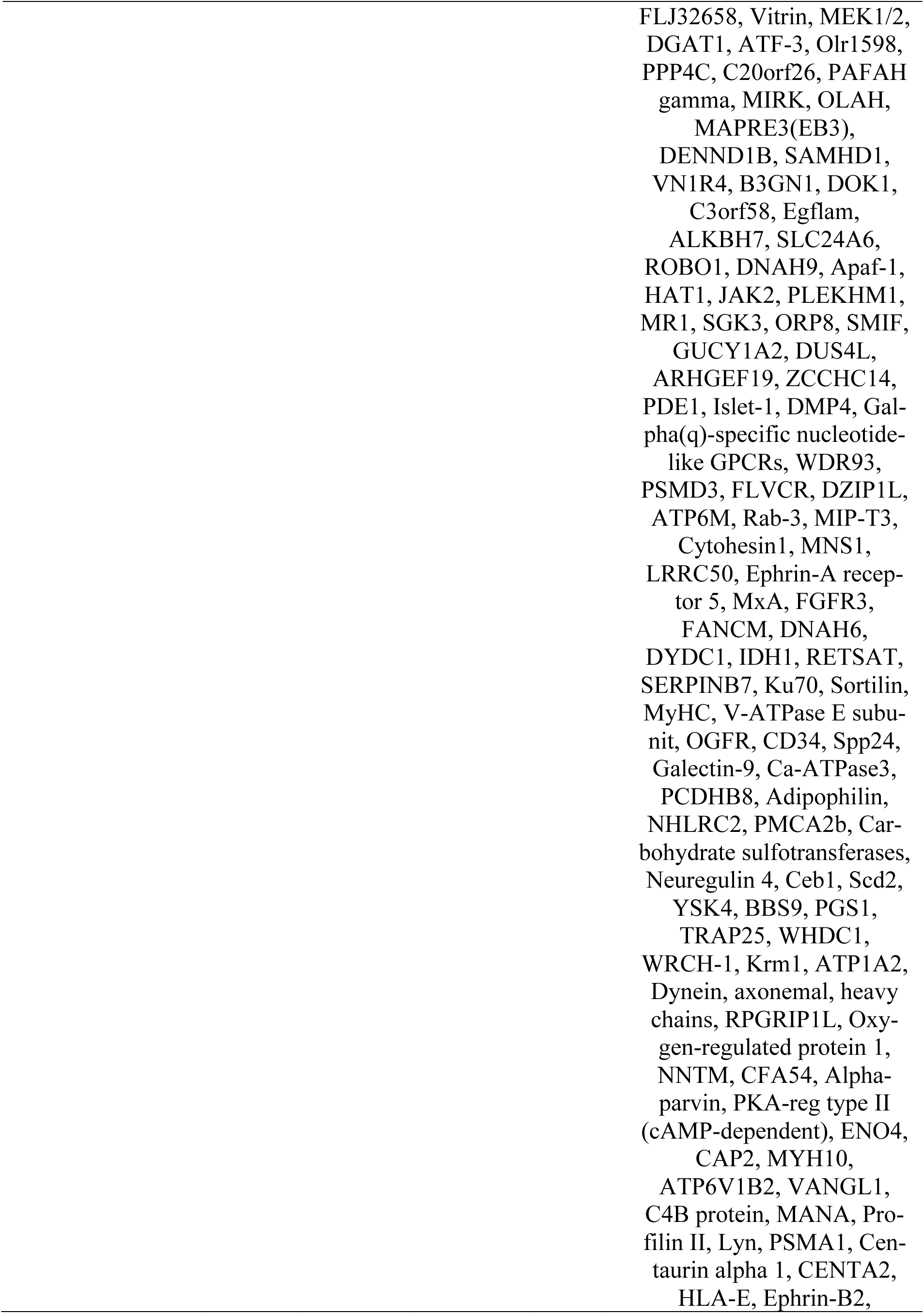
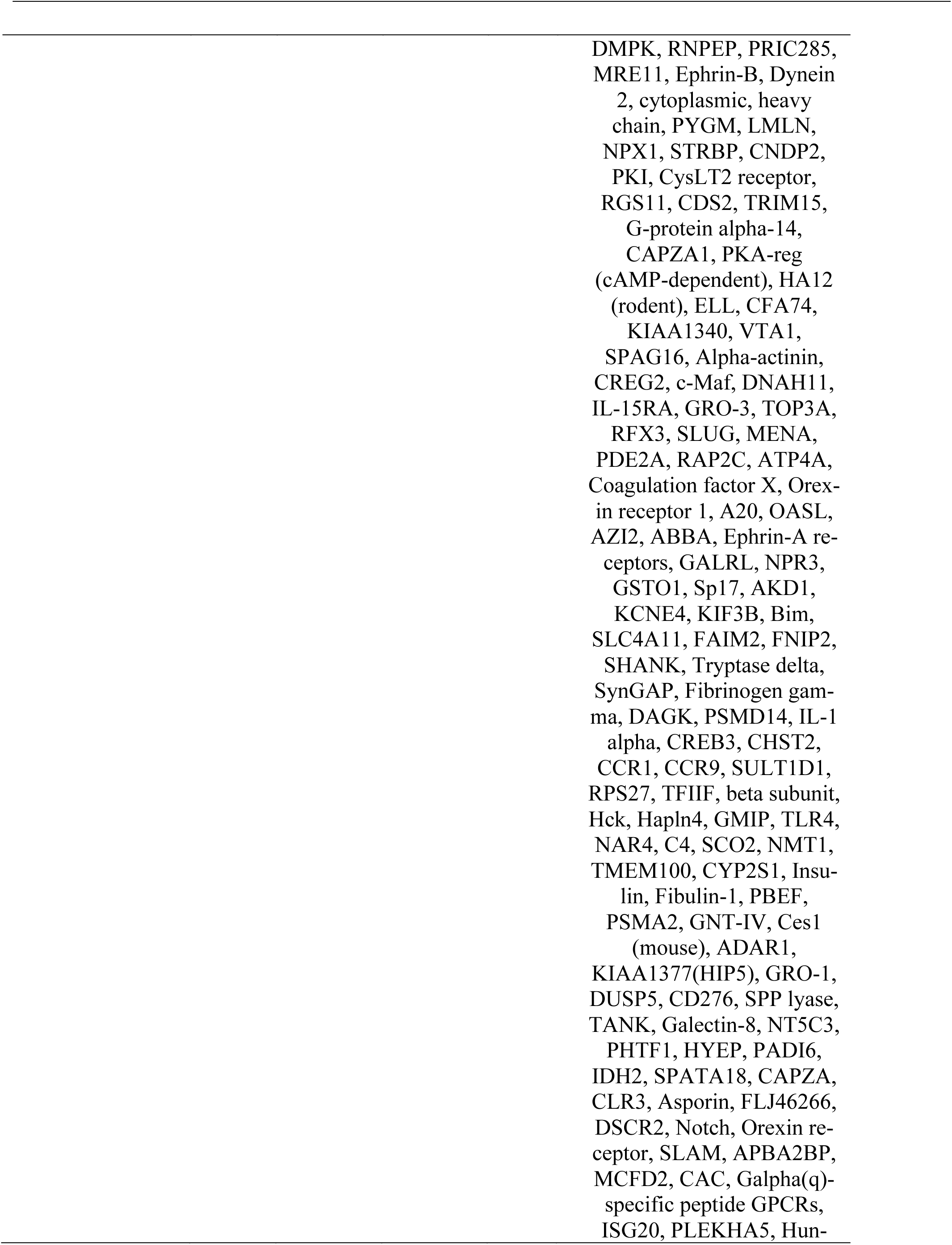
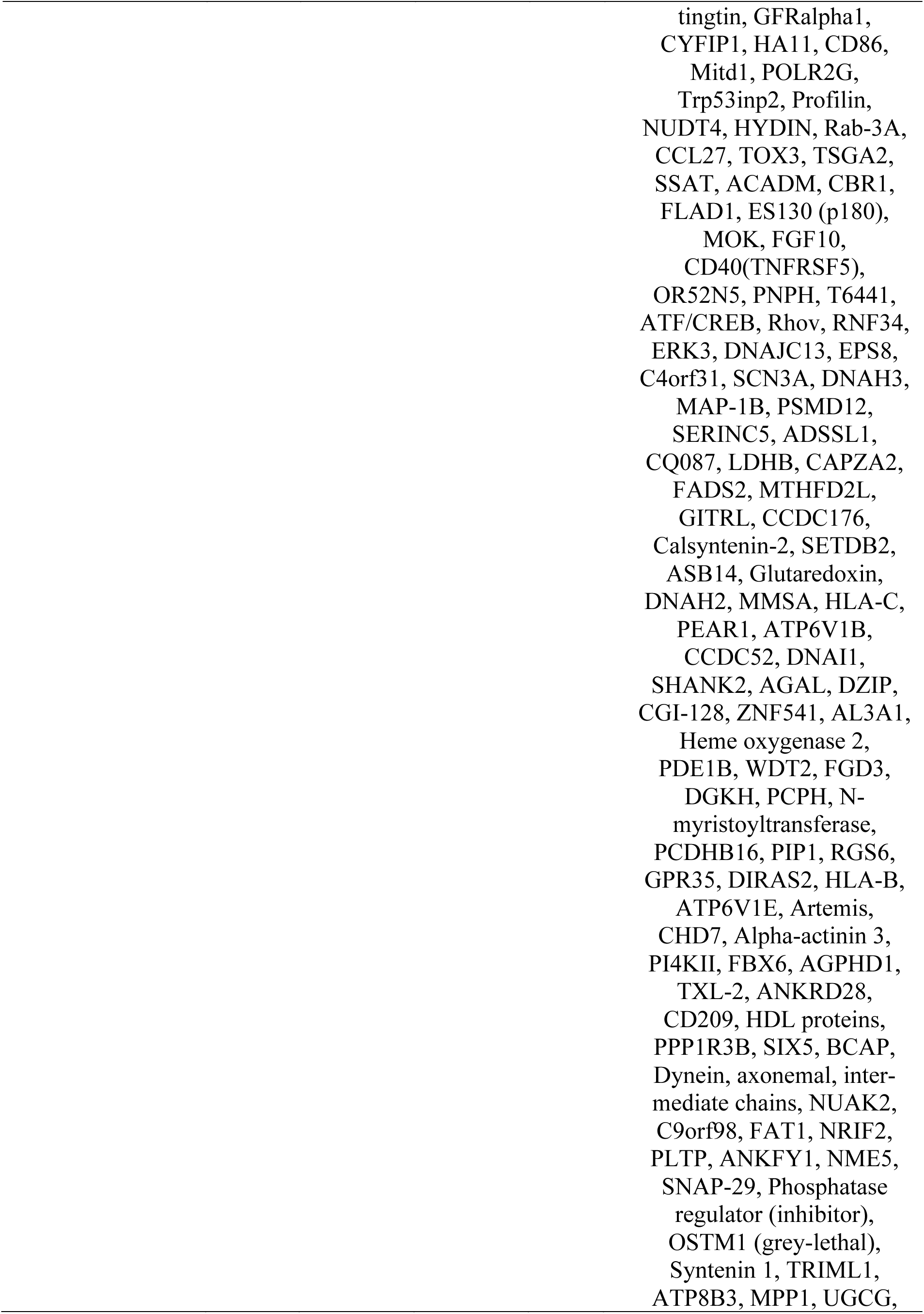
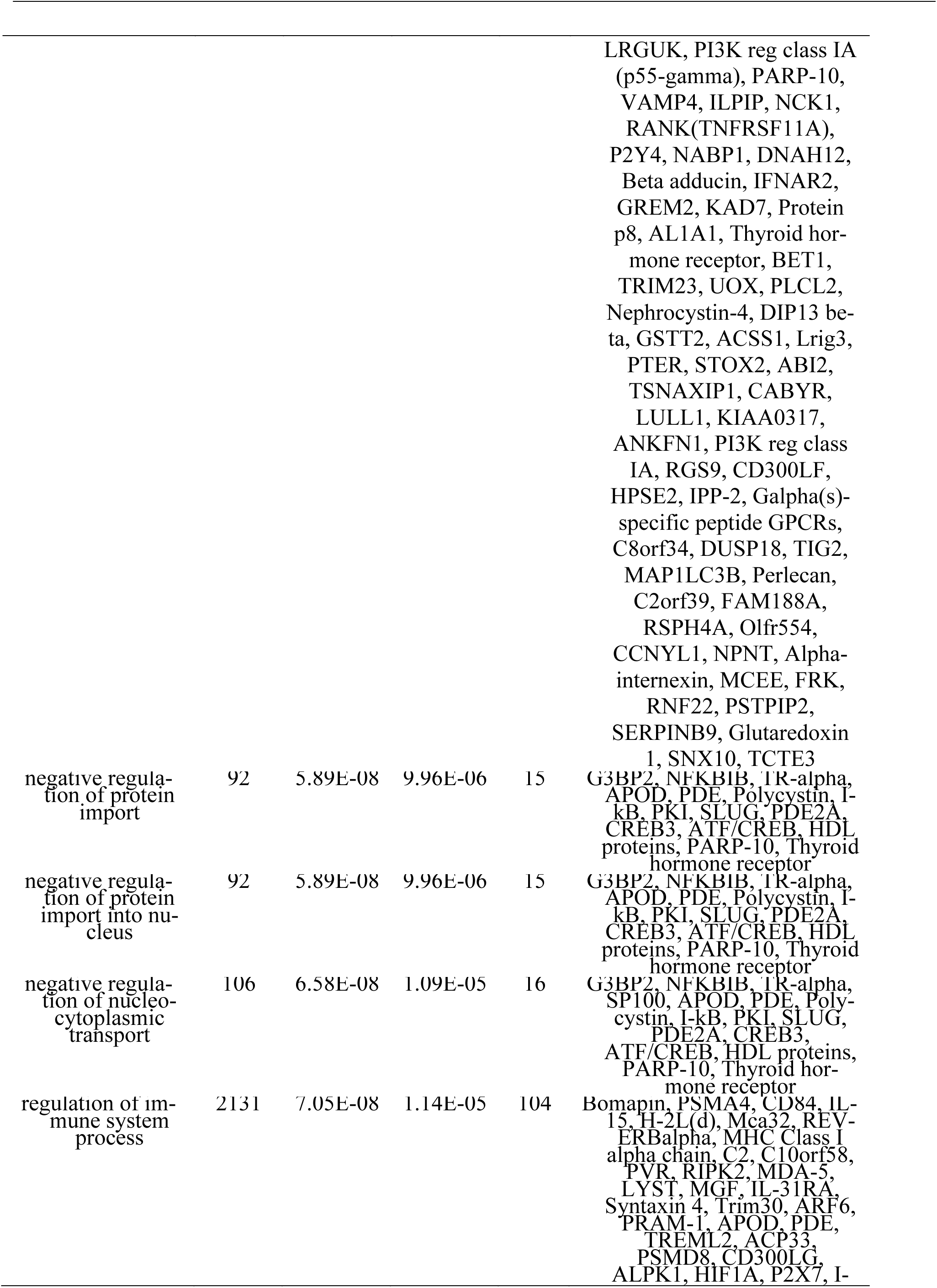
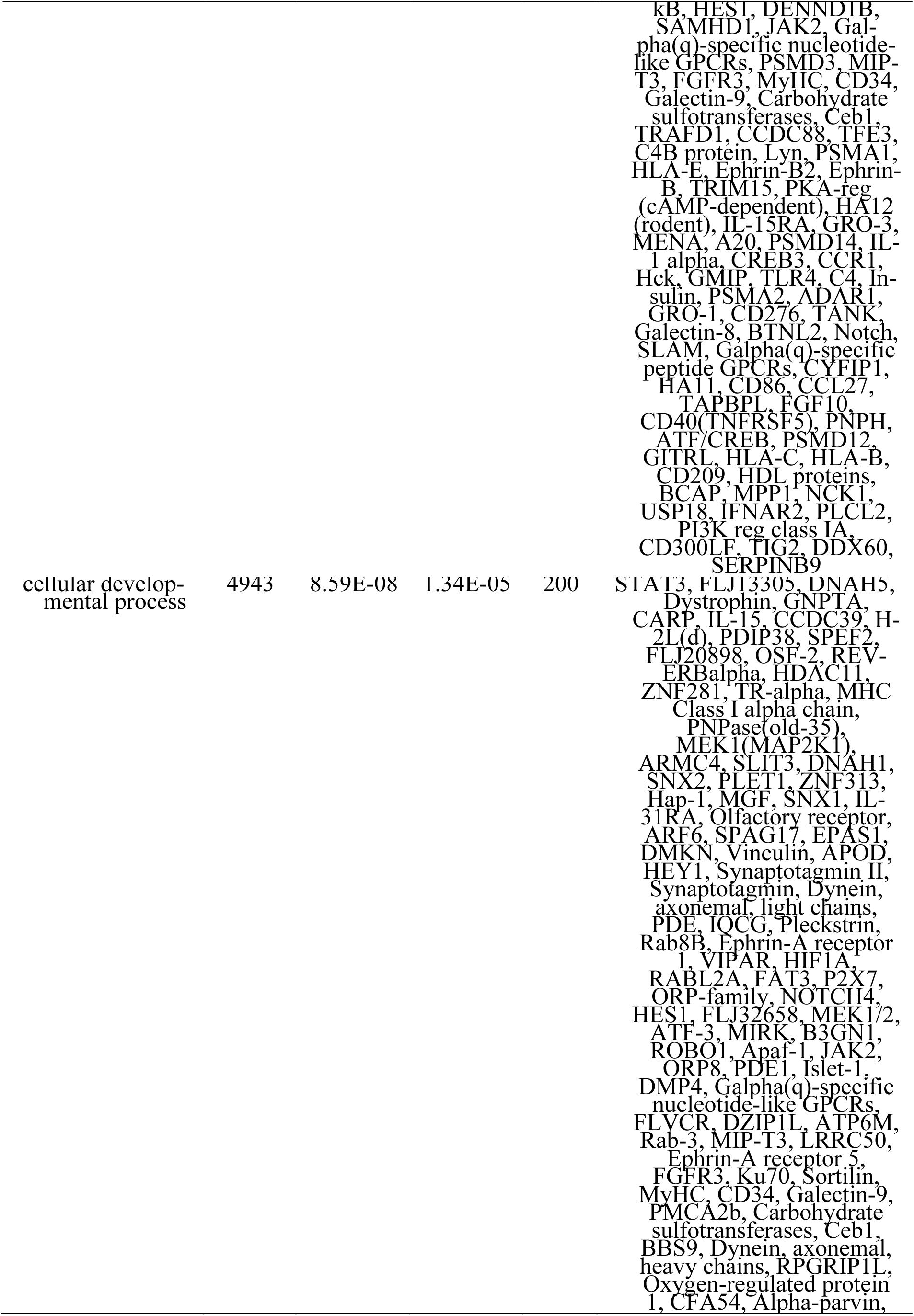
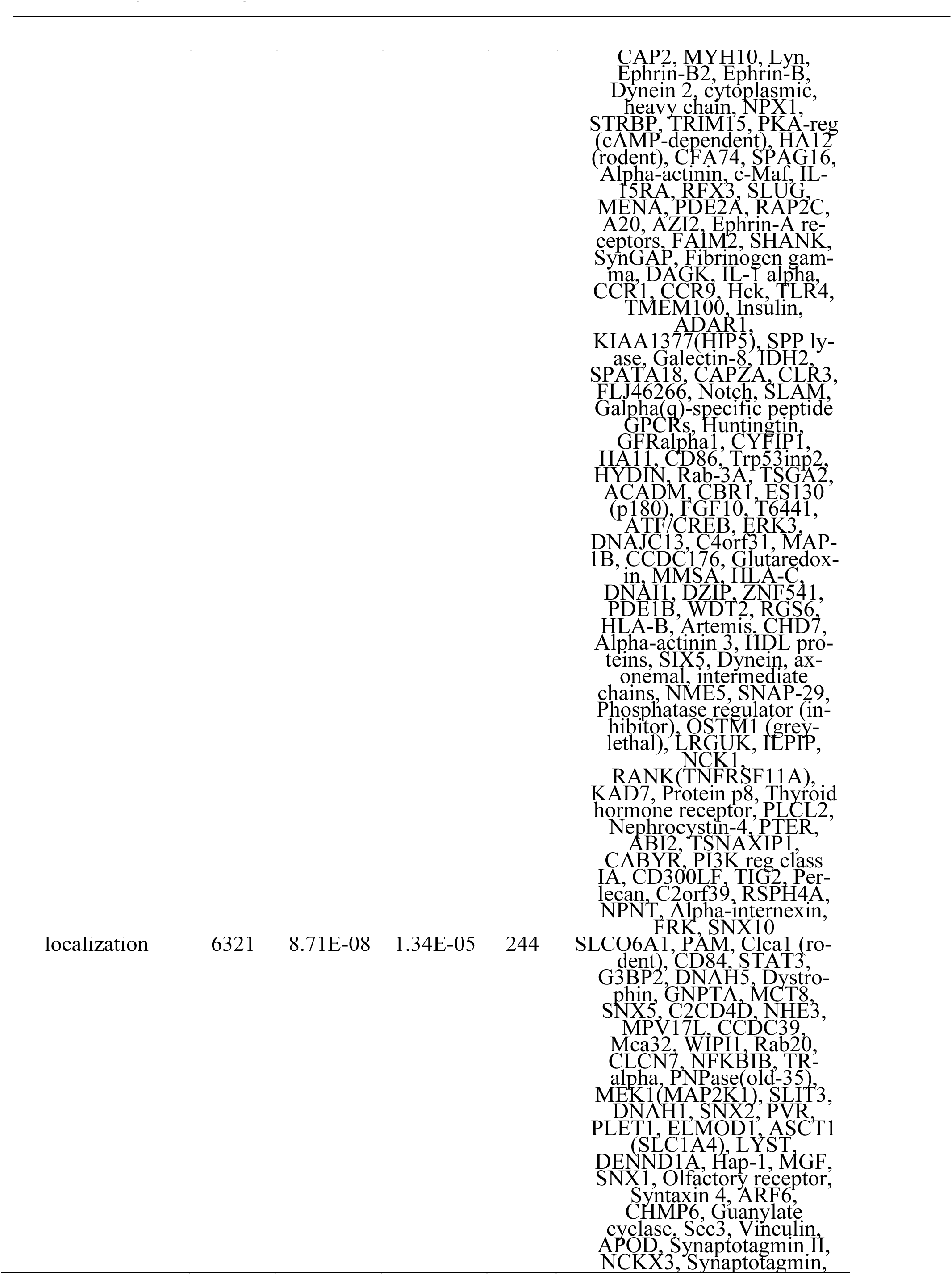
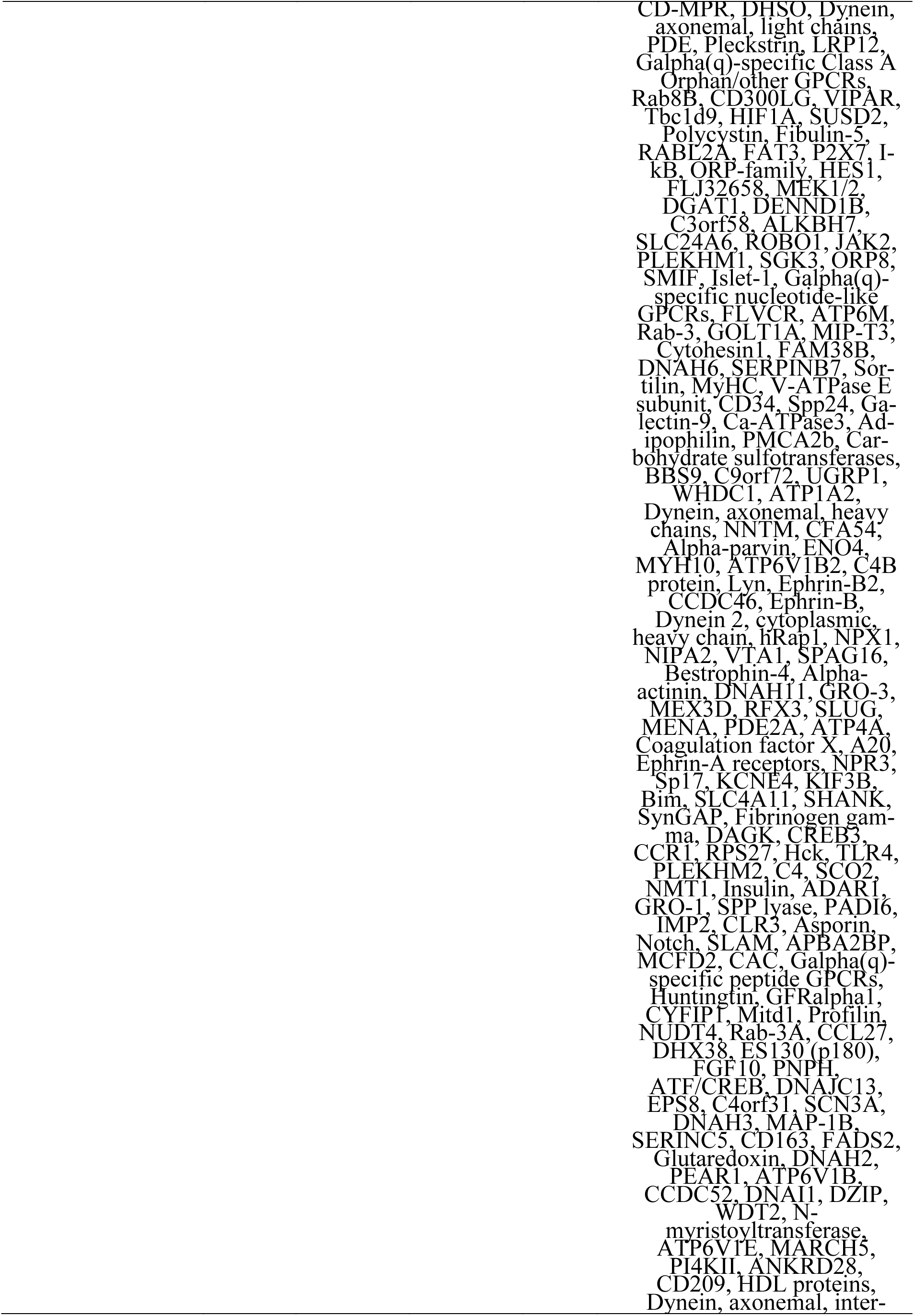
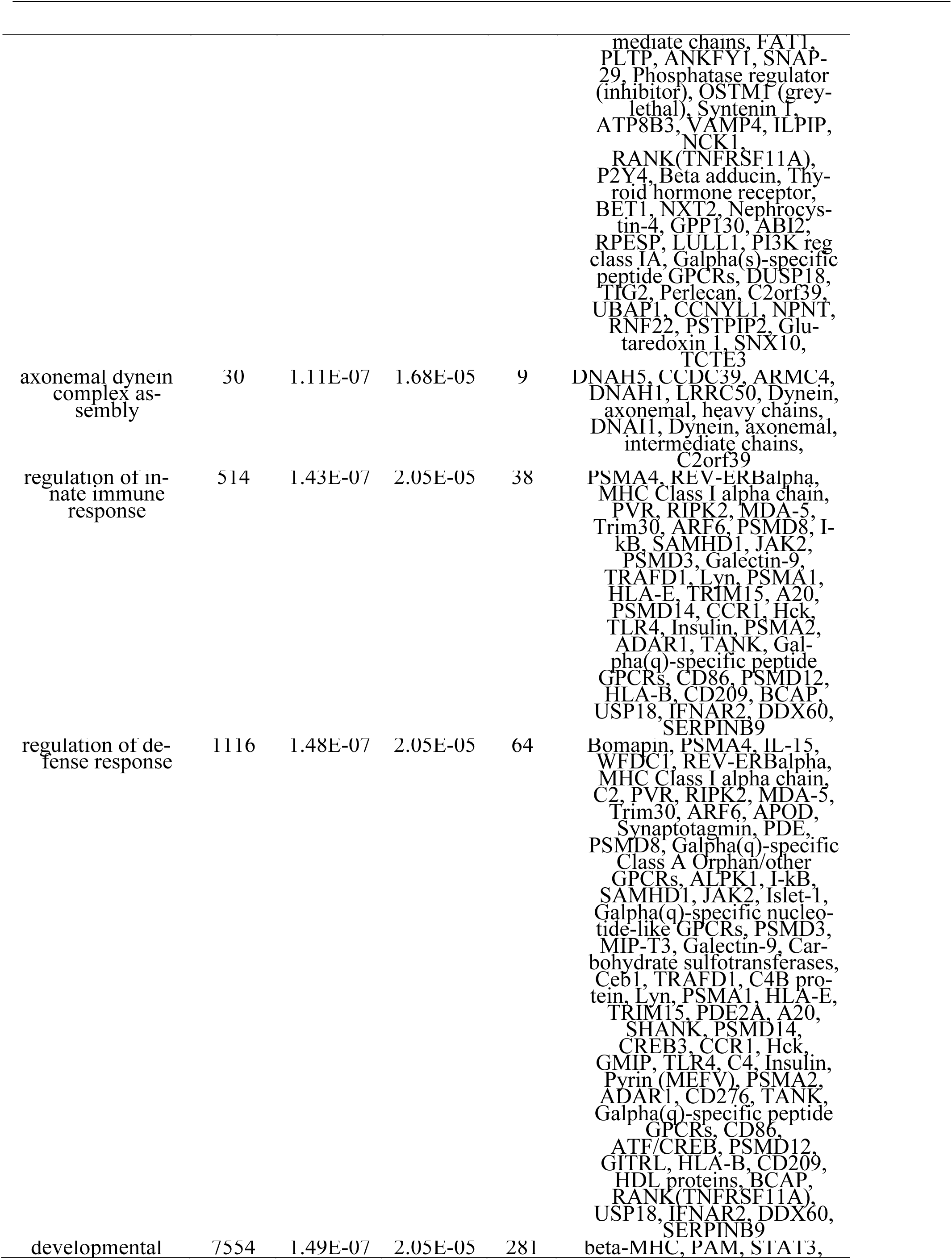
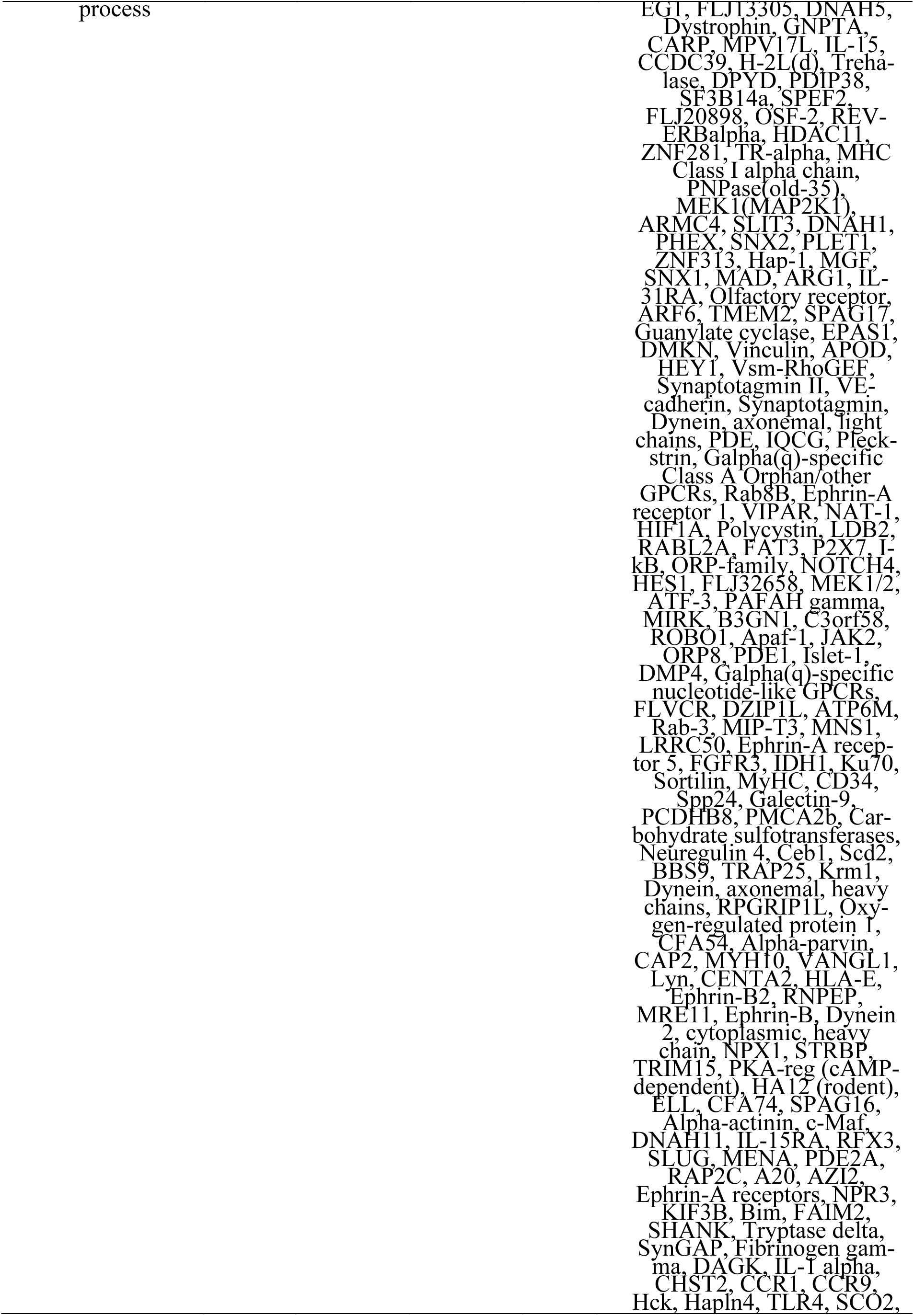
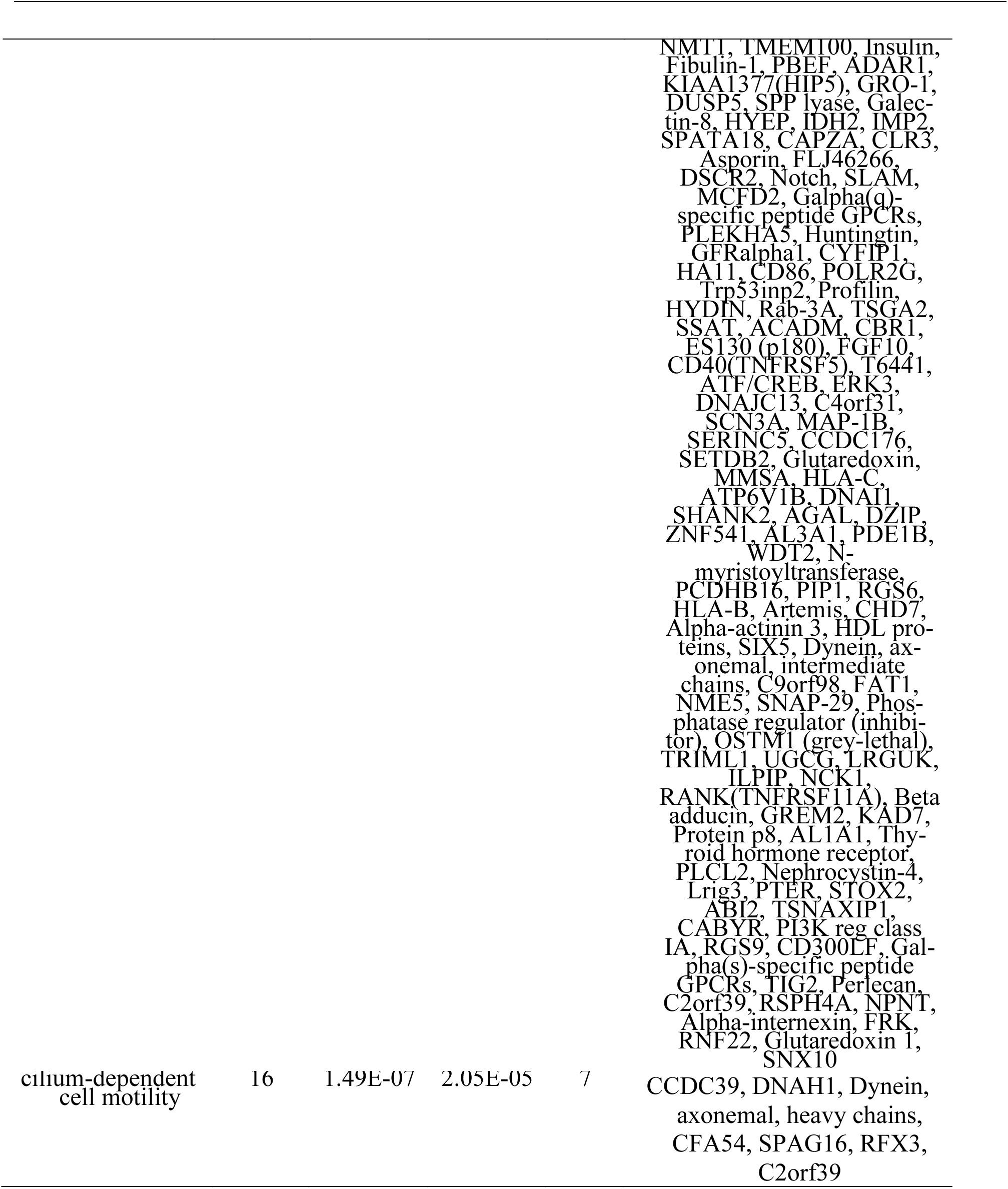
GO processes analysis of the 830 genes in cluster 2 in the H1N1 integrative network using MetaCore. Only the top 50 significantly enriched GO processes are shown here.

## References

Barabasi, A.L. et al. (2011) Network medicine: a network-based approach to human disease. Nat Rev Genet, 12, 56–68.

Blondel VD. et al. (2008) Fast unfolding of communities in large networks. Journal of Statistical Mechanics: Theory and Experiment, 2008 P10008.

Cancer Genome Atlas Research Network. (2008) Comprehensive genomic characterization defines human glioblastoma genes and core pathways. Nature, 455,1061–1068.

Chandler JD. et al. (2016) Metabolic pathways of lung inflammation revealed by high-resolution metabolomics (HRM) of H1N1 influenza virus infection in mice. Am J Physiol Regul Integr Comp Physiol,311, R906–R916.

Gonzalez, I. et al. (2012) Visualising associations between paired ‘omics’ data sets. BioData Min, 5, 19.

Hawkins, R.D. et al. (2010) Next-generation genomics: an integrative approach. Nat Rev Genet, 11,476–486.

Le Cao, K.A. et al. (2009) integrOmics: an R package to unravel relationships between two omics datasets. Bioinformatics, 25, 2855–2856.

Lichtblau, Y. et al. (2016) Comparative assessment of differential network analysis methods. Brief Bioinform 2016.

Liquet, B. et al. (2012) A novel approach for biomarker selection and the integration of repeated measures experiments from two assays. BMC Bioinformatics, 13, 325.

Meng C. et al. (2016) Dimension reduction techniques for the integrative analysis of multi-omics data. Brief Bioinform, 17, 628–41.

Newman, M.E. (2006) Modularity and community structure in networks. Proc Natl Acad Sci U S A 103, 8577–8582.

Odibat, O. and Reddy, C.K. (2012) Ranking differential hubs in gene co-expression networks. J Bioinform Comput Biol, 10, 1240002.

Uppal, K. et al. (2016) Computational Metabolomics: A Framework for the Million Metabolome. Chem Res Toxicol, 29, 1956–1975.

Yang, Z. et al. (2016) A Comparative Analysis of Community Detection Algorithms on Artificial Networks. Sci Rep, 6,30750.

Wanichthanarak K. et al. (2015) Genomic, Proteomic, and Metabolomic Data Integration Strategies. Biomark Insights,10, 1–6.

## References for Supplementary Material

Chandler, J.D., et al. (2016) Metabolic pathways of lung inflammation revealed by high-resolution metabolomics (HRM) of H1N1 influenza virus infection in mice. Am J Physiol Regul Integr Comp Physiol, 311, R906–R916.

Fortin, G., et al. (2009) L-carnitine, a diet component and organic cation transporter OCTN ligand, displays immunosuppressive properties and abrogates intestinal inflammation. Clin Exp Immunol, 156, 161–171.

Kohlmeier, J.E. and Woodland, D.L. (2009) Immunity to respiratory viruses. Annu Rev Immunol, 27, 61–82.

Liu, Q. et al. (2016) The cytokine storm of severe influenza and development of immunomodulatory therapy. Cell Mol Immunol, 13, 3–10.

Miyake, J.H. et al. (2000) Bile acid induction of cytokine expression by macrophages correlates with repression of hepatic cholesterol 7alpha-hydroxylase. J Biol Chem, 275, 21805–21808.

Sadik, C.D. and Luster, A.D. (2012) Lipid-cytokine-chemokine cascades orchestrate leukocyte recruitment in inflammation. J Leukoc Biol, 91, 207–215.

Shannon P. et al. (2003) Cytoscape: a software environment for integrated models of biomolecular interaction networks. Genome Res, 13, 2498–2504.

Yin, K. and Agrawal, D.K. Vitamin D and inflammatory diseases. (2014) J Inflamm Res 7, 69–87.

